# A common pathway controls cell size in the sepal and leaf epidermis leading to a non-random pattern of giant cells

**DOI:** 10.1101/2024.07.26.605215

**Authors:** Frances K. Clark, Gauthier Weissbart, Xihang Wang, Kate Harline, Chun-Biu Li, Pau Formosa-Jordan, Adrienne H. K. Roeder

**Affiliations:** Department of Molecular Biology and Genetics, Cornell University, Ithaca, NY 14853; Weill Institute for Cell and Molecular Biology and School of Integrative Plant Science, Section of Plant Biology, Cornell University, Ithaca, NY 14853; Department of Plant Developmental Biology, Max Planck Institute for Plant Breeding Research, Cologne, Germany; Department of Mathematics, Stockholm University, 106 91 Stockholm, Sweden; Cluster of Excellence on Plant Sciences (CEPLAS), Max Planck Institute for Plant Breeding Research, Cologne, Germany; Polyploidy Integration and Innovation Institute

**Author notes:** Authors for correspondence: (PFJ), (AHKR). Authors contributed equally to this work (Frances K. Clark, Gauthier Weissbart, Xihang Wang).

## Abstract

Arabidopsis leaf epidermal cells have a wide range of sizes and ploidies, but the mechanisms patterning their size and spatial distribution remain unclear. Here, we show that the same genetic pathway creating giant cells in sepals also regulates cell size in the leaf epidermis, leading to the formation of giant cells. In both sepals and leaves, giant cells are scattered among smaller cells; therefore, we asked whether their spatial arrangement is random. By comparing sepal and leaf epidermises with computationally generated randomized tissues we show that the giant cell pattern becomes less random across the epidermis as the cells surrounding giant cells divide, leading to clustered patterns in mature tissues. Our cell-autonomous and stochastic computational model reproduces the giant cell organization, suggesting that random giant cell initiation together with the divisions of surrounding cells lead to the observed clustered pattern. These findings reveal that cell-size patterning is developmentally regulated by common mechanisms in leaves and sepals, and the spatial pattern of giant cells emerges from the interplay between stochastic cell-autonomous gene expression and tissue growth.

## Introduction

The *Arabidopsis thaliana* (hereafter Arabidopsis) mature leaf blade epidermis contains three main cell types: stomatal guard cells, trichomes, and pavement cells (1). Stomatal guard cells surround stomatal pores through which gas exchange occurs, and trichomes are large branched hair cells that serve to discourage herbivory, among other functions (2). All other epidermal cells in the mature leaf blade epidermis (the expanded part of the leaf between the midrib and the margin) are classified as pavement cells. However, pavement cells are not a homogeneous group of cells, but rather exhibit a variety of sizes, ploidies, and shapes (3,4). Much research has focused on the patterning of stomata (5–7) and trichomes (8,9), leading to important insights into how the regulation of intercellular signaling, cell fate specification, the cell cycle, and polarized cell division orientation give rise to their spatial arrangement. However, the patterning of pavement cells is understudied. In particular, little is known about how some pavement cells are specified to become larger and more highly polyploid than others.

Pavement cell-size patterning has been studied in the Arabidopsis sepal. Pavement cells in the sepal vary in size and ploidy, with some cells reaching up to 800 μm in length (Fig. 1A) and having ploidies up to 32C (10). These very large pavement cells that have a characteristic highly anisotropic shape and bulge out of the epidermis have been named ‘giant cells’ (10), and these form when a cell endoreduplicates early during growth (10). Endoreduplicating cells replicate their DNA but do not enter mitosis or divide and instead continue to grow and increase their ploidy. Once a cell enters endoreduplication, it is thought to terminally differentiate and almost never re-enters the mitotic cycle (10). Similar numbers of giant cells form on sepals within an Arabidopsis plant and among plants, but the precise spatial arrangement of giant cells differs from sepal to sepal.

**Fig. 1.**
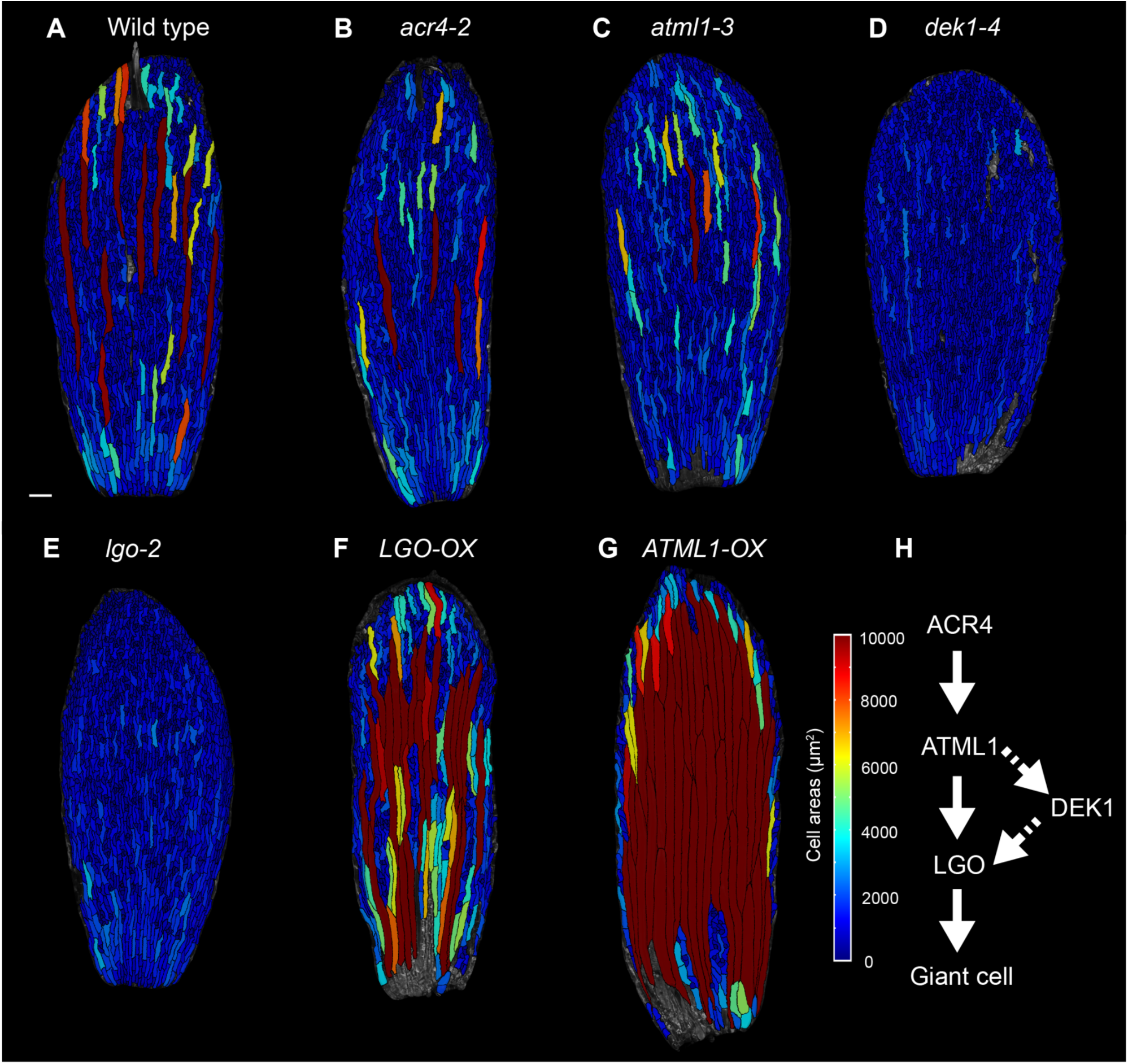
The genetic pathway that regulates giant cell development in sepals. (**A**–**G**) Cell area heat maps in μm^2^ of the abaxial (outer) surface of a stage 14 adult sepal of (**A**) wild type, (**B**) *acr4- 2*, (**C**) *atml1-3*, (**D**) *dek1-4*, (**E**) *lgo-2*, (**F**) *LGO-OX* (*pATML1::LGO*) and (**G**) *ATML1-OX* (*pPDF1::FLAG-ATML1*). Scale bar represents 100 µm. (**H**) The ordering of genes into a genetic pathway according to double-mutant phenotypic analysis. The underlying data for this figure can be found at Open Science Framework (<u>osf.io</u>), DOI:10.17605/OSF.IO/RFCWS.

Forward-genetic screens have identified the genes involved in sepal giant cell patterning, and double mutant analysis has allowed these genes to be ordered within a genetic pathway (10–13) (Fig. 1). The homeodomain leucine zipper (HD-ZIP) Class IV transcription factor Arabidopsis thaliana MERISTEM LAYER1 (ATML1) promotes giant cell specification in a dose-dependent manner (11,12). Loss of *ATML1* function in sepals greatly reduces giant cell number, and overexpression of *ATML1* leads to ectopic giant cell formation (Fig. 1A,C,G) (11,12). ATML1 protein concentration fluctuates in the protodermal nuclei of developing sepals (12). High concentrations of ATML1 reached during the G2 phase of the cell cycle are strongly correlated with giant cell differentiation, consistent with a model in which an ATML1 concentration that surpasses a threshold in G2 results in giant cell specification, early endoreduplication, and giant cell differentiation (12). The receptor-like kinase ARABIDOPSIS CRINKLY 4 (ACR4) functions upstream of ATML1 to promote giant cell formation (11,12,14–16) (Fig. 1B,H). Loss of function of *ACR4* leads to a modest reduction in the number of giant cells (11) (Fig. 1A, B). The calpain protease DEFECTIVE KERNEL (DEK1) and the CDK inhibitor LOSS OF GIANT CELLS FROM ORGANS (LGO; also known as SIAMESE-RELATED 1, SMR1) function genetically downstream of ATML1 to promote giant cell formation (12) (Fig. 1H). A hypomorphic mutant *dek1* allele *(dek1-4*) results in the complete loss of giant cells from sepals (11) (Fig. 1D). Similarly, sepals from plants homozygous for a loss-of-function mutation in *LGO* have no giant cells (10,11) (Fig. 1E), and overexpression of *LGO* increases giant cell number (11) (Fig. 1F). It is unknown whether this genetic pathway affects cell size only in the sepal or whether it is also a more general mechanism of epidermal cell-size patterning in other organs.

Leaf pavement cell size is affected by the family of CDK inhibitors that includes LGO, known as the SIAMESE/SIAMESE-RELATED (SIM/SMR) family (17,18). SMR proteins bind to cyclin CDK complexes and inhibit their phosphorylation of downstream targets (18). *lgo* mutants lack large pavement cells and have a reduction in endoreduplication of the leaf cells as compared with those of wild type (10,17,18). In *lgo* mutants, pavement cells that should be mature continue to divide (19). Furthermore, overexpression of the closely related paralog of *LGO*, *SIM*, results in larger and more highly endoreduplicated leaf epidermal pavement cells (17). In sepals, LGO upregulates defense response genes, including glucosinolate biosynthesis genes (20), whereas in leaves, ATML1 promotes the formation of ER bodies, which contain components of the glucosinolate system, in large pavement cells (21), suggesting a common role of large cells in defense response. Whether the same upstream components of the sepal giant cell pathway also function in leaf cell-size patterning has not been thoroughly investigated. One study did compare pavement cell size in *dek1-4* and wild-type cotyledons and found no evidence that the cells differed in ploidy (22). However, true leaves were not examined.

In leaves and sepals, it is unknown whether giant cells exhibit a spatially ordered pattern across the organ, or if instead their spatial arrangement is random. Other epidermal cell types are non-randomly distributed across the leaf tissue. For instance, trichomes do not form in adjacent cells due to lateral inhibition mediated by diffusible signals (9), and stomata rarely differentiate in adjacent cells due to both lineage-specific division orientation and intercellular signaling (23). In contrast to stomata and trichomes, sepal giant cells can be in contact with one another. However, it is unknown whether such giant cell contacts are likely to be formed by chance. Due to their large shapes, quantifying the spatial arrangement of giant cells has remained challenging, and standard methods for assessing point pattern randomness are not applicable (24–27).

Here, we imaged and analyzed large areas of leaves to obtain a holistic understanding of both the cell-size distributions and the spatial arrangements of epidermal pavement cells in the leaf blade. We discovered that the genetic pathway that controls sepal giant cell formation also has a broader role in patterning epidermal pavement cell size in leaves. We quantified the spatial organization of giant cells using simulated randomized tissues and found that giant cells tend to cluster together in both mature leaves and sepals more than expected by chance, reflecting the tissue’s developmental history. Using modeling and data analysis, we found that giant cells emerge randomly in space at early stages of development, but the division of surrounding cells causes the spatial pattern to become non-random and clustered within the context of the whole tissue over time. Our computational modeling supports the notion that a non-random clustered pattern can emerge over developmental time in a cell-autonomous and stochastic manner.

## Results

### Arabidopsis leaves exhibit a large range of pavement cell sizes, similar to sepals

In sepals, giant cells are easily visible because they are highly elongated (Fig. S1A). Similarly, we observe large and highly anisotropic cells in cauline leaves (Fig. S1B). In rosette leaves, pavement cells of the epidermis are jigsaw puzzle-piece shaped with lobes and necks, such that cell size is not readily apparent by eye (Fig. S1C); however, heterogeneity in pavement cell sizes has been previously observed (4,28). Therefore, we wondered to what extent the distribution of cell sizes observed in sepals, ranging from giant cells to small cells, also occurs in rosette leaves. We imaged large sections of the blade (excluding midrib and margin cells) of leaf 1 or 2 from wild-type plants at 25 days post germination (dpg) when the leaves are fully expanded and mature. Leaves 1 and 2 initiate simultaneously and are indistinguishable; therefore, we refer to them interchangeably as leaf 1 or 2. We measured the area of the epidermal cells of leaves 1 or 2 and sepals on the abaxial (bottom) side (Fig. 2A, C). Throughout our analysis, we use the term size to mean cell area because area has been shown to be a more relevant measure of cell size than volume in highly vacuolated plant epidermal cells (4). We observed that the abaxial cell-size distributions for both sepals and leaves are asymmetric, with long tails representing large cells (Fig. 2E). Sepal giant cells are larger outliers in size and, consequently, the cell-size distribution in the sepal has a more extended tail than in the leaf (Fig. 2E). Still, we observed that the cell-size range in the leaf and sepal are similar and the largest cells of the sepal are about the same size as the largest cells of the leaf (Fig. 2A– E). We conclude that Arabidopsis leaves have a diverse range of cell sizes characterized by a long-tailed distribution, similar to the abaxial side of sepals.

**Fig. 2.**
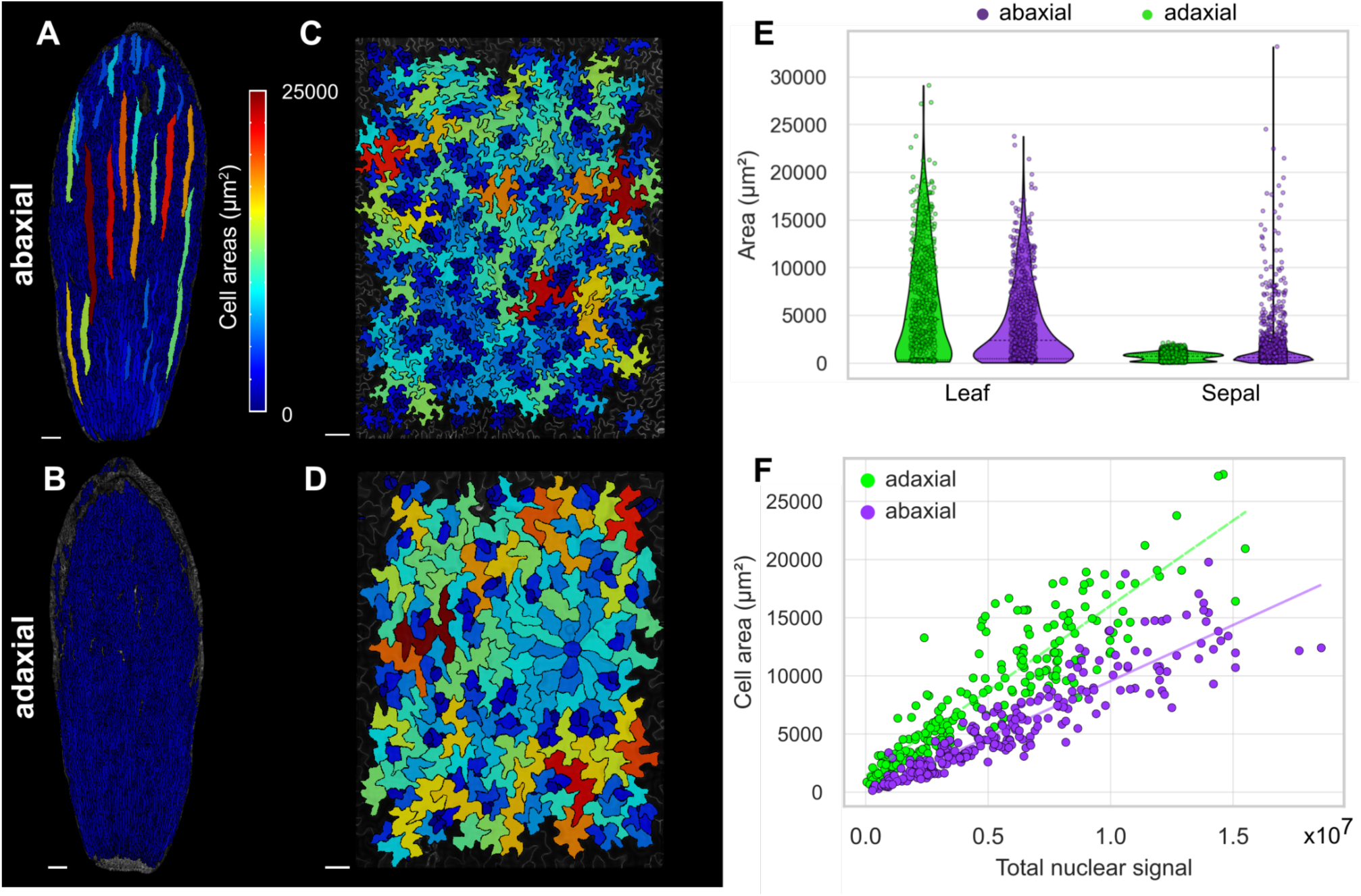
Abaxial and adaxial cell-size distribution in the wild-type leaf and sepal epidermis; size correlates with DNA content. (**A–D**) Cell area heat maps (in µm^2^) of (**A**) abaxial surface of wild-type sepal, (**B**) adaxial surface of wild-type sepal, (**C**) abaxial surface of 25-dpg wild-type leaf 1 or 2 (cell density: 234 cells mm^-2^) (**D**) adaxial surface of 25-dpg wild-type leaf 1 or 2 (cell density: 156 cells mm^-2^). Scale bars represent 100 µm. (**E**) Violin and strip plots of cell areas of abaxial and adaxial sides of 25-dpg wild-type leaves (two pooled replicates) and adult wild-type sepals (three pooled replicates). (**F**) Adaxial side (green) and abaxial side (purple) of 25-dpg leaf cell area versus DNA content as measured by H2B-TFP total nuclear fluorescence, with R^2^ = 0.85 for the abaxial side and R^2^ = 0.82 for the adaxial side (one of two replicates). See Fig. S2 for replicates. The underlying data for this figure can be found at Open Science Framework (<u>osf.io</u>), DOI:10.17605/OSF.IO/RFCWS.

### Large cells are formed on the adaxial side as well as the abaxial side of the leaf

In sepals, giant cells are restricted to the abaxial (outer) surface (Fig. 2A–B and Fig. S2A–D). We asked whether there was a difference in cell size between adaxial (top) and abaxial (bottom) surfaces of the leaf. Large cells of similar size are formed on both the adaxial and abaxial surfaces of the leaf, in contrast to the sepal (Fig. 2A–E and Fig. S2A–F). In addition, across the whole cell-size distribution, the cell density is lower and many cells are slightly more expanded on the adaxial side than the abaxial side (Fig. 2C–D and Fig. S2E–F). There are a greater number of stomata and stomatal lineage cells (very small cells) on the abaxial side compared with the adaxial side (Fig. 2C–E and Fig. S2E–F). We also observed that the abaxial cells are more lobed than the adaxial cells (Fig. 2C–D and Fig. S2E–F). Despite slight differences, the cell-size distributions of the abaxial and adaxial sides of the leaf are quite similar, particularly in the tails, where both sides exhibit a similar range of larger cells, in contrast to the sepal, where only the abaxial side has very large cells.

### Cell area correlates with DNA content

Cell area and ploidy are positively correlated in leaf epidermal cells (4). A recent study showed that fluorescence levels of histone fluorescence reporters measured from microscopy images are a good proxy to infer nuclear ploidy in Arabidopsis cotyledons (29). Hence, to validate whether cell area and ploidy were correlated in our leaves, we measured DNA content by quantifying total fluorescence of Histone 2B-TFP (*pUBQ10::H2B-TFP*) within each cell nucleus of the 25-dpg leaf images. For this reporter, the H2B-TFP signal distribution is noisy and continuous, not divided into four discrete peaks, providing an approximation of ploidy, not exact DNA content. As expected, a strong linear correlation between H2B-TFP fluorescence and cell area was observed for both the abaxial pavement cells (R^2^ = 0.85 and 0.91; n = 2) and the adaxial pavement cells (R^2^ = 0.79 and 0.82; n = 2) (Fig. 2F and Fig. S2G). Therefore, we focus on analyzing cell size, and infer that large cell size indicates high ploidy.

We wondered whether cells of similar size on the abaxial and adaxial side of the same leaf also have a similar DNA content. We found that cells of similar DNA content are larger on the adaxial side than on the abaxial side (Fig. 2F and Fig. S2G), suggesting that adaxial cells have expanded more than abaxial cells, as noted above. Because the largest sepal cells and the largest leaf cells had approximately the same areas, we asked whether the DNA content of these cells was also similar. We found that the total H2B-TFP fluorescence values of the largest cells were very similar between sepal and leaf, suggesting that these largest cells are similar in ploidy (Fig. S2H).

### Cell-size patterning emerges at the tip and progresses basipetally as the leaf differentiates

To determine how the cell-size pattern emerges in the leaf during development, we imaged both the adaxial and abaxial surfaces of each leaf at different stages of development. From 5 dpg to 9 dpg, cell size increases greatly (Fig. S3), as expected. At day 5, cells throughout the blade are fairly homogeneous in size, with a few cells starting to expand near the distal tip, and the large cells of the margin and overlying midrib already apparent (Fig. 3A). We focus on pavement cells in the blade and exclude margin and midrib cells from further analysis. The cell-size pattern consisting of large cells interspersed between small cells progressively develops basipetally from the tip (Fig. 3A–C), whereas at the base the cells remain uniformly small. The progression of cell-size patterning down the leaf is consistent with the well-established basipetal wavefront of differentiation and cessation of cell division (30). The cell area distributions showed that more large cells appear throughout development and the maximal cell size increases (Fig. 3A–B, D). By 9 dpg, cell size has been patterned almost to the base of the leaf (Fig. 3A–B).

**Fig. 3.**
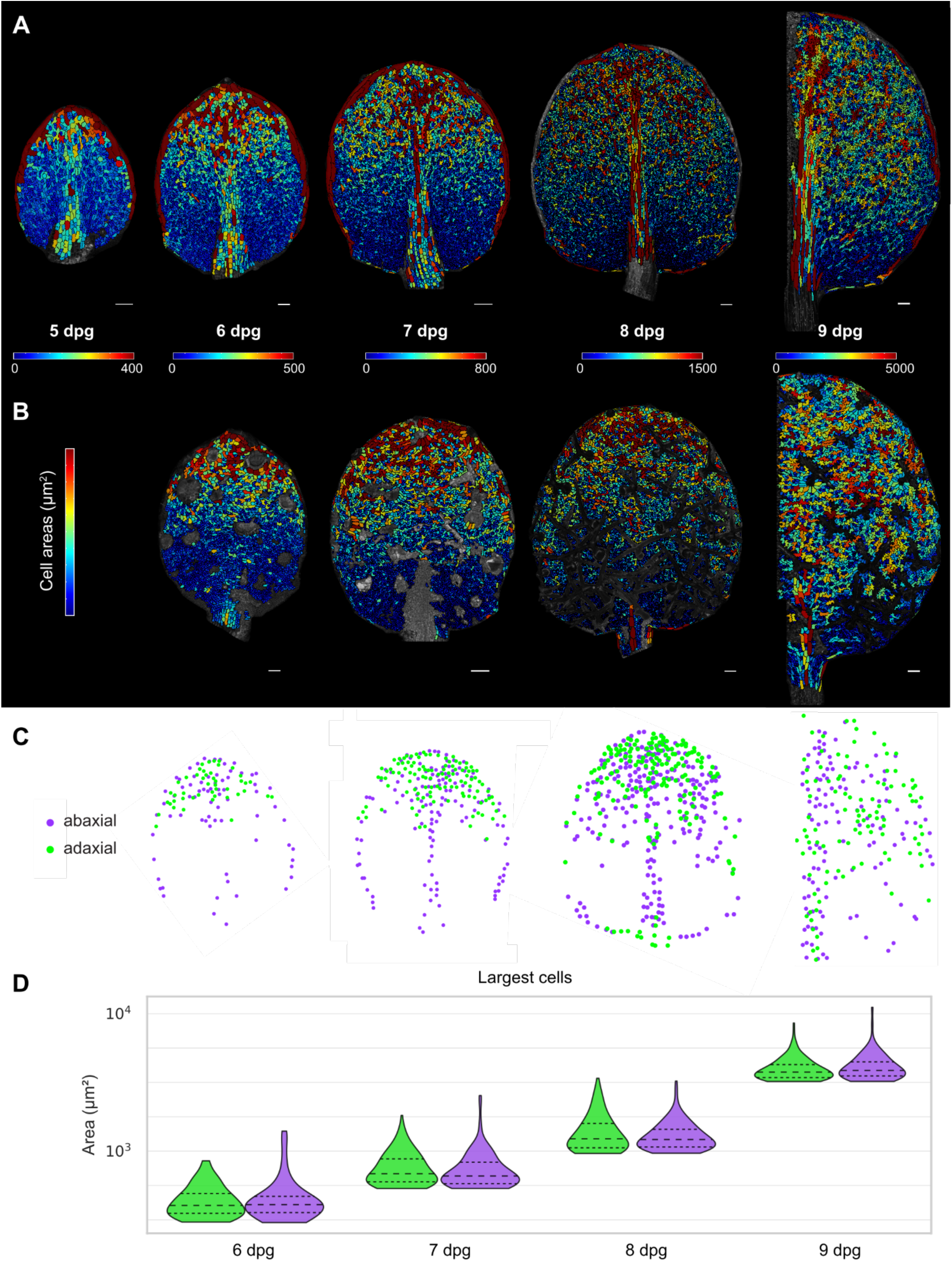
Cell-size patterning occurs as a basipetal wave simultaneously on the adaxial and abaxial sides of the leaf. (**A–B**) Cell area heat maps in µm^2^ of leaf 1 or 2 at different stages of development on (A) the abaxial side and on (B) the adaxial side of the same leaf at 5 dpg, 6 dpg, 7 dpg, 8 dpg and 9 dpg (half leaf). Unsegmented regions on adaxial leaves correspond to trichomes, which were not considered in this analysis. Each stage is associated with a distinct heat map color range. Scale bars represent 50 µm at 5 dpg and 6 dpg, and 100 µm at 7 dpg, 8 dpg and 9 dpg. (**C**) Spatial positions of the largest cells (those above an area threshold, see below) on the abaxial (purple points) and adaxial (green points) sides of the same leaf at 6 dpg, 7 dpg, 8 dpg and 9 dpg. Area thresholds for each leaf were determined from the 98th percentile cell area of the abaxial side. (Note that the midrib and margin cells are included in these overlays of large cell positions.) (**D**) Violin plots of cell areas (in µm^2^) of the largest cells (as defined in (C) and excluding margin and midrib cells) on abaxial and adaxial sides of leaves at different developmental stages. Abaxial and adaxial sides are from the same leaf. See also Fig. S3 for the leaves shown to scale. The underlying data for this figure can be found at Open Science Framework (<u>osf.io</u>), DOI:10.17605/OSF.IO/RFCWS.

We next asked whether the wavefront of cell-size patterning progresses basipetally at the same rate on the abaxial and adaxial sides of the leaf. Using images of both the abaxial and adaxial sides of the same leaf, we plotted the positions of the centers of the largest cells on both sides (including margin and midrib cells for landmarks) (Fig. 3C). Large pavement cells are observed in the same proximal–distal region on abaxial and adaxial sides during development (Fig. 3C). The region expands in the proximal direction as development progresses. This suggests that the wavefront of patterning and differentiation is coordinated across the abaxial/adaxial axis of the leaf.

### The sepal giant cell specification pathway also patterns giant cells in leaves

Because the cell-size distributions have similarities in leaves and sepals, we tested whether the giant cell specification pathway in sepals (Fig. 1H) also functions in the leaf to pattern cell size. We imaged leaf 1 or 2 at both 9 dpg and 25 dpg from wild type and giant cell pathway mutants. At 9 dpg, patterning has just extended to the base of the leaf, and the leaf is still small enough that we could image the whole upper abaxial quadrant to determine the pattern over a large fraction of the leaf blade (Fig. 4A and Figs. S4, S5). At 25 dpg, the leaf is fully differentiated, fully expanded, and the pattern is established (Fig. 4B and Figs. S6, S7). We found that cell-size patterning in the leaf is similarly affected in the mutants at both 9 dpg and 25 dpg as in the mature sepal (Fig. 4 and Fig. S8A–B). Notably, the largest cells show similar variations in their numbers across genotypes. Similar to the sepal, the size of the largest cells is moderately reduced in *acr4-2* mutants (Figs. 1B, 4A–D), and more greatly reduced in *atml1-3* mutants (Figs. 1C, 4A–D). The reduction in large cells is drastic in *dek1-4* and *lgo-2* mutant sepals and leaves, resulting in the absence of a long tail in the cell-size distribution (Figs. 1D–E, 4C–D). For these genotypes, the number of medium-sized cells is also substantially decreased (Figs. 1D–E, 4C–D). Conversely, the overexpression of *ATML1* (*ATML1-OX*) or *LGO* (*LGO-OX*) leads to an increase in the size of large cells and in fewer small cells compared to wild type, as in the sepal (Figs. 1F–G, 4A–D).

**Fig. 4.**
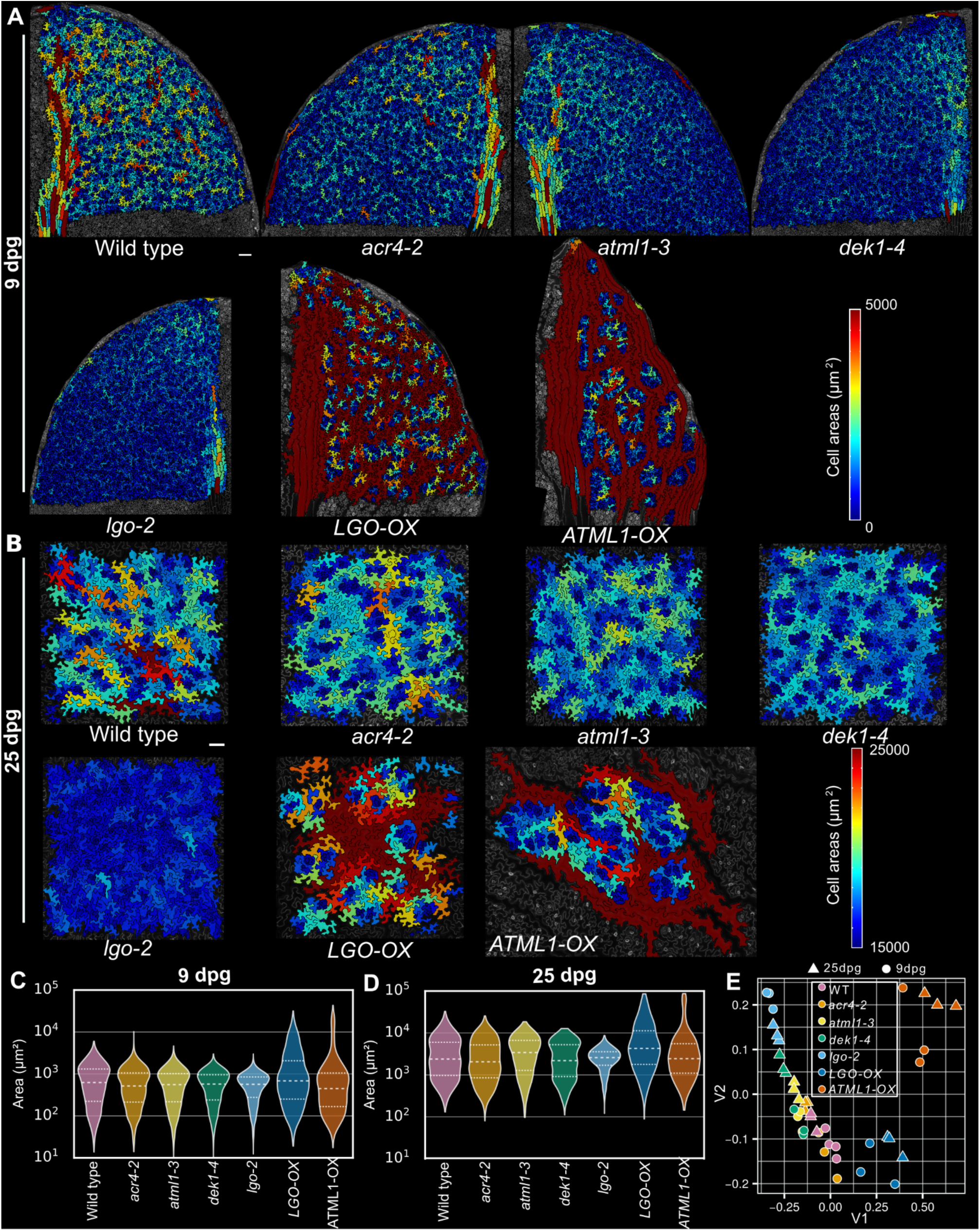
The sepal giant cell specification pathway also patterns cell size in leaves. (**A–B**) Cell area heat maps (in µm^2^) of the upper abaxial quadrant of leaf 1 or 2 at 9 dpg (A) and of a section on the abaxial side of leaf 1 or 2 at 25 dpg (B) for the following genotypes: wild type, *acr4-2*, *atml1-3*, *dek1-4*, *lgo-2*, *LGO-OX* (*pATML1::LGO*) and *ATML1-OX* (*pPDF1::FLAG-ATML1*). Scale bars represent 100 µm. Cell area heat maps of other replicates are shown in Figs. S4**–**S7. (**C–D**) Violin plots of cell area distributions on a log_10_ scale for 9-dpg replicates (C) and for 25- dpg replicates (D). Stomata were classified and removed in (C**–**D). Violin plots of individual replicates are shown in Fig. S8A**–**B. (**E**) Wasserstein distance plot of normalized cell-size distributions (see Materials and Methods) for all 9-dpg and 25-dpg replicates displayed as Euclidean distances embedded in 2D. The 25-dpg replicates are indicated by circular dots and 9- dpg replicates by triangular dots. The Wasserstein distance plot for 9-dpg and 25-dpg and Wasserstein statistical tests among replicates are shown in Figs. S4 and S6. Associated with Figs. S4–S8. The underlying data for this figure can be found at Open Science Framework (<u>osf.io</u>), DOI:10.17605/OSF.IO/RFCWS.

To quantify the variations in the number of large cells precisely, we quantitatively defined leaf giant cells on the basis of a cell area threshold. Specifically, we first classified pavement cells and stomata using a Support Vector Machine (SVM) classifier based on features of cell shape (see Materials and Methods, Fig. S9). Next, a cell-size threshold was established in the mature sepal and in the leaf, at both 9 dpg and 25 dpg, using the *atml1-3* mutants, which are known to have very few giant cells in sepals (see Materials and Methods, Fig. 1C and Fig. S9). Those cells in the 9- dpg and 25-dpg leaves as well as in the sepal that exceeded their associated threshold were categorized as giant cells (see cell-type classification outcomes in Figs. S10–S11). On the basis of this definition, we performed a quantitative comparison and statistically compared the number of giant cells per unit area among genotypes in leaves. Two-sample, two-tailed *t*-tests showed that in the 9-dpg leaf and the mature leaf, wild type had significantly more giant cells than *lgo-2* (9 dpg: p = 0.002, 25 dpg: p = 0.003), *dek1-4* (9 dpg: p = 0.002, 25 dpg: p = 0.002), *atml1-3* (9 dpg: p = 0.002, 25 dpg: p = 0.005), and *acr4-2* (9 dpg: p = 0.010, 25 dpg: p = 0.044). Conversely, *LGO-OX* had significantly more giant cells than wild type (9 dpg: p = 0.001, 25 dpg: p = 0.003). However, no statistically significant difference in the number of giant cells per unit area was observed between wild type and *ATML1-OX* (9 dpg: p = 0.213, 25 dpg: p = 0.75). Because the giant cells in *ATML1-OX* are so much bigger than wild-type giant cells, each of the giant cells in a given area of *ATML1-OX* leaf takes up a large amount of space, resulting in few giant cells per unit area despite the fact that most of the unit area is occupied by giant cells. We sought to quantify what was apparent visually by comparing the fractional area occupied by giant cells between *ATML1- OX* and wild type and found that the fractional area occupied by giant cells was significantly higher in *ATML1-OX* (9 dpg: p < 0.005, 25 dpg: p < 0.005). Thus, in *ATML1-OX* the number of giant cells is not changed but the fractional area covered by giant cells is increased.

Collectively, the similarities in the variation between the number of giant cells in the leaf and the sepal indicates that the sepal giant cell specification pathway also regulates the formation of giant cells in leaves.

### Giant cell mutants affect the entire cell-size distribution

We observed that not only are giant cells affected in these mutants, but other aspects of the cell-size distribution are also affected. For example, the number of medium-sized cells in *lgo-2* and *dek1-4* is reduced in addition to the number of giant cells (Figs. 4A–D) and, correspondingly, the number of small cells is increased in these mutants. To statistically analyze the difference in cell-size distributions, we conducted a principal coordinate analysis based on the Wasserstein distances between cell-size distributions (termed Wasserstein distance plot), which showed the difference between leaf samples according to their cell-size distributions on a 2-dimensional plane (Figs. S4H, S6H and S8C–F, see Materials and Methods). Samples clustered according to genotype, indicating that genotype controls the cell-size distribution. We observed a progressive increase in the number of giant cells along the first principal coordinate V1 from *lgo-2* mutants to *ATML1- OX* and *LGO-OX* (Fig. S4H and S6H). *ATML1-OX* and *LGO-OX* were distant from each other in this plot, which might partly reflect the fact that *LGO-OX* has more giant cells, whereas *ATML1- OX* has fewer but larger giant cells. When we created the combined Wasserstein distance plot using normalized cell-size distributions from both 9-dpg and 25-dpg leaves (see Materials and Methods), the samples continued to group according to genotype rather than developmental stage, further supporting the idea that these genes have affected the cell-size distribution by 9 dpg (Fig. 4E). Thus, we conclude that these genes affect the entire cell-size distribution.

However, some differences in the cell-size distribution are apparent between 9-dpg and mature 25- dpg leaves. Firstly, at 9 dpg, *dek1-4* and *lgo-2* mutants are very similar; however, in the fully mature 25-dpg leaves, the *lgo-2* cell-size range is notably smaller than that in the *dek1-4* mutant (Figs. 4A–D), suggesting that *lgo-2* cells continue to divide after 9 dpg. In addition, the small cells in *lgo-2* mutants were more uniform in size than all of the other genotypes because the typical small stomatal lineage cells that encircle the stomata in mature leaves were fewer in *lgo-2* (Figs. 4A–D). This altered cell-size distribution relates to the previous finding that *LGO* affects pavement cell differentiation in these stomatal lineage ground cells and that cells undergo division for a longer time in the absence of LGO (19). Secondly, although at 9 dpg the *LGO-OX* giant cells were slightly smaller than the *ATML1-OX* giant cells, at 25 dpg, the *LGO-OX* giant cells were nearly equivalent in size to *ATML1-OX* giant cells (Figs. 4A–D). In addition, we observed that more pavement cells were larger in *LGO-OX*, whereas only a few cells became giant in *ATML1-OX* (Figs. 4A–D). *ATML1-OX* leaves had a few connected giant cells separating large islands of small cells, whereas *LGO-OX* leaves showed more giant cells interspersed among smaller clusters of small cells (Figs. 4A–B). These phenotypic differences might reflect either inherent differences in ATML1 and LGO activities or the fact that *ATML1* and *LGO* overexpression transgenes are under the control of different promoters that might have differences in activity at different developmental stages.

### Relationship between the size and shape of cells and organs

In plants, compensation is the process by which organ size is maintained when cell number is altered by an accompanying change in cell size (31). We observed compensation in our leaf giant cell mutants (Fig. S12). Mature leaves of the mutants *acr4-2*, *atml1-3*, *dek1-4* and *lgo-2* are similar in size to wild-type leaves. Having fewer giant cells is compensated by having more small pavement cells (Fig. 4). However, *ATML1-OX* and *LGO-OX* mature leaves, which have much larger cells (see e.g. Fig. 4B, D), are smaller than wild-type (Fig. S12N–P). Therefore, only partial compensation for having fewer cells by having larger cells is observed in *ATML1-OX* and *LGO-OX* plants.

Additionally, *ATML1-OX* leaves are narrower than those of wild type and *LGO-OX* (Fig. S12A, F, G, I, N, O). We also observed that giant cells are more directionally elongated in *ATML1-OX* than in other genotypes (Figs. 4A–B and Figs. S4F–G, S5F–G, S6F–G, S7G–H), reflecting the elongated shape of the leaf. This suggests the existence of a relationship between giant cell shape and leaf morphology. Likewise, wild-type cauline leaves are both narrower and more elongated than wild-type rosette leaves, and also have more anisotropic elongated giant cells than in rosette leaves (Fig. S1). This observation supports the idea that cell shape reflects the anisotropy of the growing tissue (32).

### Spatial patterning of giant cells within the leaf blade

In wild-type plants, giant cells vary in position from sepal to sepal and from leaf to leaf (10–12). An open question has been whether the spatial organization of giant cells is random, or whether there is an underlying order. Classically, many specialized cell types such as stomata and trichomes are spaced such that they are not in direct contact to one another (23,33). Giant cells are frequently adjacent to each other and, therefore, it is clear that there is not a strong lateral inhibition between them. We set out to determine firstly whether giant cell position is correlated with underlying vasculature and secondly, how giant cells are spatially positioned relative to one another.

### Giant cells are not preferentially positioned overlying the vasculature

We wondered whether giant cell positioning was correlated with the position of leaf vasculature for two reasons. Firstly, we observed that large, highly endoreduplicated cells overlie the midrib of the leaf, extending all the way to the leaf tip (Fig. S13A). We wondered whether giant cells might be similarly preferentially located over the other veins. Secondly, we observed that large, highly endoreduplicated cells often appear to “peel” away from the midrib, as if following vascular branches (Fig. S13A). This phenomenon is most common in *ATML1-OX* leaves (Fig. S13C–F). To investigate whether giant cells overlie veins, we traced the veins from the original confocal image onto the heat map of cell area for a 9-dpg wild-type half leaf and four *ATML1-OX* half leaves. We found that many giant cells did not overlie the vasculature (Fig. S13B–F). Specifically, we noted that the points where giant cells peel off the midrib often do not align with where veins extend from the midvein. Furthermore, the orientation of giant cells do not follow the direction of the veins (Fig. S13B–F). Instead, veins in *ATML1-OX* plants frequently pass through patches of small cells (Fig. S13C–F). We conclude that vascular and giant cell patterns are not obviously correlated.

### Giant cells are clustered more often than expected by chance

A cell-autonomous and stochastic mechanism has been proposed to explain giant cell formation in the sepal (12). However, it remains unknown whether giant cells are randomly arranged within the tissue. To statistically assess the randomness of the pattern, we needed a random reference (or null model) to compare with our experimental replicates. Previous studies addressing this problem considered cells as points (26,34), or used a regular hexagonal grid to build a null model (35). In our case, these assumptions are not applicable due to the complexity of giant cell shapes and the heterogeneity of cell shapes and sizes that affect cellular arrangements (36). Therefore, we used the dmSET image-based method (36,37) to generate randomized tissues from the real segmented images (Fig. 5A and Fig. S14), allowing to randomly shuffle cell positions by preserving cell sizes and shapes of the original tissues (Fig. S15, see Materials and Methods). We generated 400 randomized tissues for each biological replicate for both the wild-type sepal and 25-dpg leaf.

**Fig. 5.**
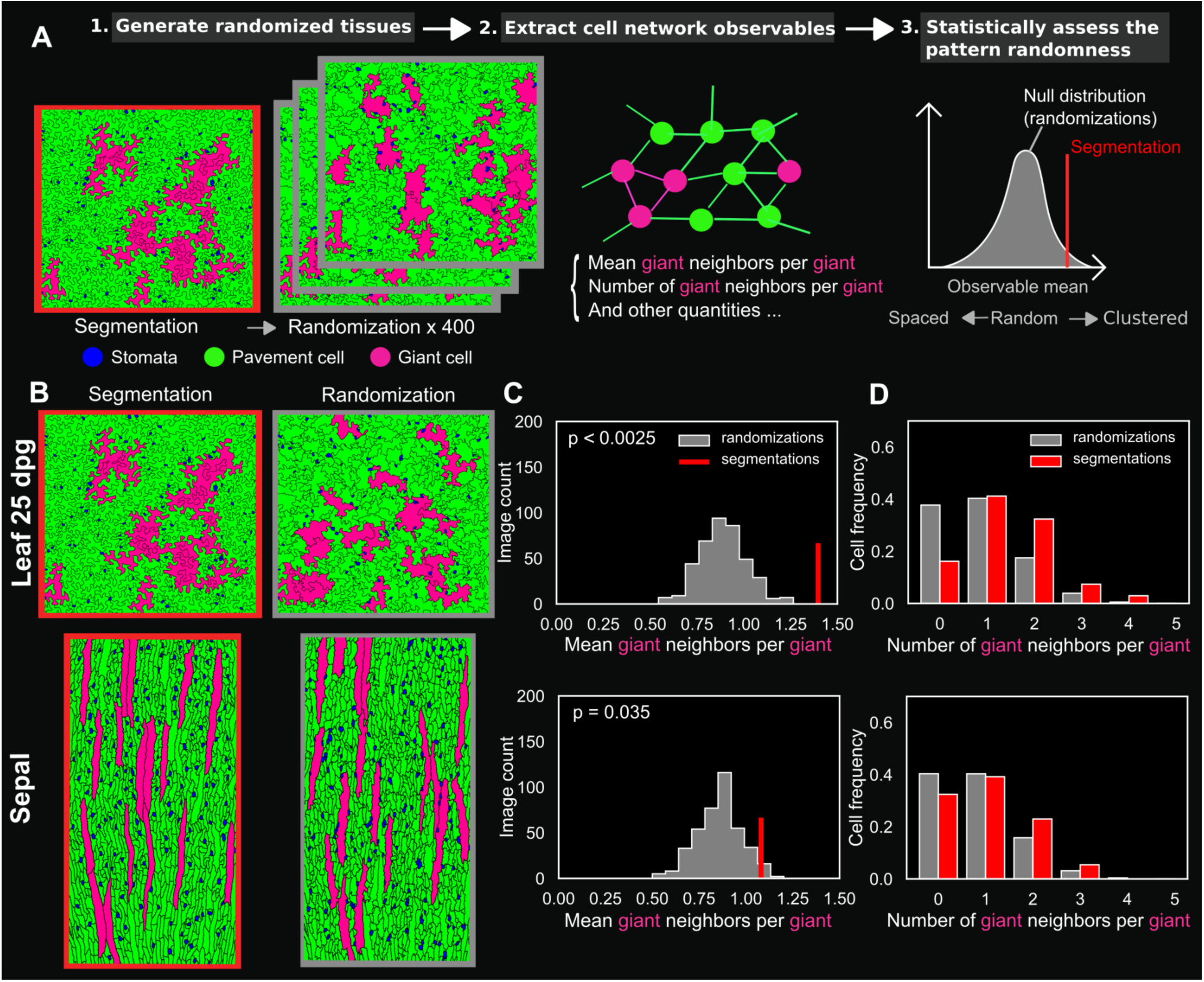
Giant cells are more clustered than expected in a randomized null model both in the wild-type leaf and sepal. (**A**) Scheme summarizing the method used to assess the randomness of the cellular patterns. Each segmentation is computationally randomized using the dmSET method into 400 randomized tissues where cell positions (and orientation in the case of the leaf) have been randomly shuffled (left; see Materials and Methods). To statistically assess the extent to which the segmented image shows a random giant cell pattern, a quantitative observable (middle) extracted from the segmentation is compared with the same observable computed in all randomized tissues, forming the estimated ’null distribution’ (right). (**B**) Example of a representative segmentation of a wild-type leaf 25 dpg (top left) and a wild-type sepal (bottom left) and one of their randomized tissue (randomization) images (right). (**C**) Mean number of giant cell neighbors (also referred to as giant neighbors) per giant cell (also referred to as giant) in leaves (top) and sepals (bottom). The value extracted from the segmentations (in red) was statistically tested against all the values extracted from the 400 pooled randomizations (in gray). The mean number of giant cell neighbors per giant cell is higher than expected in a randomized null model, and the null hypothesis can be rejected (p-value < 0.05), indicating that giant cells are clustered. (**D**) Distributions of the number of giant cell neighbors for all giant cells found in all replicates of segmentations (in red) and randomizations (in gray) in leaves (top) and sepals (bottom). Total number of giant cells counted (excluding giant cells at the image border) in the analysis: n = 68 (leaf, segmentations), n = 68 × 400 (leaf, randomizations), n = 74 (sepal, segmentations), n = 74 × 400 (sepal, randomizations). See also Figs. S14, S15, S16, S21 and S22. The code and data associated with this figure can be found at Open Science Framework (<u>osf.io</u>), DOI:10.17605/OSF.IO/RFCWS.

Several measures (Fig. S16), such as the mean number of giant cell neighbors per giant cell, which captures the amount of contacts between giant cells, were computed in the experimental data and the corresponding randomized tissues. To statistically assess the randomness of the giant cell pattern, these measures in the real biological tissues (segmentation) were compared with the same measures in all the randomized tissues (randomizations), which formed a null distribution (Fig. 5A and Fig. S16).

In the randomized tissues, cell sizes were well preserved, but cell shapes were affected in the leaf (Fig. 5A–B and Fig. S15D–E). To ensure that these shape artifacts did not introduce bias in our analyses, we tested our method on a randomly selected population of cells in both leaves and sepals (see Materials and Methods) and confirmed the absence of significant bias in the null models (Fig. S17). Additionally, we reconstructed the original leaf tissues with shape artifacts similar to those in the randomized tissues (Fig. S18), and found that giant cell connectivity was largely preserved and that the results remained consistent (Fig. S18) (see Materials and Methods).

For both wild type 25-dpg leaves and mature sepals, when considering the six pooled replicates, the mean number of giant cell neighbors per giant cell was greater than in a randomized null model, and the null hypothesis could be rejected (p < 0.05) (Fig. 5C). This result shows the presence of clustering among giant cells both in the leaf and the sepal. It was less probable to find isolated giant cells, and more probable to find giant cells in contact with two or more other giant cells compared with what was expected by chance (Fig. 5D). Similar results were found in the leaf using an alternative randomization method we developed based on cutting and merging cells (Fig. S19). The non-random pattern of giant cells was also supported by the analysis of other spatial measures (Fig. S16). The similar distribution of the number of giant cell neighbors in leaves and sepals (Fig. 5C–D) reflects a similar spatial organization, supporting the idea of common patterning mechanisms.

### Different cell sizes are organized into different spatial patterns

To investigate whether the clustered pattern is exclusive to giant cells, we applied the same analysis to distinct sub-populations of pavement cells in the leaf tissues. Four populations of pavement cells were defined: giant cells (Fig. 6A–C), middle-sized cells (Fig. 6D–F), small cells (Fig. 6G–I), and a control population of randomly selected pavement cells of any size (Fig. 6J–L). In contrast to the clustered pattern of giant cells (Fig. 6A–C), middle-sized pavement cells exhibited a more random organization (Fig. 6D–F; the null hypothesis could not be rejected, with p = 0.195). Conversely, small pavement cells showed a clustered organization (Fig. 6G–I), because the mean number of neighbors between small pavement cells significantly exceeded the value observed in the randomized tissues. Notably, these small cells were clustered around the stomata, and their spatial arrangement is probably a consequence of the stomatal patterning process. A random cellular pattern was found in randomly selected pavement cells, as expected (Fig. 6J–L). Overall, these analyses highlight a relationship between pavement cell size and cell spatial organization within the tissue. Furthermore, these findings underscore the distinctive clustered arrangement of giant cells in comparison to middle-sized and randomly selected pavement cells.

**Fig. 6.**
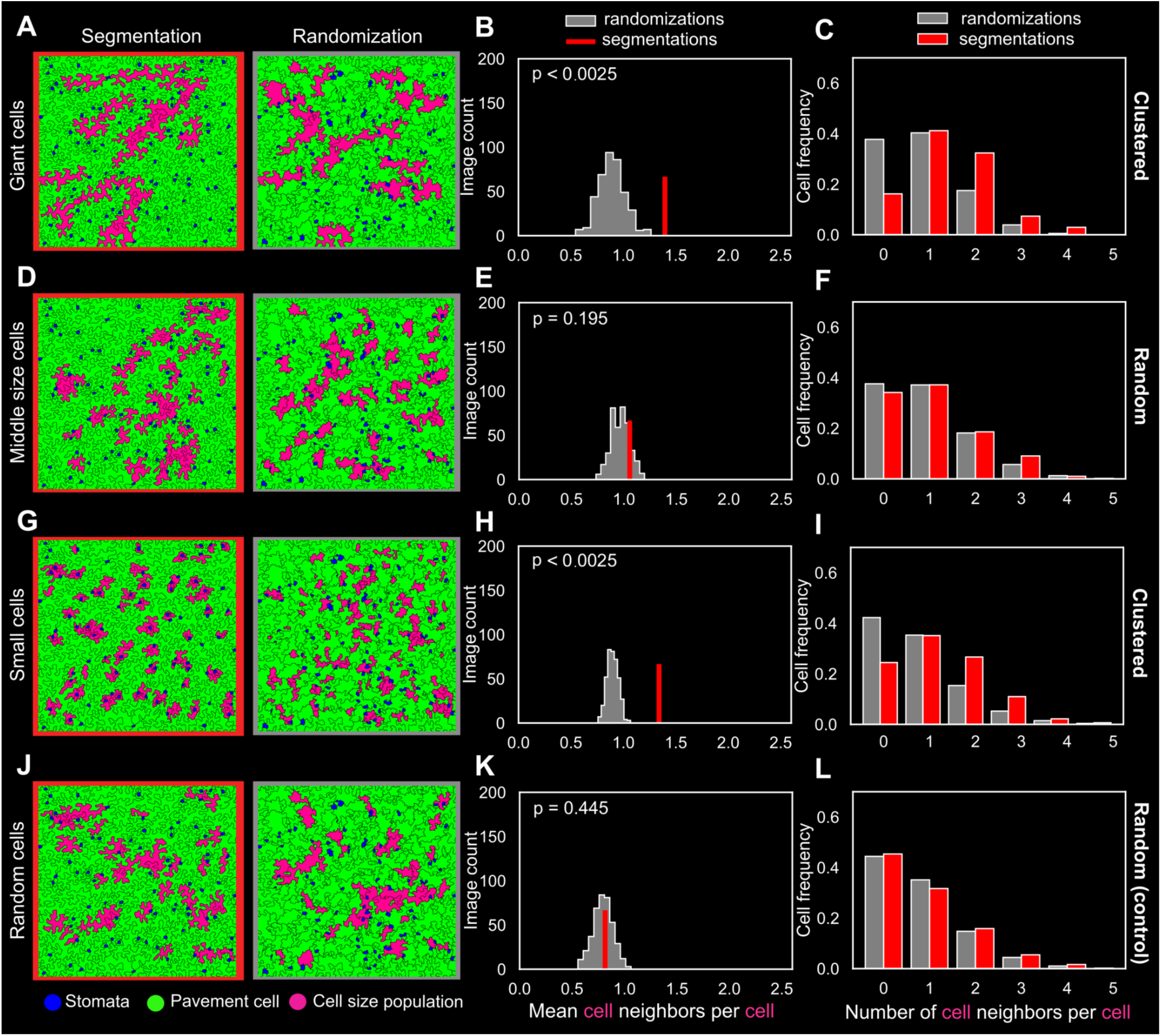
Different cell sizes display different spatial patterns in the wild-type leaf. The method used to assess the randomness of the giant cell patterns (Fig. 5) was applied here on different pavement cell-size populations within the mature 25-dpg leaf: (**A–C**) giant, (**D–F**) mid-size (around 5,000 µm^2^), (**G–I**) small (smallest pavement cells), and (**J–L**) random (randomly selected pavement cells). The number of cells in each category was determined such that the total cell area of the cell population was approximately equal to the area occupied by the giant cells. (**A**, **D**, **G**, **J**) Example of representative segmentation of a 25-dpg wild-type leaf (left) and one of its corresponding randomized tissue randomization (right), where cell locations have been computationally shuffled. Cells colored in magenta represent the cells within the studied pavement cell-size population. (**B**, **E**, **H**, **K**) Mean number of cell neighbors per cell within the same size population. (**B**) The mean number of giant cell neighbors per giant cell is higher than expected by chance (p < 0.05), indicating that giant cells are clustered. Same data as in Fig. 5C, top. (**E**) Middle-size cells are less clustered than giant cells and more randomly organized (the null hypothesis cannot be rejected, p = 0.195). (**H**) The mean number of small cell neighbors per small cell is significantly higher than in the randomized tissues (p < 0.05), highlighting that small cells form clusters. (**K**) As expected, the randomly selected pavement cells (with area > 2,000 µm^2^) show a value that falls right in the center of the null distribution (p = 0.445). (**C**, **F**, **I**, **L**) Distributions of the number of cell neighbors belonging to the studied cell population per cell of that population in the segmentations (in red) and the randomizations (in gray). All six replicates were pooled together. Total number of cells in cell populations counted in the analysis: n = 68 (giant cells, segmentations), n = 68 × 400 (giant cells, randomizations), n = 199 (middle-size cells, segmentations), n = 199 × 400 (middle-size cells, randomizations), n = 639 (small cells, segmentations), n = 639 × 400 (small cells, randomizations), n = 162 (random cells, segmentations), n = 162 × 400 (random cells, randomizations). The code and data associated with this figure can be found at Open Science Framework (<u>osf.io</u>), DOI:10.17605/OSF.IO/RFCWS.

It was previously shown that, apart from small cells, pavement cells follow the theoretical topological laws based on space-filling (i.e., via entropic considerations), with larger cells being on average surrounded by smaller ones in young *spch* leaf tissues (38). Our analyses on more mature wild-type leaves reveal that larger cells are surrounded by larger cells (and have fewer neighbors) than what is expected by purely random space-filling given by our null model (Fig. 6, Fig. S18D).

### A cell-autonomous stochastic model can recapitulate giant cell clustering

To investigate how giant cell clustering emerges during leaf and sepal epidermal development, we wondered whether the existing cell-autonomous and stochastic model for giant cell specification in sepals (12) could also recapitulate the clustered feature of the giant cell pattern. In this multicellular computational model, the concentration of ATML1 stochastically fluctuates and is regulated by a self-catalytic feedback loop. ATML1 regulates the expression of a downstream cell-cycle regulator target (Fig. 7A). At the end of a cell cycle, a cell either divides or endoreduplicates if the ATML1 target exceeds a specific threshold during the G2 phase. We used this model (12) to investigate the resulting spatial organization of giant cells in simulated tissues (Fig. 7A–B; see Materials and Methods).

**Fig. 7.**
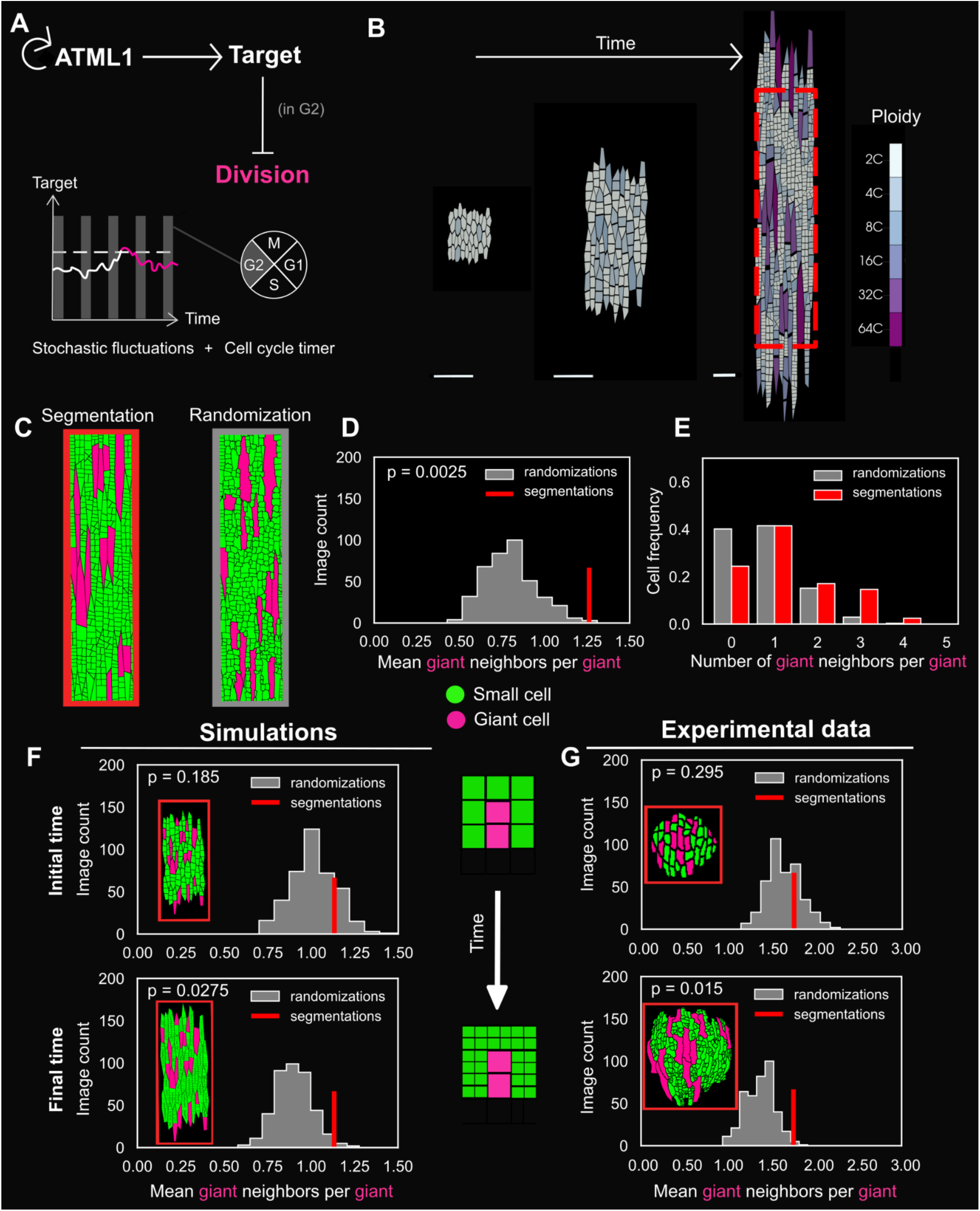
A cell-autonomous stochastic model can recapitulate giant cell clustering because of cell divisions of surrounding cells. (**A**) Cartoon of the computational model for giant cell patterning. ATML1 activates a target (LGO), which, if above a certain threshold and during the G2 cell-cycle phase, prevents cell division and instead drives the entry into endoreduplication and giant cell formation. (**B**) Simulation snapshots of the simulated growing sepal, at three different time points. Color codes indicate the cell ploidy levels. Scale bars represent the same size in arbitrary units. (**C**) A rectangular section of the simulation output (e.g., see rectangle shown in (B)) is used to quantify the giant cell pattern. “Segmentation” refers to one simulation output (left) and “Randomization” to one randomization of the simulated output (right). Giant cells, labeled in magenta, were defined by a size threshold (see Material and Methods). (**D**) Mean number of giant cell neighbors (also referred to as giant neighbors) per giant cell (also referred to as giant) in the simulations (called segmentation in red) and in its randomizations (in gray). The mean number of giant cell neighbors per giant cell is higher than expected in a randomized null model (p < 0.05), indicating a clustered pattern of giant cells. (**E**) Distribution of the number of giant cell neighbors per giant cell. The results obtained in (D, E) are comparable with the results in experimental sepal replicates (Fig. 5C, D). Five simulation outputs with five different initial conditions were performed and combined for the analysis. Total number of giant cells (excluding giant cells at the image border) counted in the analysis: n = 42 (segmentations), n = 42 × 400 (randomizations). (**F– G**) Statistical assessment of the randomness of the giant cell pattern (comparing the “segmentations” in red with the randomized tissues in gray) at initial time (top) and final time point (bottom) in (**F**) the simulations (at t = 55 and t = 135) and (**G**) the real tissues (at stage 4 and stage 9). At the initial time point, the null hypothesis could not be rejected but the mean giant cell neighbors per giant cell became significantly greater than in a randomized null model (p < 0.05) at the final time point. At the left of panel (F), a cartoon represents two neighboring giant cells surrounded by an increasing number of cells as the tissue develops. All five replicates (in simulations) and three replicates (in experimental data) were pooled. Total number of giant cells counted in the analysis: n = 83 (simulations, segmentations), n = 83 × 400 (simulations, randomizations), n = 49 (experimental data, segmentations), n = 14 × 400 (experimental data, randomizations). The dataset in (C) was also used for an independent analysis in Hervieux et al. 2016. See randomization snapshots related to this figure in Fig. S20. The code and data associated with this figure can be found at Open Science Framework (<u>osf.io</u>), DOI:10.17605/OSF.IO/RFCWS.

To assess the randomness of the simulated giant cell pattern, we applied the same method as in the experimental images (Fig. 5A) to images of the final simulation time point (Fig. 7B–C). Giant cells were also defined by a size threshold, which was established such that all cells of ploidy 16C or above were considered to be giant (see Materials and Methods). We observed that the mean number of giant cell neighbors per giant cell was greater than expected if giant cells were randomly distributed (p < 0.05, Fig. 7D), showing that the current cell-autonomous model can also produce a clustered giant cell pattern. Furthermore, the distribution of the number of giant cell neighbors per giant cell (Fig. 7E) was similar to the distribution observed in the experimental sepals (Fig. 5D, bottom). This raises the question of what mechanisms are responsible for cell clustering in a cell-autonomous, multicellular model of dividing cells.

### Cell division contributes to the clustering of giant cells

To understand how the giant cell clustering behavior emerges in our computational model, we analyzed how the cellular spatial pattern changes over time within the tissue. We hypothesize that the initial giant cell pattern arises randomly throughout the epidermis, due to the stochastic nature of ATML1 concentration fluctuations that trigger endoreduplication, and occasionally lead to giant cell contacts. As non-giant cells continue to divide, giant cells would appear more clustered in the fully grown tissue. To test this hypothesis in our simulations, we selected the first-arising giant cells (see Materials and Methods) and quantified their spatial organization both at an early time point and at the end of the simulation (Fig. 7F and Fig. S20A–B). We found that these giant cells were more randomly distributed at the initial time point, as the null hypothesis could not be rejected (p = 0.185, Fig. 7F), whereas they were more clustered compared to the randomized tissues at the final time point (p < 0.05, Fig. 7F). Indeed, although the giant cell contacts were preserved over time (red bar in Fig. 7F), we observed a shift in the null distribution of the mean number of giant neighbors per giant cell between the initial and the final time point (Fig. 7F). As new cells arise from cell division, the number of potential cellular configurations (i.e., the number of possible spatial cellular arrangements) increases, which decreases the probability of observing giant cell clusters under a random model where all cells have random positions. Therefore, even if giant cell contacts are preserved, their arrangement in the context of the entire tissue becomes more clustered over time.

To investigate the emergence of the giant cell spatial pattern over time in real tissues, we used time-lapse data of developing sepals (39), where cells were tracked over time, and we similarly quantified the patterns of the first-arising giant cells at the first available time point (sepal at stage 4, 24-h time point) and a later one (sepal at stage 9, 120-h time point; see Materials and Methods) (Fig. 7G and Fig. S20C–D). Similar to the simulations, we observed that giant cells were more randomly distributed in younger sepals and were more clustered in the more developed sepals when compared to the randomized tissues (Fig. 7G). This analysis indicates that the stochastic and cell-autonomous model is a plausible model to explain the spatial organization of giant cells. Moreover, it shows that cell clustering can emerge in a growing tissue without the need for cell– cell communication but instead as a result of cell divisions.

## Discussion

We investigated pavement cell-size patterning in the Arabidopsis leaf epidermis. We found that the same genetic pathway that controls giant cell formation in sepals also controls cell size and giant cell formation in the leaf. Specifically, the receptor-like kinase ACR4, the transcription factor ATML1, the calpain protease DEK1, and the CDK inhibitor LGO are important for the formation of leaf giant cells. Just as in the sepal, overexpression of *LGO* results in an increased number of giant cells and overexpression of *ATML1* leads to a larger area occupied by giant cells. Although giant cells are only present on the abaxial epidermis of sepals, they are present on both the abaxial and adaxial surfaces of leaves. We observed that giant cells are scattered across the surface, sometimes in contact with one another, in both leaves and sepals. Our analysis demonstrated that giant cells are more likely to be in contact than in a randomized tissue null model in both organs.

Many patterning systems rely on cell–cell communication to generate proper spacing (7,40), and the emergence of clustered patterns in certain cell types is often attributed to cell–cell communication mechanisms in static tissues (36,41,42). However, giant cell specification occurs within the context of tissue growth and cell division. Therefore, it is important to consider the influence of these dynamic factors as well. We revisited our previous cell-autonomous model for giant cell specification in which ATML1 stochastically fluctuates, and confirmed that giant cell clustering could arise in that model as a result of cell division, without the need for cell–cell communication. To understand how clustering emerges, we tracked giant cells from their initial emergence both in our modeled tissue and in published experimental time-lapse data of growing sepals (39), and analyzed the evolution of their spatial pattern over time. This analysis suggested that the giant cell pattern initially arises randomly in space in the primordium and becomes more clustered in the context of the fully grown tissue. Therefore, the decrease in the randomness of the giant cell pattern over time appears to be caused by the division of cells surrounding the giant cells, including dividing stomata lineage cells. Rather than resulting from active cell–cell communication, giant cell clustering reflects the history of cell division and growth of the tissue. Nevertheless, in a proliferating tissue, other mechanisms might operate at the same time that result in giant cell clustering. For instance, correlative effects on cells belonging to the same lineage might influence cell fate decisions, e.g., due to the inheritance of molecular factors from mother to daughter cells (43). Ultimately, further experimental and theoretical studies are needed to determine the relative importance of cell–cell communication and cell proliferation in leading to giant cell clustering.

In the past, researchers have attempted to increase organ size by increasing cell size by promoting endoreduplication, but these efforts have not been successful (44). This is because compensation occurs, in which smaller cell size is accompanied by an increase in cell number, so that organ size is relatively conserved (45,46). Consistent with this, we observed that leaf 1 or 2 of wild type, *atml1-3*, and *lgo-2* plants are approximately the same size at maturity. Furthermore, instead of having larger leaves, the *ATML1-OX* and *LGO-OX* genotypes that have larger cells actually have slightly smaller leaves than the wild type at maturity. These observations are consistent with what is observed for sepals, because *ATML1-OX* and *LGO-OX* sepals are slightly smaller than wild-type sepals (10,45). We have previously shown that mitotic division substitutes for endoreduplication to compensate and maintain organ size in mutants lacking giant cells (10). Our images suggest that this mechanism also operates in leaves.

Although giant cell number does not greatly influence organ size, organ shape is altered in sepals and leaves. *ATML1-OX* and *LGO-OX* sepals are narrower than those of wild type and curve outward, so that the bud opens prematurely (11). We speculate that the anisotropy of sepal giant cells drives the change in sepal shape. In *ATML1-OX* leaves, where giant cells are highly anisotropic, we observe a similar change in leaf shape, in which *ATML1-OX* leaf 1 and 2 are more pointed and oblong compared with the rounded wild-type leaf 1 and 2. By contrast, giant cells in *LGO-OX* leaves are isotropic and are more similar to wild-type giant cells, and *LGO-OX* leaves are more rounded. Our results suggest that ATML1 is sufficient to induce anisotropic cell growth, whereas LGO is not. Tang et al. (2023) have shown that the change in shape between rounded juvenile rosette leaves and more elongated adult rosette leaves is accompanied by the appearance of highly anisotropic giant cells at the leaf base (32). However, they showed that loss of these directional, elongated giant cells does not change adult rosette leaf shape; the adult rosette leaf 7 remains elongated in *lgo-2* leaves where giant cells are not present (32). Thus, the relationship between giant cell shape, anisotropic growth, and organ shape is complex. Further work at the single-cell level will be needed to elucidate the influence of giant cells on the shapes of different tissues.

Despite the similarities between cell-size patterning in leaves and sepals, subtle differences also exist. Firstly, the distribution of epidermal cell sizes in the leaf is broader than in the sepal, where cells are fairly uniformly small except for a scattering of giant cells (Fig. 2E). Secondly, leaves have giant cells on both abaxial and adaxial blades, whereas sepals have giant cells only on the abaxial side and not on the adaxial side that faces the petals. The petal blade does not have giant cells on either abaxial or adaxial sides (47); thus, sepals might be an organ whose identity is transitional between vegetative and floral organs (48). We observe a similar phase change in the anisotropy of giant cells. Rosette leaf giant cells are jigsaw puzzle-piece shaped and relatively isotropic. Later in the plant life cycle, giant cells in cauline leaves begin to be more anisotropic along the proximal–distal axis and start to resemble sepal giant cells. This supports the hypothesis that cauline leaves represent an intermediate state between rosette leaves and sepals (49). Finally, sepal giant cells are highly anisotropic along the proximal–distal axis. Although sepals and leaves have notable yet subtle differences in cell size, cell-size patterning is regulated by the same developmental pathway in both organs.

The genetic pathway that regulates giant cell specification has been co-opted from the epidermal specification pathway, which is a developmental pathway necessary for epidermal and thus plant development (14,50,51). Without proper epidermal specification, the plant embryo will not progress past the globular stage of development (50–52). The fact that this fundamental epidermal developmental pathway also patterns giant cells illustrates a common theme in development, namely, that regulatory proteins are commonly reused for more than one developmental process (53).

Taken together, our analysis and theoretical work on patterning during tissue growth highlights that unexpected effects can occur, and that these are difficult to infer from the canonical view of pattern formation arising in a static tissue. In this instance, an initially random pattern of giant cells becomes non-random as the surrounding cells divide. Thus, the effects of cell proliferation might also be important to determine the spatial distribution of specialized cell types in other tissues.

## Materials and Methods

### Plant growth conditions

All seeds were sown on LM111 soil in pots and were stratified in the dark for 3 days at 4°C. The pots were then transferred to Percival plant growth chambers set to 60% humidity, 22°C temperature, and 24-h light provided by Philips 800 Series 32-Watt fluorescent bulbs (f32t8/tl841) (∼100 μmol m^−2^ s^−1^). Days post germination (dpg) were counted from the time the pots were transferred to plant growth chambers.

### Cloning fluorescent nuclear markers

To create a teal fluorescent (TFP) nuclear marker ubiquitously expressed under the UBIQUITIN 10 promoter (*pUBQ10::H2B-TFP; pAR393*), an H2B-TFP fusion with an AAAPAAAAA linker was generated by PCR. TFP was amplified by PCR with primers oAR440 (5′-gct gcc gct cca gct gca gct gcc gct ATG GTT TCT AAG GGA GAA GAA ACT ACT ATG-3′) and oAR438 (5′-cct cga gtc aCT TAT AAA GTT CAT CCA TAC CAT CAG TAG-3′). The lower-case letters in the primer sequences represent linkers, restriction sites, and cloning sequences that were added to the gene sequences. H2B was PCR amplified with oAR369 (5′-CAC CGG ATC CAC AAT GGC GAA GGC AGA TAA G-3′) and oAR439 (5′-agc ggc agc tgc agc tgg agc ggc agc AGA ACT CGT AAA CTT CGT AAC CGC CTT AG-3′). Sequences encoding H2B-TFP were fused via overlapping PCR with oAR369 and oAR438 primers. The H2B-TFP PCR product was cloned into pENTR D TOPO to create the pAR198 entry clone. H2B-TFP was recombined into pUB-Dest with LR Clonase II according to the manufacturer’s instructions to generate *pUBQ10::H2B-TFP* (pAR393). pAR393 was transformed into Arabidopsis Col-0 plants expressing the pLH13 *p35S::mCitrine-RCI2A* yellow fluorescent plasma membrane marker (45) via Agrobacterium tumefaciens (strain GV3101)-mediated floral dipping (54) and selection with glufosinate-ammonium (“Basta” Neta Scientific OAK-044851-25g).

### Generation of mutant plant lines containing fluorescent plasma membrane and nuclear markers for imaging

Mutant alleles and overexpression transgenes were crossed to plants expressing fluorescent cell-membrane and nuclear markers to obtain plants for imaging. The following mutant alleles were used: *acr4-2*, *atml1-3*, *dek1-4*, and *lgo-2*. In addition, two lines overexpressing either ATML1 (*pPDF1::FLAG-ATML1*) or *LGO* (*pATML1::LGO*) in a Col-0 background were used. All of these alleles/transgenes are in the Columbia-0 (Col-0) accession. *acr4-2* (SAIL_240_B04) contains a T-DNA insertion in the codon of the second of seven 39-amino acid repeats of the beta propeller extracellular domain, which is upstream of the transmembrane domain and the kinase domain and is therefore presumed to be loss-of-function allele. The *acr4-2* mutant was obtained from Gwyneth Ingram (14), who obtained it from Syngenta (55). The *atml1-3* allele is a T-DNA insertion in the homeodomain and is a loss-of-function mutant (11). The *atml1-3* was obtained from the Arabidopsis Biological Resource Center (ABRC; accession number SALK_033408) (11). The *dek1-4* allele contains a point mutation that changes a conserved arginine to a cysteine within calpain domain III (ABRC accession CS68904) (11). Complete loss of function of DEK1 is lethal (50); therefore, *dek1-4* must retain some function. The *dek1-4* phenotype is recessive and therefore, *dek1-4* is likely hypomorphic (11). The *dek1-4* mutant was originally isolated in the Landsberg erecta accession and was subsequently back-crossed twice into Col-0 (12). *lgo-2* contains a T-DNA insertion within the coding sequence of the gene and is a loss-of-function allele (10). *lgo-2* was obtained from the ABRC (accession number SALK_033905 and is available as a homozygous mutant as accession CS69160). *pPDF1::FLAG-ATML1* (*ATML1-OX*) was obtained from Gwyneth Ingram (56). *pATML1::LGO* (*LGO-OX*) has been deposited for distribution at ABRC under accession CS69162 (11,45). *acr4-2*, *atml1-3*, *dek1-4*, *lgo-2*, *pPDF1::FLAG-ATML1*, and *pATML1::LGO* were each crossed to plants expressing both a *p35S::mCitrine-RCI2A* fluorescent plasma membrane marker (pLH13) and a *pUBQ::H2B-TFP* fluorescent nuclear marker (pAR393). The F2 progeny were genotyped for *acr4-2*, *atml1-3*, *dek1-4*, and *lgo-2* (primer sequences in Table S1) and lines were isolated that were homozygous for these alleles and that also expressed both *pUBQ::H2B-TFP* and *p35S::mCitrine-RCI2A*. We could not obtain *atml1-3* homozygous plants that also contained the *p35S::mCitrine-RCI2A* transgene after crossing, which was probably because the *ATML1* gene was linked to the insertion site of the *p35S::mCitrine-RCI2A* transgene. To obtain plants expressing *p35S::mCitrine-RCI2A* in a homozygous *atml1-3* background, the plasmid containing p35S::mCitrine-RCI2A was transformed into *atml1-3* homozygous plants with *pUBQ::H2B-TFP* through Agrobacterium tumefaciens (strain GV3101)-mediated floral dipping (53). A T1 line was chosen that strongly expressed the mCitrine membrane signal and this line was used for future experiments. For the overexpression transgenes *pPDF1::FLAG-ATML1* and *pATML1::LGO*, seeds were collected from F2 plants and F3 plants were genotyped for *pPDF1::FLAG-ATML1* or *pATML1::LGO* (Table S1). Those F2 plants that produced only F3 plants having *pPDF1::FLAG-ATML1* or *pATML1::LGO* were isolated as homozygous for *pPDF1::FLAG-ATML1* or *pATML1::LGO*, respectively. Those lines with *pPDF1::FLAG-ATML1* or *pATML1::LGO* homozygous and that expressed both *pUBQ::H2B-TFP* and *p35S::mCitrine- RCI2A* were used for imaging.

### Sample preparation for imaging

Leaves and sepals were mounted in 0.01% (v/v) Triton X-100 for imaging. Leaves were imaged between two coverslips, and sepals were imaged on a slide with a coverslip. Curvy leaves were cut with a razor blade to ensure they could be placed flat under the coverslip. Samples were imaged immediately after preparation. Sepals were imaged at stage 14 (57).

### Imaging with confocal microscopy

A ZEISS LSM 710 Axio Examiner confocal microscope with a W Plan-Apochromat 20×/1.0 DIC D-0.17 M27 75 mm water-immersion objective lens was used to image leaf 1 or 2 of the Arabidopsis rosette and mature (stage 14) sepals. A 458 nm laser was used to excite *pUBQ::H2B- TFP* (collection range 463–500 nm) and a 514 nm laser was used to excite *p35S::mCitrine-RCI2A* (collection range 525–645 nm). Images were captured with a 1× zoom. The gain and laser power varied slightly between images to accommodate slight differences in signal intensity between samples. Each image was composed of several tiles. The dimensions of each voxel were 0.415 µm (x) by 0.415 µm (y) by 1 µm (z).

For the images to compare cell size with nuclear fluorescence on the abaxial and adaxial faces of the same organ (two leaf replicates and three sepal replicates; Fig. 2 and Fig. S2), the 458-nm laser power and gain used for imaging *pUBQ::H2B-TFP* were adjusted so that the TFP signal was below saturation and was then held constant for all images.

Leaf areas were calculated from confocal images of entire leaves taken using a 2.5× objective for each 9-dpg and 25-dpg leaf replicate.

### Image processing

Tiles were stitched in the horizontal direction by ZEISS stitching software (overlap of 5% and threshold of 0.7) and in the vertical direction with MorphoGraphX (58,59) using the process “Stacks/Multistack/Merge Stacks” (parameters: method = max; interpolation = linear). Assembled images were saved as MorphoGraphX stack files. A surface mesh was created in MorphoGraphX from each image to perform segmentation and analysis on the epidermis. First, extraneous parts of the image were removed with the Voxel Edit tool. (Such extraneous parts of the image include trichomes on the adaxial images and pollen grains/nematode eggs on some leaf images.) Then, an image was subjected to Gaussian Blur using the process "Stack/Filter/Gaussian Blur Stack" (parameters: x = 2; y = 2; z = 2). Next, the tissue surface was identified with the process “Stack/Morphology/Edge Detect” (parameters: threshold varied between 2,300–7,000 according to individual image brightness; multiplier = 2.0; adapt factor = 0.3; fill value = 30,000). These steps extracted a surface of the leaf. The process “Stack/Morphology/Fill Holes” was applied to some images when holes were apparent in the surface (parameters: x-radius = 20; y-radius = 20; threshold = 10,000; depth = 0; fill value = 30,000). This surface was then used to generate a mesh with the process “Mesh/Creation/Marching Cube Surface” (parameters: cube size = 5 μm; threshold = 20,000). The mesh was smoothed with “Mesh/Structure/Smooth mesh” (parameters: number of passes varied between 20–45; Walls Only = no). The mesh obtained was then subdivided with the process “Mesh/Structure/Subdivide” either once or twice depending on its size. In order to obtain the cell membrane signal on the surface, the process “Meshes/Signal/Project Signal” (parameters: min/max distances ranged between 5–15 μm; MinSig = 0.0; MaxSig = 60,000) was used to project the *p35S::mCitrine-RCI2A* plasma membrane marker original signal onto the mesh at an optimal depth. The depth range yielding the clearest cell membrane signals with minimal distortion was selected. To perform cell segmentation, each individual cell in the leaf was first manually identified with a cell label marking (seed). Using these seeds, watershed segmentation was performed using the process “Meshes/Segmentation/Watershed Segmentation”. Adjacent pairs of stomatal guard cells were segmented together to form a single cell; these were classified as stomata. Errors in segmentation were identified and corrected by removing the label for those cells and reseeding. The Heat Map processes computed the cell area and other morphological cell features as well as the position of every cell. The cell area data was exported into a data table file for each image. In addition, other cellular shape features were computed and exported into a data table for the cell type classification (Materials and Methods: cell type classification).

To analyze the nuclear signal from images of leaves expressing *pUBQ::H2B-TFP*, vertical stitching of tiles (already horizontally stitched with Zeiss stitching software) was performed in MorphoGraphX (max method and linear interpolation). Because the *TFP* reporter was expressed under the *UBIQUITIN 10* promoter, TFP was localized in nuclei of the mesophyll cells in addition to cells of the epidermis. Mesophyll nuclei were removed with the Voxel Edit tool. Nuclei were identified as being from the mesophyll by lining up the nuclear signal images with their corresponding membrane signal images and comparing the nuclei within the bounds of each epidermal cell membrane. When compared with an epidermal cell nucleus, mesophyll cell nuclei were often dimmer and lower down and therefore, excess nuclei were removed according to these criteria so that each epidermal cell had one nucleus. When it was ambiguous which of two nuclei in a single cell was from the mesophyll or epidermis, both nuclei were removed and excluded from the analysis. Segmentation of the nuclei was performed in MorphoGraphX so that the total signal could be calculated for each nucleus. To do so, the confocal image was first subjected to “Stack/Filters/Brighten Darken” (parameter: 1). Next, a gaussian blur was performed using “Stack/Filters/Gaussian Blur” (parameters: x = 1, y = 1, z = 1), followed by a binarization with the process “Stack/Filters/Binarize” (parameters: threshold = 2,000), which functioned to select pixels above a threshold value to identify the edges of each nucleus. A lower threshold value was chosen so that we could identify the entire nuclei even for dim nuclei. The Voxel edit tool was used to separate nuclei that inflated into one another. We then created a mesh from the binarized image using the process “Mesh/Creation. Marching Cubes 3D” (parameters: cube size = 1, min voxel = 0, smooth passes = 3, label = 0). To ensure that the mesh covered all fluorescence of each nucleus, we expanded the mesh using “Mesh/Structure/Shrink Mesh” with a negative value (parameter: distance = -1). Individual nuclei were manually seeded and then the watershed segmentation was performed with the process “Mesh/Segmentation/Watershed Segmentation” to identify each nucleus. The Heat Map function calculated the total H2B-TFP fluorescence within each nucleus, as a representation of DNA content. To study correlations between total nuclear H2B-TFP signal and cell size, individual cells from cell area meshes were matched with their constituent nuclei from nuclear signal meshes using MorphoGraphX parent tracking. For the leaf replicates, total nuclear H2B-TFP signal was calculated for as many cells as possible from both the abaxial and adaxial sides. For the sepal replicates, total nuclear H2B-TFP signal was calculated only on the abaxial side and only for the largest cells.

To create the heat maps overlaid with vasculature in Fig. S13, confocal images of leaves expressing *p35S::mCitrine-RCI2A* were used to create surfaces and were segmented as described above to create cell area heat maps. The mCitrine-RCI2A confocal images were found to have signal in the vasculature, so that the trajectories of veins could be traced in images from the abaxial surface of the image, one can see. The mCitrine-RCI2A confocal images were transformed around the *z-*axis in MorphoGraphX. For each leaf, the cell area heat map and the mCitrine-RCI2A confocal image transformed around the *z-*axis were aligned in MorphoGraphX and PNG screen captures were taken of each. These PNGs were then loaded into Adobe Illustrator and the veins were traced in white onto the heat maps.

Please note that wild-type 25 dpg leaf replicates 1, 3, and 4, *lgo-2* 25 dpg leaf replicates 1 and 2, and *LGO-OX* replicates 1, 2, and 3 were used for an independent analysis of cell shape (specifically lobeyness) (60).

### Statistical Analysis

To analyze the relationship between total nuclear H2B-TFP signal (DNA content) and cell area for the leaves in Fig. 2 and Fig. S2, linear regressions were performed on R statistical software (https://www.r-project.org/). To compare the total nuclear H2B-TFP signal (DNA content) of the cells of largest area between sepals and leaves, the cell area at the 98^th^ percentile was calculated for each of the three abaxial sepal replicates and these three cell areas were averaged for an area threshold of 4,308 µm^2^. Cell area versus total nuclear H2B-TFP signal (DNA content) was plotted for cells above this 4,308 µm^2^ area threshold for the abaxial sepals and the abaxial and adaxial leaves.

To compare positions of the largest cells on the abaxial and adaxial sides of each leaf at different stages of development (Fig. 3), the abaxial and adaxial images were aligned in MorphoGraphX. Then, cell area heat maps were created and the x and y coordinates of the center of each cell were calculated (Fig. 3C). Cell area thresholds for each leaf were determined from the 98^th^ percentile cell area of the abaxial side, and the positions of cells above these area thresholds were plotted for the abaxial and adaxial images of each leaf.

R statistical software was used to analyze the cell-size distributions and create the violin plots and Wasserstein plots. The threshold for significance was set to alpha = 0.05. To create the Wasserstein plots, a Wasserstein test was performed between each pair of replicate distributions. A test statistic (also known as Wasserstein distance) and p-values were returned for every test. The p-values are listed in Fig. S8C–D. Classical multidimensional scaling was performed to create a 2D coordinate for each replicate distribution based on the Wasserstein distances, and points from these coordinates were plotted. To ensure that the distances between 2D points adequately reflected the Wasserstein distances among replicate distributions, we plotted the Wasserstein distances against the Euclidean distances between points (Fig. S8E–F). The linear relationships between Wasserstein distances and Euclidean distances showed that the 2D graph accurately represents the differences between distributions.

To create the Wasserstein plot of the combined 9-dpg and 25-dpg cell area data, cell areas of each replicate in each genotype were normalized by the mean cell area for that replicate. In this way, each replicate has a mean of 1. This eliminated the difference in the mean values of the cell size between the 9-dpg and 25-dpg leaves, such that all higher moments of the area distributions could be compared rather than the absolute sizes.

To statistically compare the differences in the number of giant cells across genotypes in the leaf at 9 dpg and at 25 dpg, two-sample, two-tailed *t*-tests that assumed equal variance were performed on the number of giant cells per segmented area between wild type and the different genotypes.

To statistically assess the randomness of the cellular patterns, see section “Statistical analysis of the cellular patterns” below.

### Cell type classification

To automatically distinguish stomata from pavement cells, a supervised classification algorithm was used based on cell shape features (Fig. S9). Cell shape features were computed from each 2.5D mesh using the MorphoGraphX (58,59) process “Mesh/Heat Map/Analysis/Cell Analysis 2D” and were extracted with “Mesh/Attributes/Save as CSV” into a data table. Three distinct training datasets were created using a single wild-type replicate – one for the sepal, one for the leaf at 25 dpg and one for the leaf at 9 dpg. To get the different training datasets, we manually selected some pavement cells and stomata and labeled them as different cell types, ran the classification processes available within MorphoGraphX, and manually corrected the cells that were wrongly identified. These training datasets were then used to train a supervised learning algorithm (Support Vector Machine quadratic) using the Classification Learner App in MATLAB (61,62) . The following cellular shape features were selected to train the classifier in the 25-dpg leaf: area, average radius, length of the major axis, maximum radius, perimeter, circularity, lobeyness (ratio of the cell perimeter over that of its convex hull; convex shapes have lobeyness 1) (59), and rectangularity (ratio of the cell area over the area of the minimum bounding rectangle in the cell) (59). For the sepal, the aspect ratio and the length of the minor axis were also taken into account. For the 9-dpg leaf, where the variety of cell types was more complex, three cell types were defined (pavement cell, meristemoid and stomata) and meristemoid and stomata were combined in the post-processing script. The shape features used to train the classifier were area, average radius, minimum radius, perimeter, circularity, lobeyness, and visibility stomata (it counts the proportion of straight lines that connects the cell outline without passing through a cell boundary)(59,63). To automatically predict cell types in all replicates, a developed MATLAB script containing the trained classifier and a post-classifier filter, which corrects for potentially wrong predictions on the basis of known shape criteria, was applied. Manual corrections were finally performed, in which misclassified cells were re-labeled with their correct cell type.

Giant cells were defined by a cell-size threshold (Fig. S9). Because a few giant cells were expected in *atml1-3* mutants, *atml1-3* mutants were used as a reference to build this threshold. Fewer than 0.7% of the pavement cells were considered to be giant cells in *atml1-3* tissues, which was supported through visual observation in the sepal. Consequently, the giant cell-size threshold was set as that corresponding to the average between the 99.3rd percentile cell-size value with the cell-size value immediately above it in the distribution, taking into account the data of three *atml1-3* pooled replicates. For consistency, the same method was applied to the sepal, and to 9-dpg and 25- dpg leaves, which gave three different threshold values (sepal: 5,290 µm^2^, leaf 9 dpg: 2,570 µm^2^, leaf 25 dpg: 14,160 µm^2^). The percentiles were only calculated on rectangular sections (omitting cells at the outline of the organs) of the sepal to maintain consistency across different organs. Classification output examples in different genotypes are shown in Figs. S10 and S11.

### Randomization of the experimental images

To assess the randomness of the cellular patterns, it was essential to establish a random reference, or null model, against which the observed pattern could be compared. To produce the required random reference, the image-based method dmSET (36,37) was applied to generate 400 synthetic random equivalent tissues from each segmented image. Cell positions and orientations were randomly shuffled into new images (named randomizations) (37), while preserving individual approximate cell shapes and sizes (Fig. S15). Only the incomplete cells at the border of the images were fixed. This randomization method avoids potential biases arising from the heterogeneity of cell sizes and shapes in the tissues, which affect the number of neighboring cells. We ensured that cellular properties, and more specifically cell area and cell lobeyness (defined as the perimeter of the cell divided by the perimeter of its convex hull), were approximately conserved in the randomized leaf tissues (Fig. S15D–E; the Pearson coefficient was 0.98 for cell area correlation and was 0.94 for cell lobeyness correlation). In the sepal randomized tissues, cell orientations were constrained between -𝜋/6 and +𝜋/6 compared with their initial orientation, to maintain the anisotropy of the tissue. Cell shape properties were also approximately conserved (Fig. S15I–J). A custom-made MATLAB script was subsequently applied to both original and randomized images to correct errors introduced by the dmSET method and to compute cell shape properties and cellular network information that was used to quantify the cellular pattern.

Before randomizing the different sepal and leaf replicate images (Fig. 5, and Fig. S14), each 2.5D mesh was first converted into a 2D pixel image using the process “Stack/Mesh Interaction/Mesh To Image” (with a pixel size of 1μm) in MorphoGraphX. Subsequently, a square crop (in the leaf) or rectangular crop (in the sepal) that maximized the tissue section was performed in the segmented images. These 2D segmented images were then randomized using the dmSET method.

To study the change in the giant cell spatial pattern over time, published time-lapse sepal data were used (39) that were randomized at two different time points (sepal at stage 4: 24 h, and at stage 8: 96 h). Cell segmentation and cell lineage tracking were already performed in (39). Using MorphoGraphX, sepal cells were manually selected at the later time point, and the exact corresponding mother cells at the first time point were established using the lineage tracking analysis from (39). In order to quantify the spatial pattern of the same giant cells at two different time points, giant cells at both stages were defined as the pavement cells that did not divide during this period of time. This approach allowed the comparison of the change in tissue organization consistently at two different time points. Then, the 2.5D meshes were projected into 2D images. These images were subsequently randomized using the dmSET method. Here, to facilitate the study of the same giant cells over time, the images were not cropped and the entire studied tissue was randomized, including the cells at the edges. To achieve this, the background region, located outside the tissue of interest, was considered as a single cell that remained fixed in the randomized tissues. Examples of randomizations are shown in Fig. S20C–D. Three different sepal replicates were used for these analyses of the time-lapse data.

“Segmentation” and “Randomization” images appearing in figures such as Figs. 5 and 6 and Figs. S14, S17, S18, and S20 were produced with a Python script using the multi-labeled images generated by dmSET.

Several approaches were used to test the robustness of our method and ensure that there was no bias in our null model. In the randomized tissues, cell sizes were well preserved (Fig. S15D), but cell shapes were affected in the leaf (Fig. S15E). Because the leaf pavement cells in the randomizations exhibit different shapes (more convex shapes and noisier edges) compared to the original cells, we developed an additional alternative method for generating randomized tissues (Fig. S19), called Cut and Merge Cells (CMC). In this approach, each leaf replicate was first over-segmented using MorphoGraphX (autoseeding with r=4 μm combined with manual seeding) to create templates composed of small fragments of pavement cells (Fig. S19A). These templates were then used to automatically generate a random pattern of giant cells, preserving their original sizes and numbers (Fig. S19A). Giant cells were created recursively: their positions were randomly allocated to a first pavement cell fragment, which was then merged with neighboring cell fragments until the original giant cell size was reached (Fig. S19A). Similar results were obtained by using this randomization method (Fig. S19B–C), confirming the tendency of giant cells to cluster more than would be expected by chance. However, while cell shapes produced by this method exhibited less noisy edges, they remained less lobed and more convex compared to those in the original segmentation (Fig. S19F). Moreover, this method does not randomize all cells in the tissue, such as stomata.

We wondered whether the shape artifacts introduced by the dmSET method could introduce bias into our null model by affecting the contacts between cells in the randomized tissues differently than in the original segmentations. To explore this possibility, we generated “reconstructions” of the original images using the same dmSET method as for the randomizations, except that each cell’s position from the initial segmentation was preserved (with some added uniform noise having a maximum of ±5 μm in the y and x directions) (Fig. S18A–C, E–F). This resulted in cell shape artifacts that were nearly identical between the reconstructions and the randomizations (Fig. S18J– L), making them more comparable. Indeed, the differences in cell lobeyness (defined as the perimeter of the cell divided by the perimeter of its convex hull) from the original tissues were similar between the reconstructions and randomizations for large cells (Fig. S18J). The number of cell neighbors differed slightly from the original segmentations due to the introduced shape artifacts (Fig. S18M–N), but we found that giant cell contacts remained mostly the same in the reconstructions. Specifically, we observed approximately 6% fewer contacts between giant cells in the reconstructions compared with the segmentations. When comparing reconstructions with randomizations (which are comparable because of their similar cell shapes), the results remained consistent (Fig. S18O), with giant cells being more clustered than expected by chance (p = 0.005). In the sepal, cell shape was quite preserved (Fig. S15I–J; the Pearson coefficient was > 0.99 for cell area and perimeter). Sepal cell edges are more noisy in the randomized tissues, with a higher cell perimeter (Fig. S15J); this could potentially lead to more contacts in the randomized tissues, which would not affect our conclusion.

Furthermore, we evaluated our method by using a randomly selected population of pavement cells in both leaves and sepals (Fig. S17). The null hypothesis could not be rejected and this result was robust across five different random patterns in all replicates (one is shown in Fig. S17). Specifically, we obtained p-values ranging from 0.395 to 0.488 in leaves and from 0.178 to 0.335 in sepals. The number of giant cell contacts was either slightly more or less than the expected mean random value, attributable to variability in the random patterns. This indicates that the artificial random pattern did not deviate from true randomness, suggesting that our null models do not present a significant bias.

### Statistical analysis of the cellular patterns

By comparing a spatial observable in the cellular network of the actual segmentation with the corresponding observable in the cellular networks of the 400 generated randomized tissues, whether the considered observable is likely to be observed by chance can be statistically tested (36). Hence, the use of this method on observables measuring distances or contacts between the studied cells allows the assessment of whether the arrangement of the cells within the tissue is random, clustered or dispersed (Fig. 5A).

To quantify the patterns, a custom-made Python script was used to extract pertinent observables (i.e., spatial quantities) from the cellular network, which used the NetworkX Python library (64). In this manuscript, we mainly focused on the number of giant cell neighbors per giant cell to quantify the number of local contacts between giant cells. Other observables have been quantified, such as the minimum shortest path between giant cells, and the number of giant cells in a cluster (Fig. S16). When dealing with cropped images, giant cells (or any cell population studied, see in Fig. 6) at the border of the image were not considered in the analysis.

The number of giant cell neighbors was extracted for every giant cell, and the mean number of giant cell neighbors per giant cell across all giant cells was computed within each experimental replicate. Similarly to the methodology described by the authors of the dmSET method (36), the mean value extracted from the segmentation image was compared with the approximated null distribution formed by the 400 mean values extracted from the randomized images. We first performed the analysis on each replicate independently (Figs. S21 and S22). As the cell-size distributions in the different replicates showed similarities across replicates (Fig. 5 and Fig. S21 and S22), replicates were pooled to increase the sample size and statistical power. Six image replicates were used for both the leaf and sepal wild-type (Fig. 5), and three replicates were used for the wild-type sepal time-lapsed images (Fig. 7). To test the null hypothesis assessing the randomness of the observed metric, a p-value p was obtained as the ratio of the number of random images (defined here as one random image resulting from pooling one random image per replicate) displaying the same or a more extreme value than the one obtained in the segmentation replicates (one-sided test). If the value fell within the null distribution with an associated high p-value (p > 0.05), the null hypothesis could not be rejected, indicating that the observed quantity could likely be expected by chance. If the value fell outside the null distribution, we assigned p < 0.0025, with 0.0025 corresponding to the inverse of the number of random images (400) used to create the null distribution.

In addition, the distribution of the number of giant cell neighbors for all giant cells from the pooled experimental replicates was studied, which provided more insights into their spatial organization. This was compared qualitatively with the distribution expected in a random tissue, extracted from the 400 randomized tissues of all replicates.

All plots derived from these analyses were performed with Python, with the use of the matplotlib (65) and seaborn packages (66).

### Mathematical model for giant cell fate commitment and numerical simulations

To simulate the giant cell fate decisions, our published stochastic and cell-autonomous multicellular model in a growing tissue was used (12). In that model, the transcription factor ATML1 stochastically fluctuates and drives the expression of its target LGO. In the simulated growing tissue, cells divide using a timer with some stochasticity. When the timer of a cell reaches a threshold Θ_C,S_, cells undergo the S-phase, and therefore cells transition from being diploid (2C) to tetraploid (4C). By default, cells that reach a second and higher timer threshold Θ_C,D_ will undergo division. However, those cells that have reached a certain LGO concentration threshold Θ_T_ after undergoing the S-phase will not divide and are maintained in an endoreduplication cycle, which increases their ploidy.

To model the dynamics of the concentrations for ATML1 and LGO, and the Timer variable, we use chemical Langevin equations (67), which are differential equations with a corresponding deterministic part, consisting of production, degradation and regulatory terms, followed by a stochastic part modelling thermodynamic fluctuations that contains a square root, whose radicand has the sum of the absolute values of the production, degradation, and regulatory terms. The dynamics of the ATML1 concentration, LGO concentration, and Timer variable in cell *i*, denoted by [*ATML1*]_i_, [*Target*]_i_ and *Timer*_i_ respectively, follow the Langevin equations given by

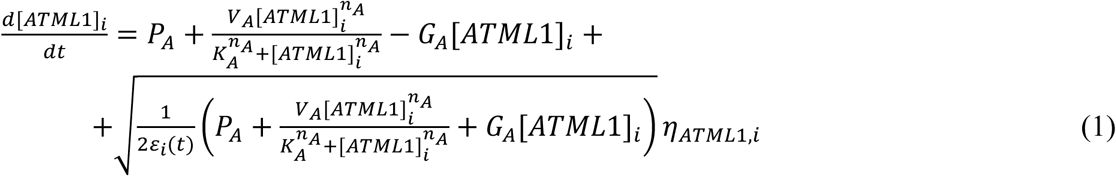

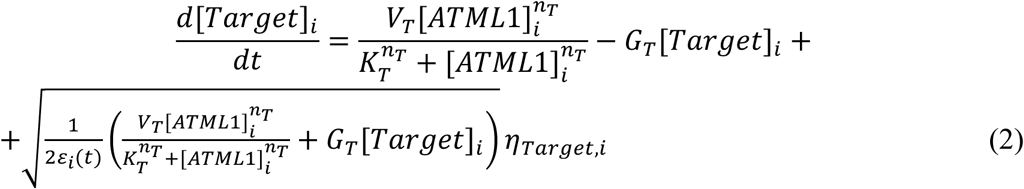

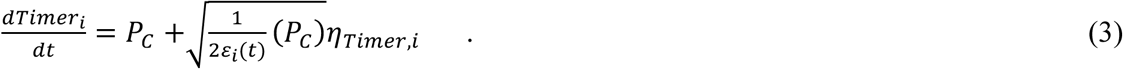

Eqn. (1) stands for the rate of change of ATML1 in cell *i*, and its terms on the right-hand side describe constitutive expression, self-activation (implemented via a Hill function) (68), linear degradation, and the corresponding stochastic term in ATML1; Eqn. (2), stands for the rate of change of the Target in cell *i*, and its terms on the right-hand side describe ATML1-induced expression of the Target (using also a Hill function), linear degradation, and the corresponding stochastic term in the Target; Eqn. (3) stands for the rate of change of the Timer in cell *i*, and its terms on the right-hand side are a constitutive production and the corresponding stochastic term. The parameters of the equations are as follows: *P_X_* is the basal production rate for the *X* variable (where *X* is either *A* for ATML1, *T* for Target concentration or *C* for the Timer variable), *V_X_* is the prefactor of the ATML1-dependent production rate for the *X* variable, *K_X_* is the ATML1 concentration at which the ATML1-dependent production rate has its half-maximal value, *n_X_* is the Hill coefficient, and *G_X_* is the linear degradation rate for the *X* variable. *ε_i_(t)* is a normalized cell area, *ε_i_(t)* = *E_0_E_i_(t),* where *E_0_* is an effective cell area, and *E_i_(t)* is the area of cell *i* in arbitrary units. *η_Xi_* is a random Gaussian variable with zero mean that fulfills〈*η_Xi_(t)η_X’j_(t’)*〉= *δ(t-t’) δ_XX’_δ_ij_*, where *i* and *j* are cell indices, *X* and *X’* the modeled variables, *δ_XX’_* and *δ_ij_* are Kronecker deltas and *δ*(*t-t’*) is the Dirac delta function.

Upon cell division, the Timer was reset. To implement the resetting, the following rule was applied at each time step:

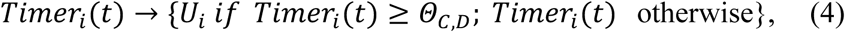

where *U_i_* is a uniform randomly distributed number in the interval [0, 0.5) and *Θ_C,D_* is the cell division threshold for the Timer.

The multicellular template on which the simulations were run and initial conditions were the same as in Meyer et al. (2017). Initial conditions for ATML1 and Target were randomly uniformly distributed in the interval of [0,1) and [0,0.1), respectively. The Timer initial conditions were set in correlation with the cell areas in the initial template with some stochasticity, as performed in Meyer et al. (2017). The differences between the used initial conditions were just in the ATML1, Target and Timer initial cellular values, determined by different random numbers.

Tissue growth and division were also implemented as in Meyer et al. (2017). The multicellular tissue grows anisotropically, to emulate the patterning process in the sepal. This anisotropic growth was implemented by imposing a displacement of the vertices with respect to the center of mass of the tissue, with a given radial and vertical exponential rate. After each simulation step, dilution effects due to growth in the modeled variables were taken into account. Cells divided using the shortest path rule together with the constraint of having the division plane through the center of mass of the cell. We assumed that molecules are homogeneously distributed within cells, and therefore, upon cell division, sister cells have the same concentration of ATML1 and Target variables at birth, but can have different cell sizes.

Numerical simulations were performed with Tissue software (12,13,69), and the integration was performed using an Îto interpretation of the Langevin equations with a Heun algorithm (70). Integration was performed with a time step dt = 0.1, and simulations were stopped at time 135. Parameter values for the simulations are given in Table S2. Parameters were chosen such that the wild-type behaviour reported in Meyer et al. (2017) could be recapitulated. The outcome of the simulation in Fig. 7B was displayed using Paraview software (71).

We recently proposed a more detailed model of the ATML1 regulatory network to study how giant cell specification and cell fate maintenance depends on VLCFA (13), which is still a stochastic and cell-autonomous model. Here, however, for the sake of simplicity, and the intention of using a minimal, stochastic and cell-autonomous phenomenological model, the former ATML1 model was used (12).

### Randomizations of the outcomes from the numerical simulations

To assess the randomness of the giant cell pattern in the numerical simulations (Fig. 7), the same method was employed as that used for the experimental images. Although randomizations of the tissues were performed similarly (see the “Randomization of the experimental images” section above), a Python script was developed to display the output of the simulation as a multi-labeled image, where each cell was colored with a different label. These images could therefore be randomized using the dmSET method (36,37). To compare the simulated giant cell pattern (Fig. 7B) to the giant cell pattern found in the experimental mature sepals (Fig. 5B), the output image was cropped using the maximal rectangle in the tissue, and giant cells were also defined by a size threshold, ensuring that all cells with a ploidy of 16C or higher were categorized as giant cells (Fig. 7C). The few 8C cells that exceeded this threshold were also considered as giant cells.

To study the change in the giant cell spatial pattern over time (Fig. 7F), the same simulations were used, but only the first-arising giant cells (i.e., cells that stopped dividing after time t = 55 of the simulations for being committed to endoreduplicate) were studied. The same method was used to assess the randomness of the cellular patterns on these giant cells both at time t = 55 and time t = 135. Here, instead of cropping the image, the whole tissue was randomized (using the dmSET method), including the cells at the edges, such that exactly the same giant cells were considered at both time points. Examples of randomizations are shown in Fig. S20A–B. The analysis was performed over five simulation replicates, with different cellular random initial conditions.

Related “Segmentation” and “Randomization” images appearing in Fig. 7 and Fig. S20 were produced with a Python script using the multi-labeled images generated by dmSET.

## Acknowledgements

The authors would like to thank two reviewers for their insights, which helped improve the manuscript. We thank Nicholas J. Russell, John Chandler, Josep Mercadal, Elise Laruelle and Philippe Andrey for critical comments on the manuscript. We thank Stephen Starkman, Emily Phung, and Isabel Delo for assistance with segmentation, and Violeta Gibelli for assistance with cell classification corrections. We thank Richard Smith and Soeren Strauss for their help in using MorphoGraphX; we are grateful that Richard Smith created a process to generate 2D images from meshes. We also thank Philippe Andrey for his insights on image analysis and statistics and Elise Laruelle for her help in using the dmSET randomization code, as well as the Cornell Statistics Consulting Unit, specifically May Boggess, for her help with statistics and coding.

## Author Contributions

Conception and design of experiments: FC, GW, PFJ, AHKR. Imaging: FC.

Image analysis: FC, GW, XW. Data analysis: GW, FC, CB, XW.

Data curation: GW, FC, AHKR, PFJ. Analysis of pattern randomness: GW. Modeling: GW, PFJ.

Interpretation of results: GW, FC, AHKR, PFJ.

Writing - Original draft preparation: FC, GW, PFJ, AHKR. Writing - Review and editing: GW, FC, CB, XW,, AHKR, PFJ. Visualization: GW, FC.

## Data availability

Microscopy data, MorphoGraphX meshes for the images, as well as material used for data analysis and modeling are available at Open Science Framework (osf.io), DOI:10.17605/OSF.IO/RFCWS.

## References

1. Zuch DT, Doyle SM, Majda M, Smith RS, Robert S, Torii KU. Cell biology of the leaf epidermis: Fate specification, morphogenesis, and coordination. Plant Cell. 2022;34(1):209– 27.

2. Hülskamp M. Trichomes. Curr Biol. 2019 Apr 22;29(8):R273–4.

3. Katagiri Y, Hasegawa J, Fujikura U, Hoshino R, Matsunaga S, Tsukaya H. The coordination of ploidy and cell size differs between cell layers in leaves. Development. 2016 Apr 1;143(7):1120–5.

4. Melaragno JE, Mehrotra B, Coleman AW. Relationship between endopolyploidy and cell size in epidermal tissue of Arabidopsis. Plant Cell. 1993;5(11):1661–8.

5. Bergmann DC, Sack FD. Stomatal development. Annu Rev Plant Biol. 2007;58:163–81.

6. Horst RJ, Fujita H, Lee JS, Rychel AL, Garrick JM, Kawaguchi M, et al. Molecular Framework of a Regulatory Circuit Initiating Two-Dimensional Spatial Patterning of Stomatal Lineage. PLoS Genet. 2015;11(7):e1005374.

7. Torii KU. Stomatal development in the context of epidermal tissues. Ann Bot. 2021;128(2):137–48.

8. Hülskamp M. Plant trichomes: A model for cell differentiation. Nat Rev Mol Cell Biol. 2004;5(6):471–80.

9. Balkunde R, Pesch M, Hülskamp M. Trichome patterning in Arabidopsis thaliana from genetic to molecular models. Curr Top Dev Biol. 2010;91:299–321.

10. Roeder AHK, Chickarmane V, Cunha A, Obara B, Manjunath BS, Meyerowitz EM. Variability in the Control of Cell Division Underlies Sepal Epidermal Patterning in Arabidopsis thaliana. PLOS Biol. 2010;8(5):e1000367.

11. Roeder AHK, Cunha A, Ohno CK, Meyerowitz EM. Cell cycle regulates cell type in the Arabidopsis sepal. Development. 2012 Dec 1;139(23):4416–27.

12. Meyer HM, Teles J, Formosa-Jordan P, Refahi Y, San-Bento R, Ingram G, et al. Fluctuations of the transcription factor ATML1 generate the pattern of giant cells in the Arabidopsis sepal. Elife. 2017;6:e19131.

13. Vadde BVL, Russell NJ, Bagde SR, Askey B, Saint-Antoine M, Brownfield B, et al. The transcription factor ATML1 maintains giant cell identity by inducing synthesis of its own long-chain fatty acid-containing ligands. bioRxiv. 2024.

14. Gifford ML, Dean S, Ingram GC. The Arabidopsis ACR4 gene plays a role in cell layer organisation during ovule integument and sepal margin development. Development. 2003;130(18):4249–58.

15. Tanaka H, Watanabe M, Watanabe D, Tanaka T, Machida C, Machida Y. ACR4, a putative receptor kinase gene of Arabidopsis thaliana, that is expressed in the outer cell layers of embryos and plants, is involved in proper embryogenesis. Plant Cell Physiol. 2002;43(4):419– 28.

16. De Smet I, Vassileva V, De Rybel B, Levesque MP, Grunewald W, Van Damme D, et al. Receptor-Like Kinase ACR4 Restricts Formative Cell Divisions in the Arabidopsis Root. Science. 2008;322(5901):594–7.

17. Churchman ML, Brown ML, Kato N, Kirik V, Hülskamp M, Inzé D, et al. SIAMESE, a plant-specific cell cycle regulator, controls endoreplication onset in Arabidopsis thaliana. Plant Cell. 2006;18(11):3145–57.

18. Kumar N, Harashima H, Kalve S, Bramsiepe J, Wang K, Sizani BL, et al. Functional Conservation in the SIAMESE-RELATED Family of Cyclin-Dependent Kinase Inhibitors in Land Plants. Plant Cell. 2015;27(11):3065–80.

19. Dubois M, Achon I, Brench RA, Polyn S, Tenorio Berrío R, Vercauteren I, et al. SIAMESE- RELATED1 imposes differentiation of stomatal lineage ground cells into pavement cells. Nat Plants. 2023;9(7):1143–53.

20. Schwarz EM, Roeder AH. Transcriptomic Effects of the Cell Cycle Regulator LGO in Arabidopsis Sepals. Front Plant Sci. 2016;7.

21. Wilkens A, Czerniawski P, Bednarek P, Libik-Konieczny M, Yamada K. ATML1 Regulates the Differentiation of ER Body–Containing Large Pavement Cells in Rosette Leaves of Brassicaceae Plants. Plant Cell Physiol. 2024;65(7):1160–72.

22. Galletti R, Johnson KL, Scofield S, San-Bento R, Watt AM, Murray JA, et al. DEFECTIVE KERNEL 1 promotes and maintains plant epidermal differentiation. Development. 2015;142(11):1978–83.

23. Dong J, Bergmann DC. Stomatal patterning and development. Curr Top Dev Biol. 2010;91:267–97.

24. Greese B, Hulskamp M, Fleck C. Quantification of variability in trichome patterns. Front Plant Sci. 2014;5:596.

25. Liu C, Li Y, Xu L, Li M, Wang J, Yan P, et al. Stomatal Arrangement Pattern: A New Direction to Explore Plant Adaptation and Evolution. Front Plant Sci. 2021;12:655255.

26. Zeng SM, Lo EKW, Hazelton BJ, Morales MF, Torii KU. Effective range of non-cell autonomous activator and inhibitor peptides specifying plant stomatal patterning. Development. 2020;147(17).

27. Kuan C, Yang SL, Ho CMK. Using quantitative methods to understand leaf epidermal development. Quant Plant Biol. 2022;3.

28. Kawade K, Tsukaya H. Probing the stochastic property of endoreduplication in cell size determination of Arabidopsis thaliana leaf epidermal tissue. PLOS One. 2017;12(9):e0185050.

29. Russell NJ, Belato PB, Oliver LS, Chakraborty A, Roeder AHK, Fox DT, et al. Spatial ploidy inference using quantitative imaging. bioRxiv. 2025.

30. Donnelly PM, Bonetta D, Tsukaya H, Dengler RE, Dengler NG. Cell Cycling and Cell Enlargement in Developing Leaves of Arabidopsis. Dev Biol. 1999;215(2):407–19.

31. Haber AH. Nonessentiality of concurrent cell divisions for degree of polarization of leaf growth I. studies with radiation-induced mitotic inhibition. Am J Bot. 1962;49:583–9.

32. Tang HB, Wang J, Wang L, Shang GD, Xu ZG, Mai YX, et al. Anisotropic cell growth at the leaf base promotes age-related changes in leaf shape in Arabidopsis thaliana. Plant Cell. 2023;35(5):1386–407.

33. Vadde BVL, Roeder AHK, Wellmer F. Can the French flag and reaction–diffusion models explain flower patterning? Celebrating the 50th anniversary of the French flag model. J Exp Bot. 2020;71(10):2886–97.

34. Balkunde R, Deneer A, Bechtel H, Zhang B, Herberth S, Pesch M, et al. Identification of the Trichome Patterning Core Network Using Data from Weak ttg1 Alleles to Constrain the Model Space. Cell Rep. 2020;33(11):108497.

35. Mercadal J, Ferreira-Guerra M, Caño-Delgado AI, Ibañes M. A diffusion-driven switch specifies rhizoid precursor cells in Marchantia polymorpha. bioRxiv. 2023.

36. Laruelle E, Spassky N, Genovesio A. Unraveling spatial cellular pattern by computational tissue shuffling. Commun Biol. 2020;3(1):605.

37. Laruelle E, Genovesio A. Non-convex cell epithelial modeling unveils cellular interactions. IEEE 19th Int Symp Biomed Imaging ISBI. 2022;1–5.

38. Carter R, Sánchez-Corrales YE, Hartley M, Grieneisen VA, Marée AFM. Pavement cells and the topology puzzle. Development. 2017;144(23):4386–97.

39. Hervieux N, Dumond M, Sapala A, Routier-Kierzkowska AL, Kierzkowski D, Roeder AHK, et al. A Mechanical Feedback Restricts Sepal Growth and Shape in Arabidopsis. Curr Biol. 2016;26(8):1019–28.

40. Grebe M. The patterning of epidermal hairs in Arabidopsis — updated. Curr Opin Plant Biol. 2012;15(1).

41. Summers HD, Wills JW, Rees P. Spatial statistics is a comprehensive tool for quantifying cell neighbor relationships and biological processes via tissue image analysis. Cell Rep Methods. 2022;2(11).

42. Kumar G, Pandurengan RK, Parra ER, Kannan K, Haymaker C. Frontiers | Spatial modelling of the tumor microenvironment from multiplex immunofluorescence images: methods and applications. Front Immunol. 2023;14.

43. Sandler O, Mizrahi SP, Weiss N, Agam O, Simon I, Balaban NQ, et al. Lineage correlations of single cell division time as a probe of cell-cycle dynamics. Nature. 2015;519(7544).

44. Kobayashi H. Variations of endoreduplication and its potential contribution to endosperm development in rice. Plant Prod Sci. 2019;22(2):227–41.

45. Robinson DO, Coate JE, Singh A, Hong L, Bush M, Doyle JJ, et al. Ploidy and Size at Multiple Scales in the Arabidopsis Sepal. Plant Cell. 2018;30(10):2308–29.

46. Horiguchi G, Tsukaya H. Organ size regulation in plants: insights from compensation. Front Plant Sci. 2011;2:24.

47. Franks RG, Liu Z, Fischer RL, Liu Z, Fischer RL. SEUSS and LEUNIG regulate cell proliferation, vascular development and organ polarity in Arabidopsis petals. Planta. 2006;224(4).

48. Monniaux V. How to Evolve a Perianth: A Review of Cadastral Mechanisms for Perianth Identity. Front Plant Sci. 2018;9.

49. Le Gloanec C, Gómez-Felipe A, Alimchandani V, Branchini E, Bauer A, Routier-Kierzkowska AL, et al. Modulation of cell differentiation and growth underlies the shift from bud protection to light capture in cauline leaves. Plant Physiol. 2024;196(2):1214–30.

50. Johnson KL, Degnan KA, Ross Walker J, Ingram GC. AtDEK1 is essential for specification of embryonic epidermal cell fate. Plant J. 2005;44(1):114–27.

51. Abe M, Katsumata H, Komeda Y, Takahashi T. Regulation of shoot epidermal cell differentiation by a pair of homeodomain proteins in Arabidopsis. Development. 2003;130(4).

52. Ogawa E, Yamada Y, Sezaki N, Kosaka S, Kondo H, Kamata N, et al. ATML1 and PDF2 Play a Redundant and Essential Role in Arabidopsis Embryo Development. Plant Cell Physiol. 2015;56(6).

53. Li P, Elowitz MB, Klein A, Treutlein B. Communication codes in developmental signaling pathways. Development. 2019;146(12).

54. Clough SJ, Bent AF. Floral dip: a simplified method for Agrobacterium-mediated transformation of Arabidopsis thaliana. Plant J. 1998;16(6):735–43.

55. Sessions A, Burke E, Presting G, Aux G, McElver J, Patton D, et al. A high-throughput Arabidopsis reverse genetics system. Plant Cell. 2002;14(12):2985–94.

56. Rita San-Bento, Etienne Farcot, Roberta Galletti, Audrey Creff, Gwyneth Ingram. Epidermal identity is maintained by cell–cell communication via a universally active feedback loop in Arabidopsis thaliana. Plant J. 2014;77(1):46–58.

57. Smyth DR, Bowman JL, Meyerowitz EM. Early flower development in Arabidopsis. Plant Cell. 1990;2(8):755–67.

58. Barbier de Reuille P, Routier-Kierzkowska AL, Kierzkowski D, Bassel GW, Schüpbach T, Tauriello G, et al. MorphoGraphX: A platform for quantifying morphogenesis in 4D. Elife. 2015;4:e05864.

59. Strauss S, Runions A, Lane B, Eschweiler D, Bajpai N, Trozzi N, et al. Using positional information to provide context for biological image analysis with MorphoGraphX 2.0. Elife. 2022;11.

60. Trozzi N, Lane B, Perruchoud A, Wang Y, Hörmayer L, Ansel M, et al. Puzzle cell shape emerges from the interaction of growth with mechanical constraints. bioRxiv. 2023.

61. MATLAB version: 9.7.0 (R2019b). Natick, Massachusetts, United States: The MathWorks, Inc.; 2019. Available from: https://www.mathworks.com

62. Classification Learner App (R2019b). Natick, Massachusetts, United States: The MathWorks, Inc.; 2019. Available from: https://www.mathworks.com

63. van Spoordonk R, Schneider R, Sampathkumar. Mechano-chemical regulation of complex cell shape formation: Epidermal pavement cells—A case study. Quant Plant Biol. 2023;4(:e5).

64. Hagberg AA, Swart PJ, Schult DA, Swart PJ. Exploring network structure, dynamics, and function using NetworkX. In: Proceedings of the 7th Python in Science Conference (SciPy2008). Gäel Varoquaux, Travis Vaught, and Jarrod Millman (Eds); 2008. p. 11–5.

65. Hunter JD. Matplotlib: A 2D Graphics Environment. Comput Sci Eng. 2007;9(3):90–5.

66. Waskom ML. seaborn: statistical data visualization. J Open Source Softw. 2021;6(60):3021.

67. Gillespie DT. The chemical Langevin equation. 2000;297(May 2013):297–306.

68. Kadiyala U, Sprinzak D, Monk NA, Taylor SE, Verd B, Sonnen KF, et al. From genes to patterns: five key dynamical systems concepts to decode developmental regulatory mechanisms. Development. 2025;152:dev204617.

69. Hamant O, Heisler MG, Jönsson H, Krupinski P, Uyttewaal M, Bokov P, et al. Developmental Patterning by Mechanical Signals in Arabidopsis. Science. 2008;322(5908):1650–5.

70. Carrillo O, Ibanes M, Garcia-Ojalvo J, Casademunt J, Sancho JM. Intrinsic noise-induced phase transitions: beyond the noise interpretation. Phys Rev E Stat Nonlin Soft Matter Phys. 2003;67(4 Pt 2):046110.

71. Ahrens J, Geveci B, Law C. ParaView: An End-User Tool for Large Data Visualization. In: In Visualization Handbook. Elsevier. 2005.

