## Supporting Figures for "A common pathway controls cell size in the sepal and leaf epidermis leading to a non-random pattern of giant cells"

### Supplementary Figures

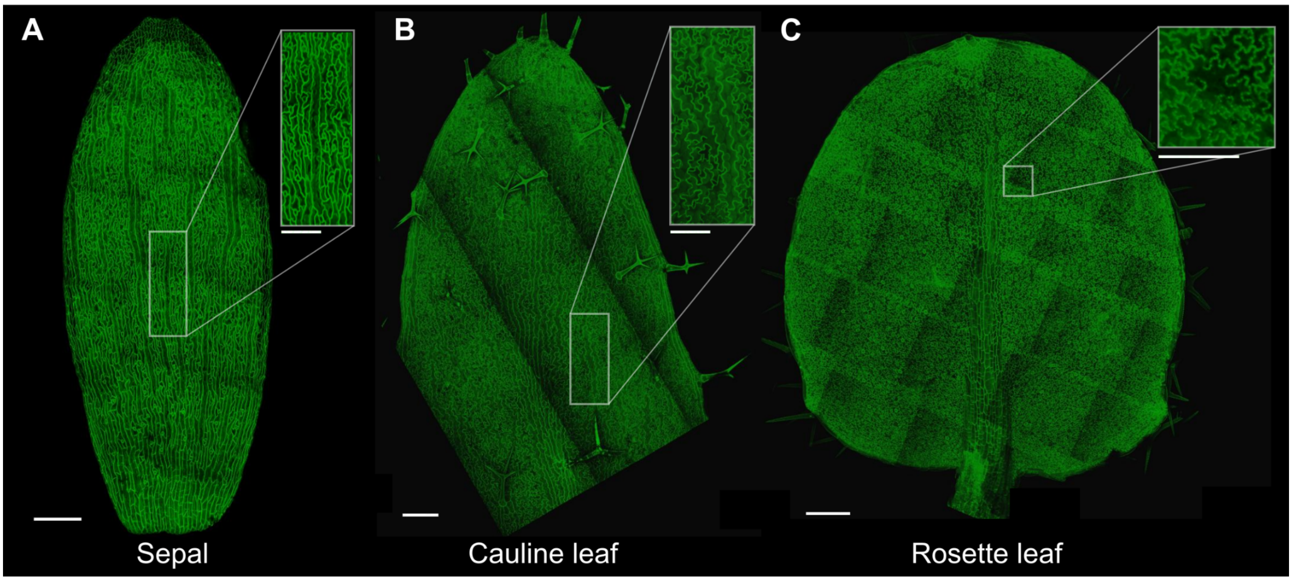


**Fig. S1. Cauline leaves have elongated giant cells similar to sepals.** (**A**) Abaxial side of a wild-type mature sepal expressing a cell membrane marker (*p35S::mCitrine-RCI2A*). (**B**) Tip section of the abaxial side of a wild-type cauline leaf expressing a cell membrane marker (*p35S::mCitrine-RCI2A*). (**C**) A developing abaxial side of a wild-type rosette leaf 1 or 2 at 8 dpg expressing a cell membrane marker (*p35S::mCitrine-RCI2A*). Scale bars associated with the overview images (bottom) represent 200 µm, and scale bars associated with the magnified images represent 100 µm. The underlying data for this figure can be found at Open Science Framework (osf.io), DOI:10.17605/OSF.IO/RFCWS.

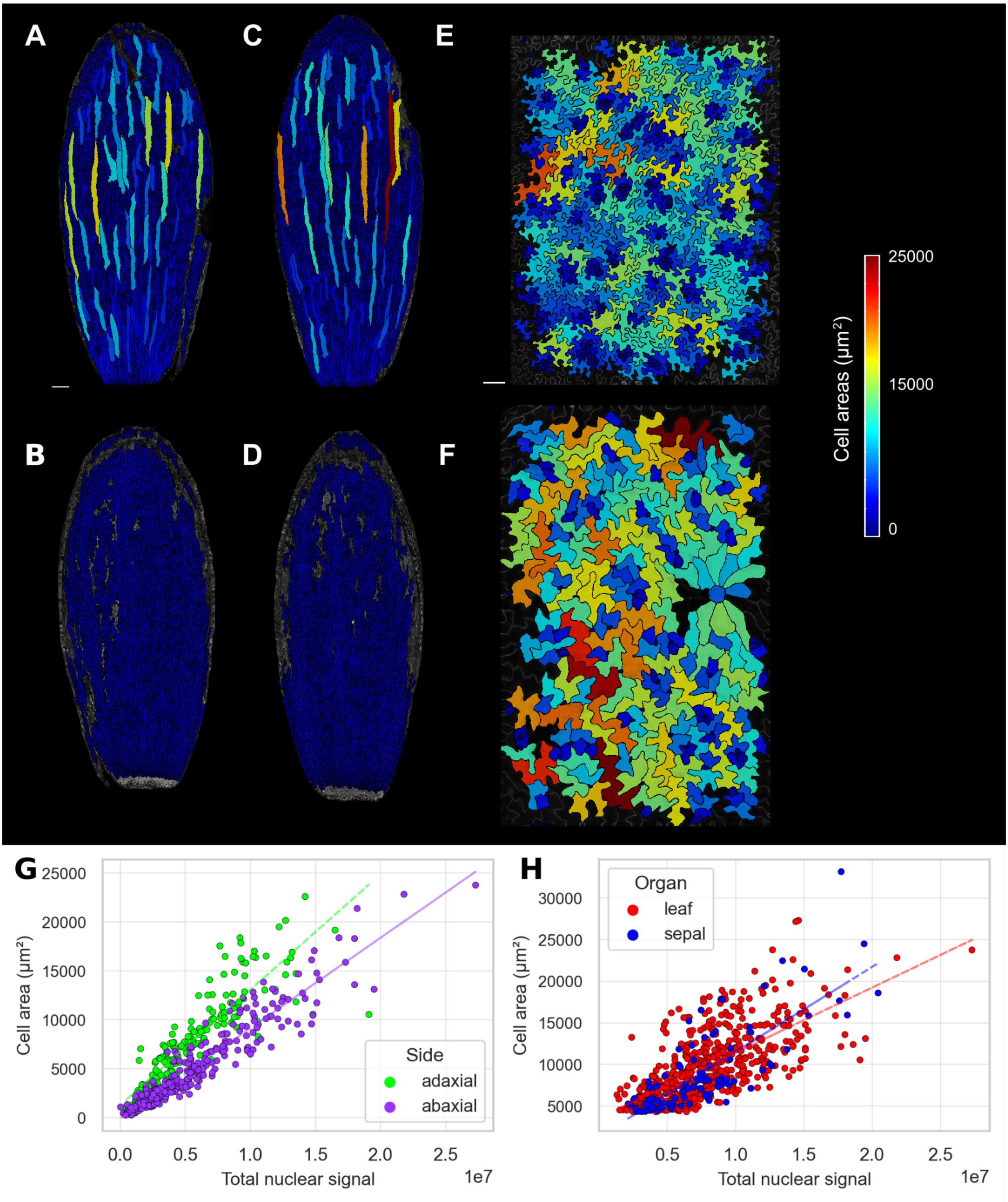


**Fig. S2. Replicates of abaxial and adaxial cell-size distribution in the wild-type leaf and sepal epidermis; cell size correlates with DNA content.** Cell area heat maps in µm$^2$ of (**A, C**) abaxial surfaces of wild-type sepals, (**B, D**) adaxial surfaces of wild-type sepals, (**E**) abaxial surface of 25-dpg wild-type leaf 1 or 2 (cell density: 284 cells mm$^{-2}$) and (**F**) adaxial surface of 25-dpg wild-type leaf 1 or 2 (cell density: 177 cells mm$^{-2}$). Scale bars represent 100 µm. (**G**) Abaxial and abaxial side of 25-dpg leaf cell area versus DNA content (one of two replicates) as measured by H2B-TFP total nuclear fluorescence, with $R^2 = 0.91$ for the abaxial side and $R^2 = 0.79$ for the adaxial side. Associated with Fig. 2. (**H**) Cell area of the largest cells (area > 4,308 µm$^2$) versus DNA content as measured by H2B-TFP total nuclear fluorescence in both the abaxial and adaxial side of the 25-dpg leaf (red) and in the abaxial side of the adult sepal (blue). The underlying data for this figure can be found at Open Science Framework (osf.io), DOI:10.17605/OSF.IO/RFCWS.

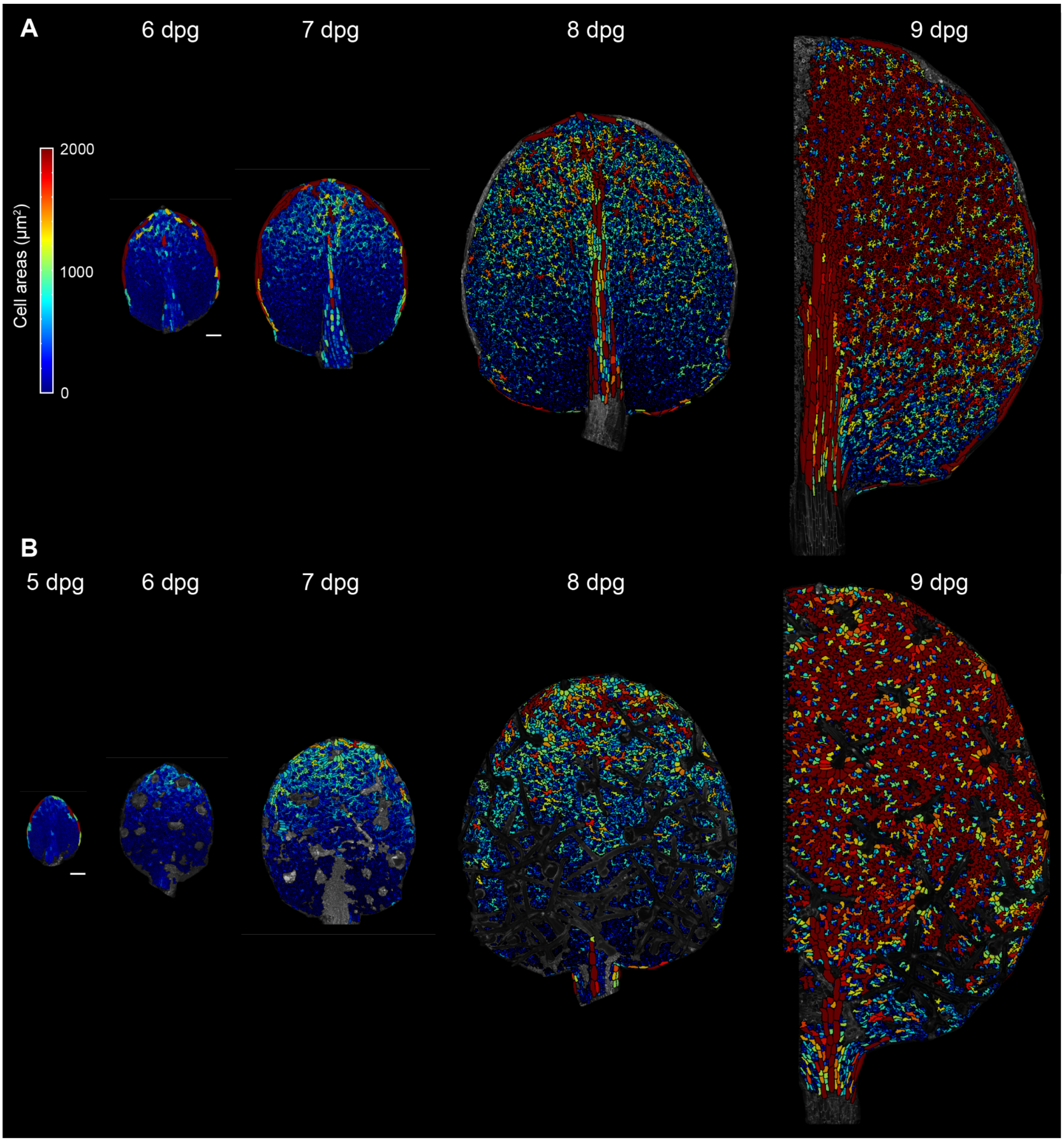


**Fig. S3. Cell-size patterning occurs as a basipetal wave simultaneously in the adaxial and abaxial sides of the leaf.** (**A**–**B**) Cell area heat maps in µm$^2$ for wild-type leaf 1 or 2 leaves at 5–9 dpg on (**A**) the abaxial side of the leaf and on (**B**) the adaxial side of the same leaf. Scale bar represents 100 µm. Leaves are to scale and have the same heat map color range. Associated with Fig. 3. The underlying data for this figure can be found at Open Science Framework (osf.io), DOI:10.17605/OSF.IO/RFCWS.

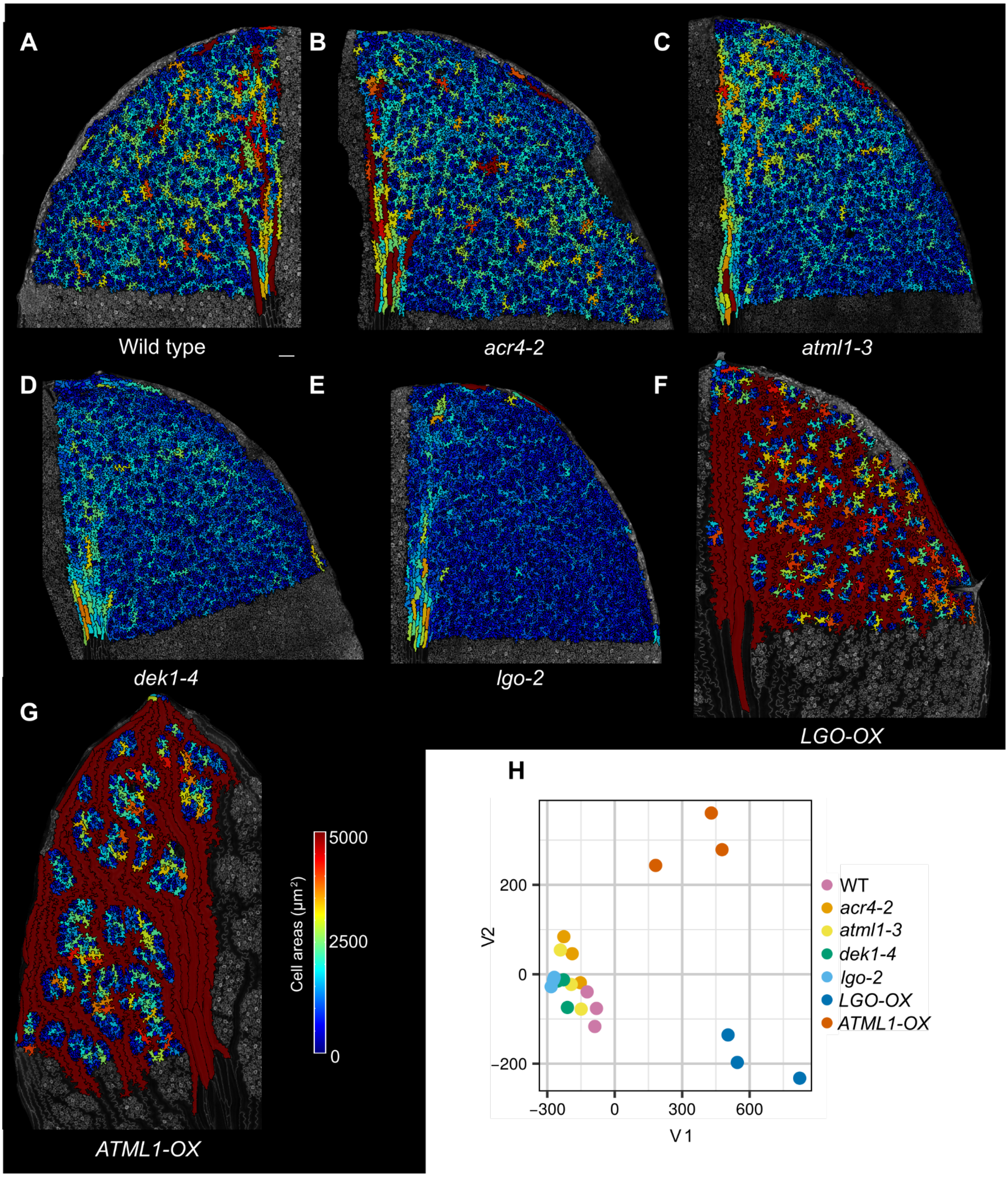


**Fig. S4. The sepal giant cell specification pathway also patterns cell size in 9-dpg leaves, replicate 2.** Cell area heat maps in µm$^2$ of the upper abaxial quadrant of leaf 1 or 2 at 9 dpg for the genotypes (**A**) wild type, (**B**) *acr4-2*, (**C**) *atml1-3*, (**D**) *dek1-4*, (**E**) *lgo-2*, (**F**) *LGO-OX* (*pATML1::LGO*) and (**G**) *ATML1-OX* (*pPDF1::ATML1*). Scale bar represents 100 µm. Second replicate associated with Fig. 4. (**H**) 2D Wasserstein distance plot for 9-dpg replicates. Cell area heat maps of other replicates are shown in Fig. 4 and Fig. S5. The Wasserstein statistical tests among replicates are shown in Fig. S8. The underlying data for this figure can be found at Open Science Framework (osf.io), DOI:10.17605/OSF.IO/RFCWS.

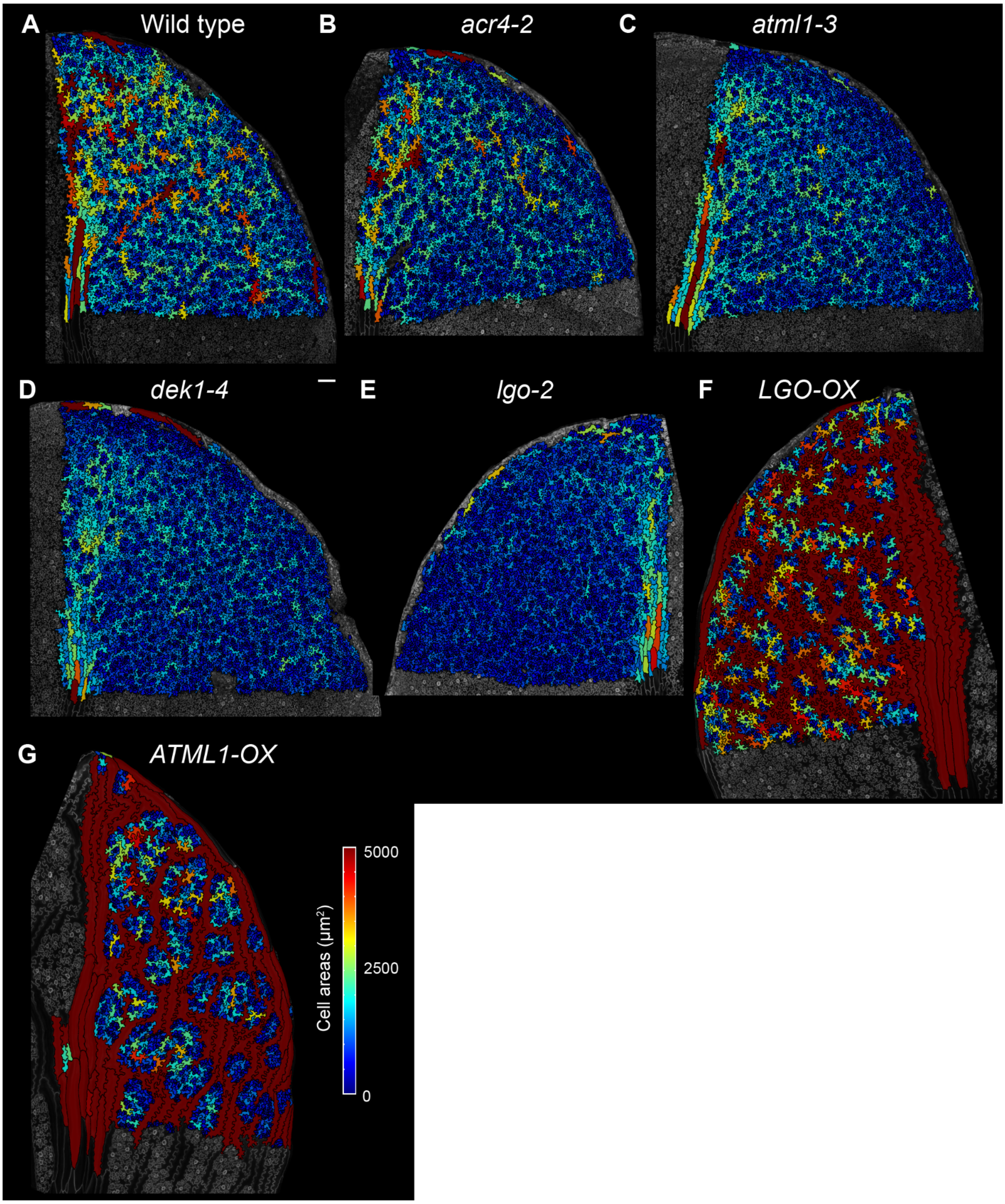


**Fig. S5. The sepal giant cell specification pathway also patterns cell size in 9-dpg leaves, replicate 3.** Cell area heat maps in µm$^2$ of the upper abaxial quadrant of leaf 1 or 2 at 9 dpg for the genotypes (**A**) wild type, (**B**) *acr4-2*, (**C**) *atml1-3*, (**D**) *dek1-4*, (**E**) *lgo-2*, (**F**) *LGO-OX* (*pATML1::LGO*) and **(G)** *ATML1-OX* (*pPDF1::ATML1*). Scale bar represents 100 µm. Third replicate associated with Fig. 4. The underlying data for this figure can be found at Open Science Framework (osf.io), DOI:10.17605/OSF.IO/RFCWS.

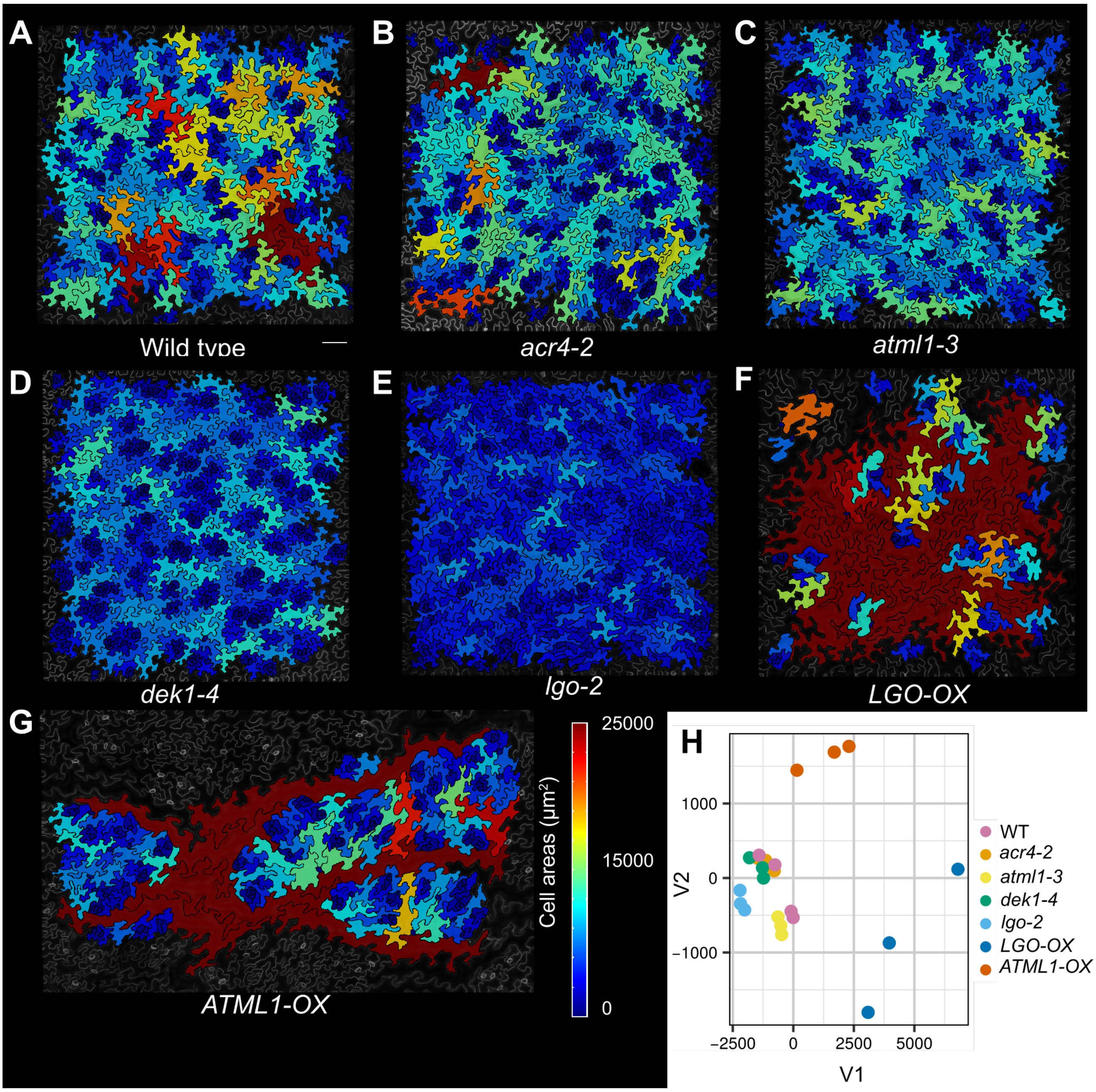


**Fig. S6. The sepal giant cell specification pathway also patterns cell size in 25-dpg mature leaves, replicate 2.** Cell area heat maps in μm$^2$ of a region approximately midway between midrib and margin and between tip and base on the abaxial side of leaf 1 or 2 at 25 dpg for the genotypes (**A**) wild type, (**B**) *acr4-2*, (**C**) *atml1-3*, (**D**) *dek1-4*, (**E**) *lgo-2*, (**F**) *LGO-OX* (*pATML1::LGO*) and (**G**) *ATML1-OX* (*pPDF1::ATML1*). Scale bar represents 100 μm. Second replicate associated with Fig. 4. (**H**) Wasserstein distances for 25-dpg replicates displayed as Euclidean distances embedded in 2D. Cell area heat maps of other replicates are shown in Figs. 4 and S7. Datasets from (E) and (F) are also used for an independent analysis in Trozzi et al., 2023. The underlying data for this figure can be found at Open Science Framework (osf.io), DOI:10.17605/OSF.IO/RFCWS.

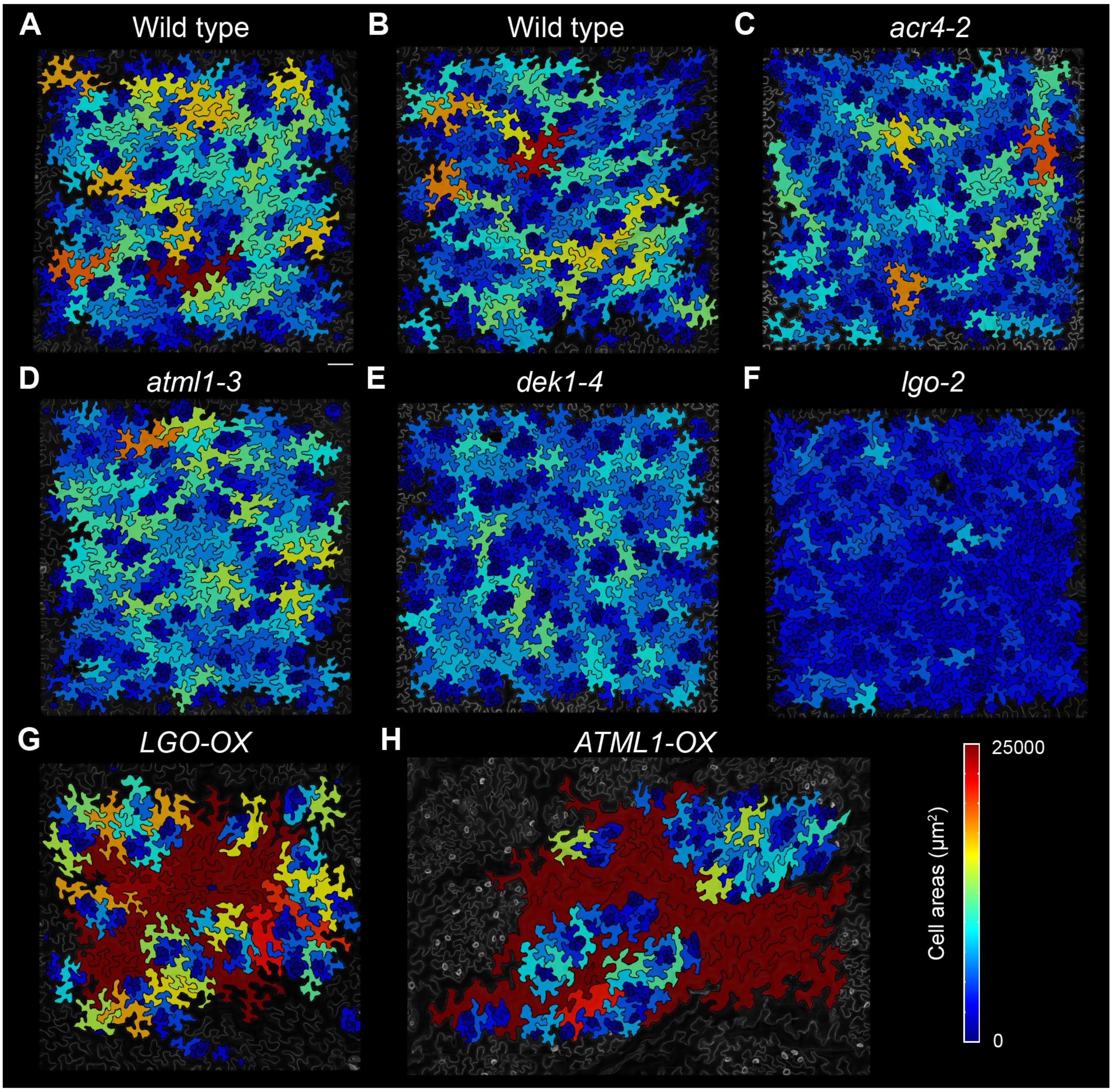


**Fig. S7. The sepal giant cell specification pathway also patterns cell size in 25-dpg mature leaves, replicate 3.** Cell area heat maps in µm$^2$ of an area approximately midway between midrib and margin and between tip and base on the abaxial side of leaf 1 or 2 at 25 dpg for the genotypes (**A–B**) wild type (two replicates), (**C**) *acr4-2*, (**D**) *atml1-3*, (**E**) *dek1-4*, (**F**) *lgo-2*, (**G**) *LGO-OX* (*pATML1::LGO*) and (**H**) *ATML1-OX* (*pPDF1::ATML1*). Scale bar is 100 µm. Third replicate associated with Fig. 4. Datasets from (A), (B), (F) and (G) are also used for an independent analysis in Trozzi et al., 2023. The underlying data for this figure can be found at Open Science Framework (osf.io), DOI:10.17605/OSF.IO/RFCWS.

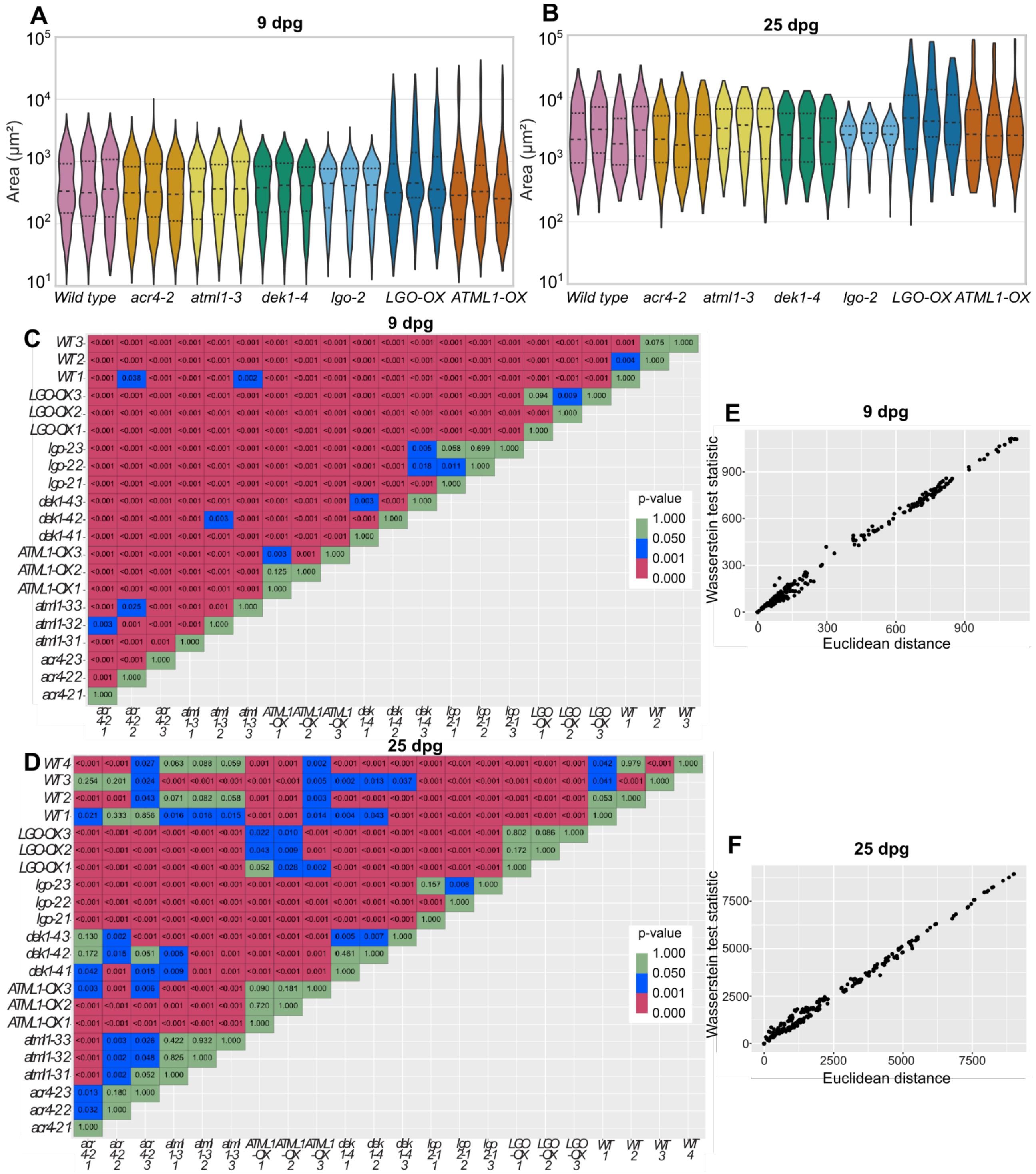


**Fig. S8. Statistical tests on the cell-size distributions and statistical tests in young and mature leaves.** (**A–B**) Violin plots of cell area densities on a $\log_{10}$ scale for individual replicates of (**A**) 9-dpg and (**B**) 25-dpg leaves. Stomata were removed in both (A) and (B). Associated with Fig. 4C and D. (**C–D**) p-values of the Wasserstein tests for all the replicate pair comparisons for (**C**) 9-dpg and (**D**) 25-dpg leaves. (**E–F**) Wasserstein test statistics plotted against their corresponding Euclidean distances for all (**E**) 9-dpg and (**F**) 25-dpg test pairs. Associated with Fig. 4. The underlying data for this figure can be found at Open Science Framework (osf.io), DOI:10.17605/OSF.IO/RFCWS.

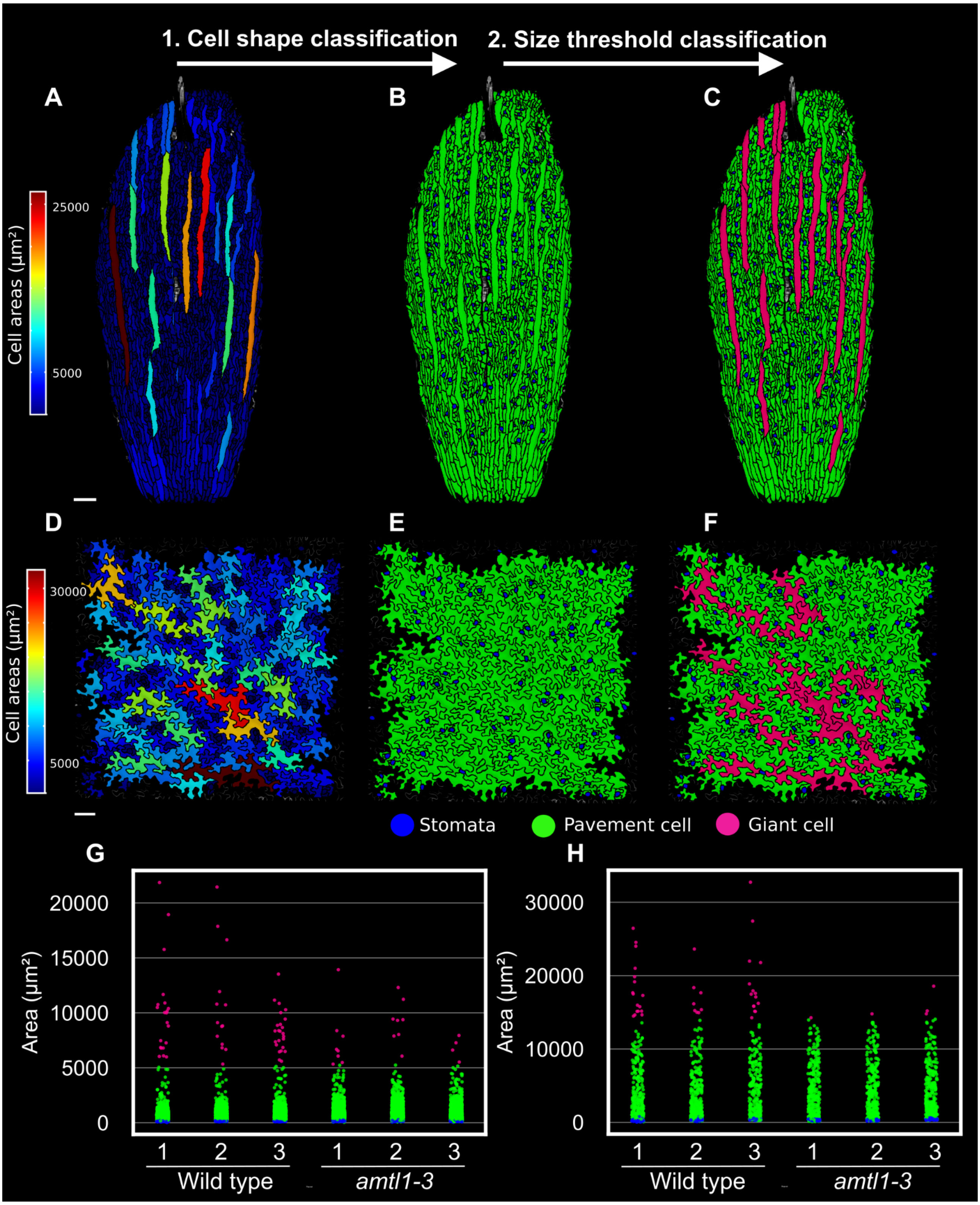


**Fig. S9. Classification of cell types in the leaf and the sepal**. (**A–F**) Steps for cell-type classification, with a wild-type sepal (A–C) and a wild-type leaf 25 dpg (D–F) as examples. First, cellular shape features were computed from the segmented meshes in MorphoGraphX (**A, D**) to train a classification algorithm (SVM quadratic) and predict the stomata (blue) and the pavement (green) cell types (**B, E**). Heat map colors represent the cell area in (A,D). Second, to classify giant cells (**C, F**), a size threshold was established based on observations in *atml1-3* sepals, such as a small fraction of pavement cells (0.7% on average) are giant cells in *atml1-3* mutants (see G, H). Heat map colors represent cell type in (B, C, D, E). (**G, H**) Cell size in three wild-type replicates and the three *atml1-3* replicates that were used to construct the threshold in leaf 25 dpg (G) and in the sepal (H). Points are colored according to the cell-type classification. Cells exceeding the defined size threshold (see Materials and Methods) are defined as giant cells (magenta). Associated with Fig. S10. The code and data associated with this figure can be found at Open Science Framework (osf.io), DOI:10.17605/OSF.IO/RFCWS.

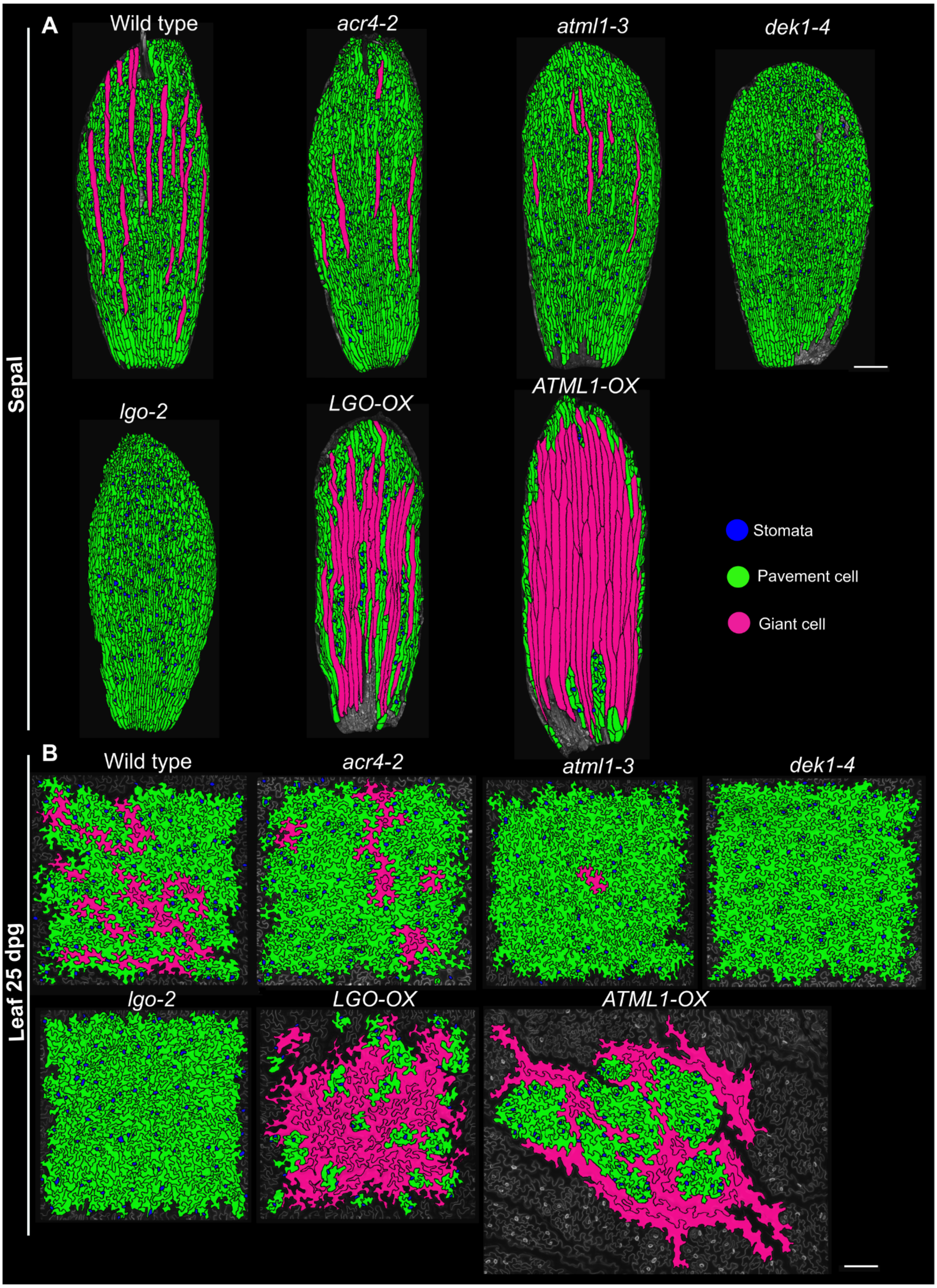


**Fig. S10. Cell type classification in the leaf and sepal**. (**A–B**) Segmented meshes of one replicate for each genotype after cell type classification in the mature sepal (**A**) and the mature leaf (**B**). Cells are colored with their corresponding cell type: pavement cells (in green), stomata (in blue) and giant cells (in magenta). Stomata and pavement cells were first classified using a trained classification algorithm based on cell shape features. Giant cells were defined as the largest cells, using a size threshold based on *atml1-3* mutants (see Materials and Methods). Scale bars represent 200 µm. See also Figs. S9 and S11. The code and data associated with this figure can be found at Open Science Framework (osf.io), DOI:10.17605/OSF.IO/RFCWS.

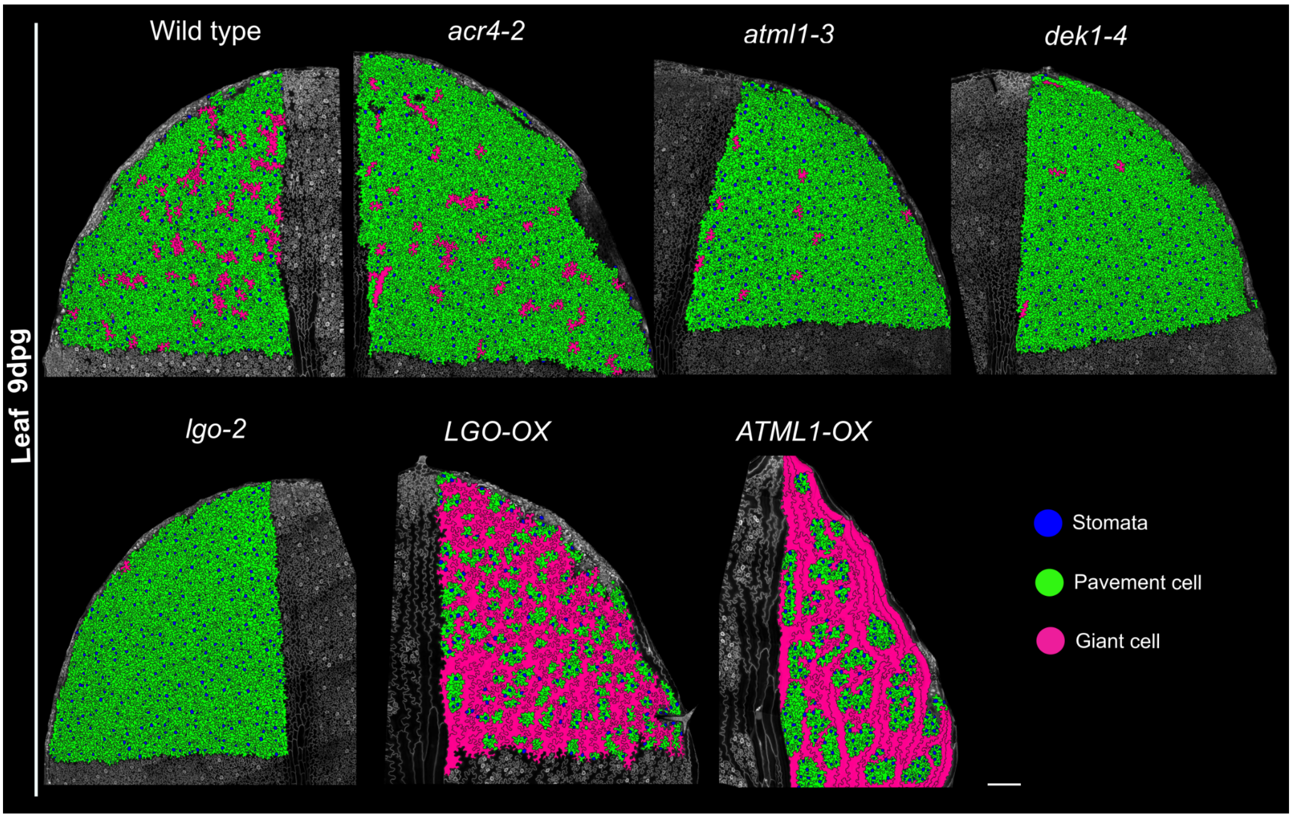


**Fig. S11. Cell type classification in the 9-dpg leaves.** Meshes of one replicate for each genotype in the 9-dpg leaf sections. Cells are colored with their corresponding cell type: pavement cells (in green), stomata (in blue) and giant cells (in magenta). Stomata and pavement cells were first classified using a trained SVM classification algorithm based on cell shape features. Because meristemoids and stomata are difficult to distinguish at this stage, both were classified as stomata. Giant cells were defined by a cell-size threshold based on *atml1-3* mutants (see Materials and Methods and Fig. S9). Scale bar represents 200 µm. Associated with Fig. S10. The code and data associated with this figure can be found at Open Science Framework (osf.io), DOI:10.17605/OSF.IO/RFCWS.

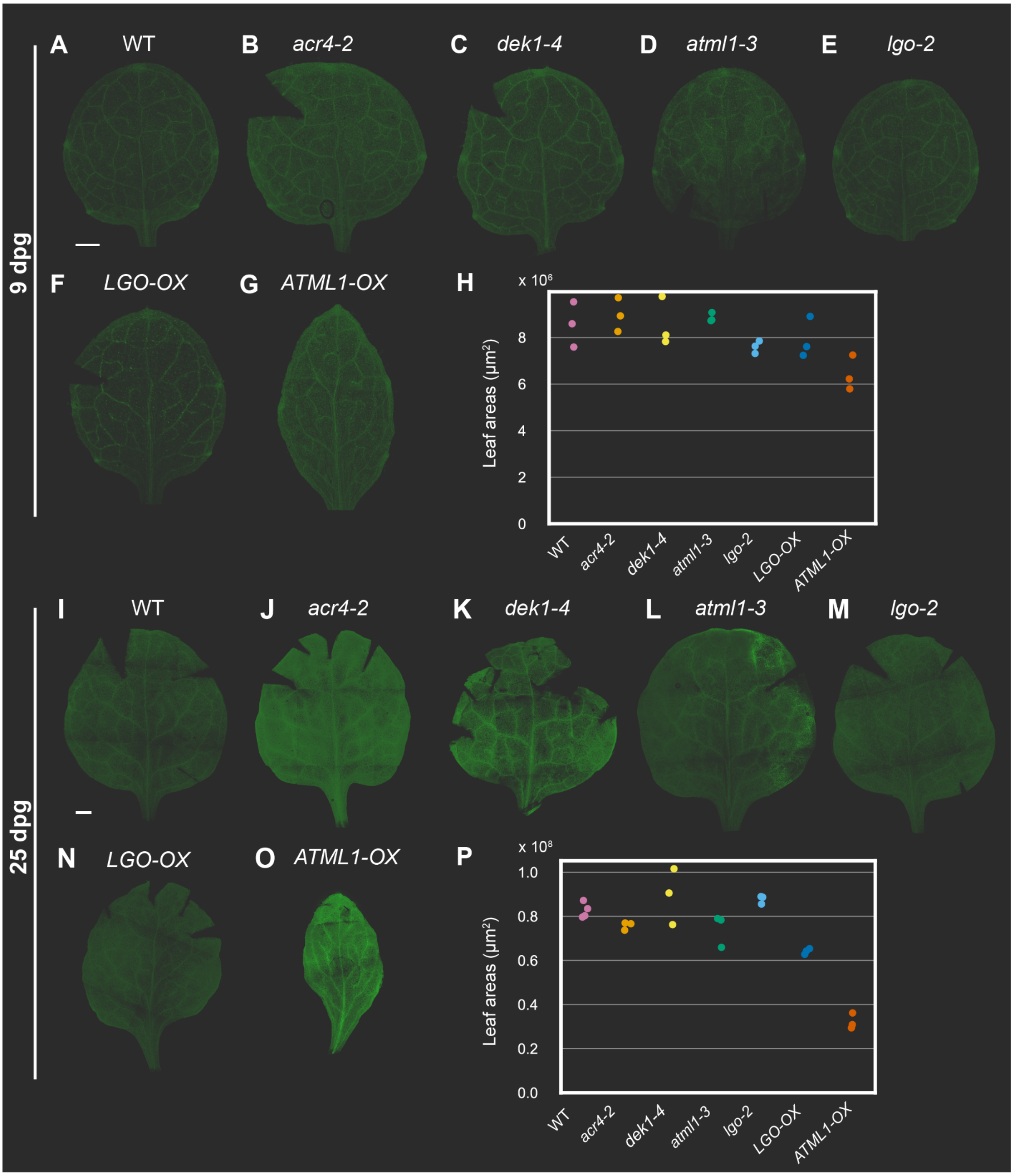


**Fig. S12. Cell-size patterning has little effect on leaf size, except in *ATML1-OX* and *LGO-OX*** (**A–G**) Images of 9 dpg full leaves with *p35S::mCitrine-RCI2A* used for segmenting the upper quadrant (Fig. 4). One replicate of each genotype is shown. Leaves of similar sizes were chosen to developmentally stage match as closely as possible. (**H**) Leaf areas of three replicates of 9-dpg leaves for each genotype. (**I–O**) Images of 25 dpg full leaves with *p35S::mCitrine-RCI2A* used for imaging and segmenting the mature cells (Fig. 4). One replicate of each genotype is shown. (**P**) Leaf areas of three replicates of 25-dpg leaves for each genotype. Scale bar is 0.5 mm for (A–G) and 1 mm for (I–

O). The underlying data for this figure can be found at Open Science Framework (osf.io), DOI:10.17605/OSF.IO/RFCWS.

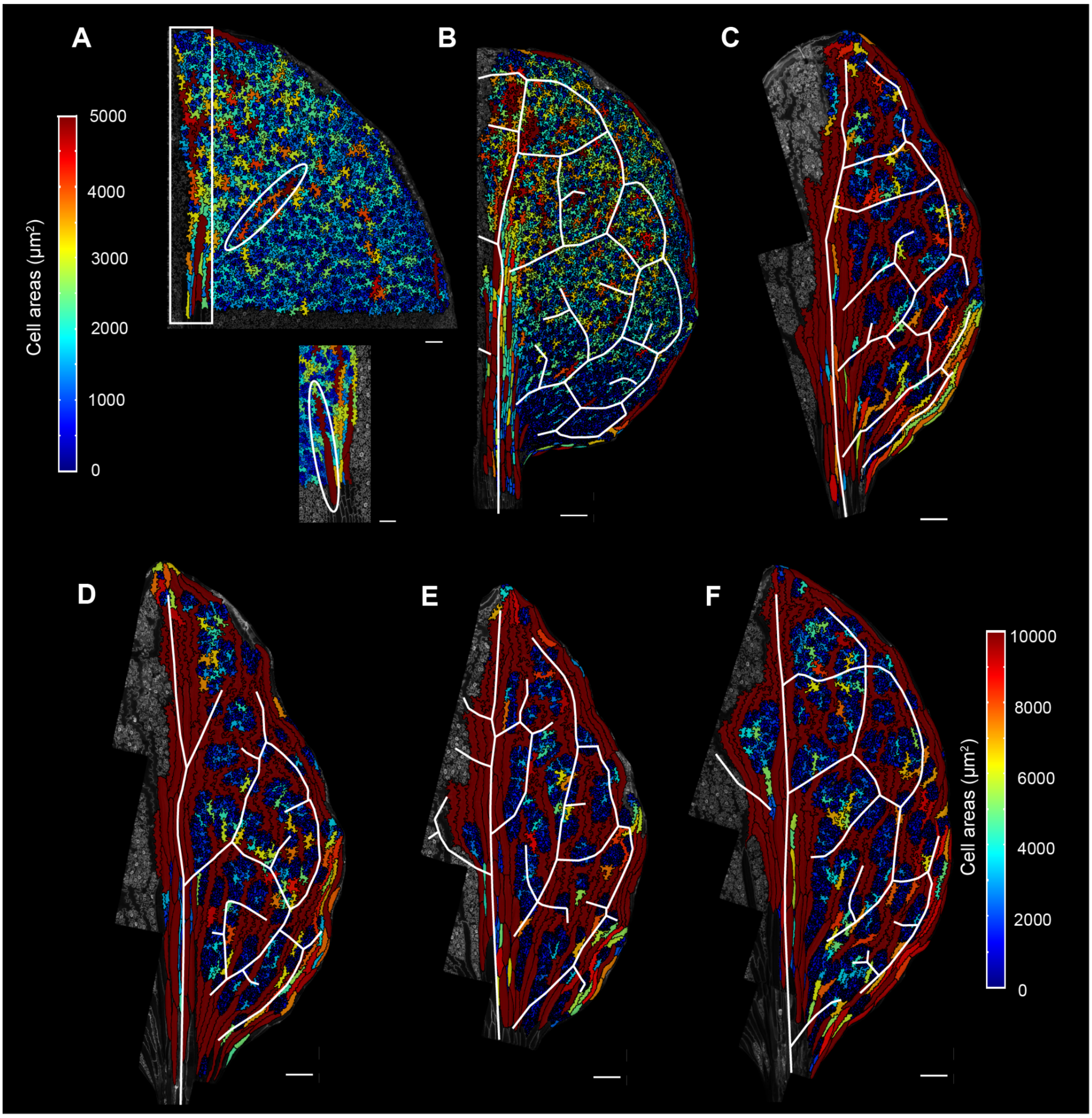


**Fig. S13. Giant cells are not preferentially positioned overlying the vasculature.** (**A**) Top: Cell area heat map of the abaxial upper quadrant of a 9-dpg wild-type leaf. Large cells overlying the midrib that extend up to the leaf tip are boxed in white. Large cells that align as if along an underlying vein are circled in white. Bottom: Cell area heat map of the abaxial midrib region of a 9-dpg wild-type leaf. A large pavement cell extending out from a large midrib cell as if following an underlying vein peeling off the midrib. (**B**) Cell area heat map of the abaxial side of half of a 9-dpg wild-type leaf with the underlying vasculature in white. The colored heat map is associated with the color bar in (A). (**C–F**) Cell area heat map of the abaxial sides of halves of *ATML1-OX* leaves with the underlying vasculature in white (four replicates; see Materials and Methods). Colored heat maps are associated with the color bar in (F). All scale bars represent 100 µm. The underlying data for this figure can be found at Open Science Framework (osf.io), DOI:10.17605/OSF.IO/RFCWS.

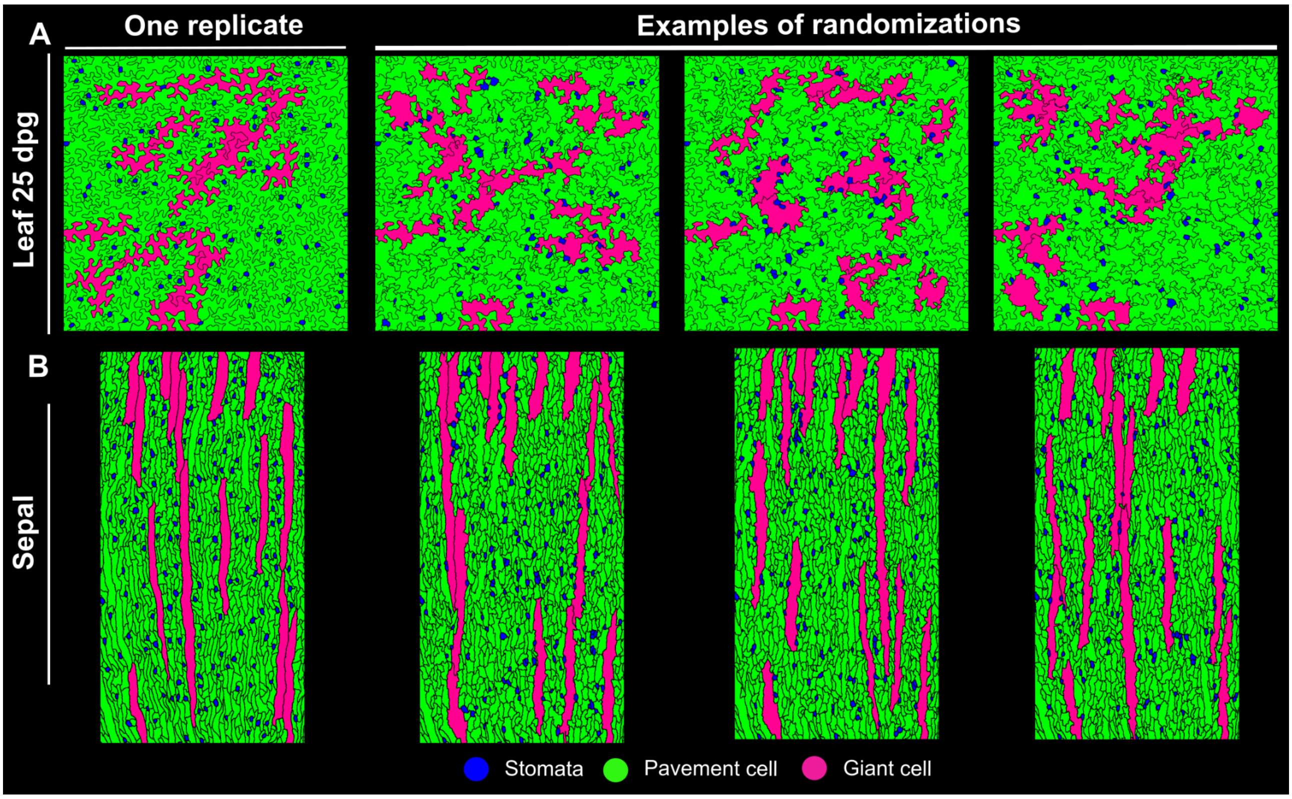


**Fig. S14. Examples of randomizations in the leaf and in the sepal.** (**A**) One replicate of the leaf after cell type classification and three corresponding randomized tissues as an example. (**B**) One replicate of the sepal after cell type classification and three corresponding randomized tissues as an example. Associated with Fig. 5. The code and data associated with this figure can be found at Open Science Framework (osf.io), DOI:10.17605/OSF.IO/RFCWS.

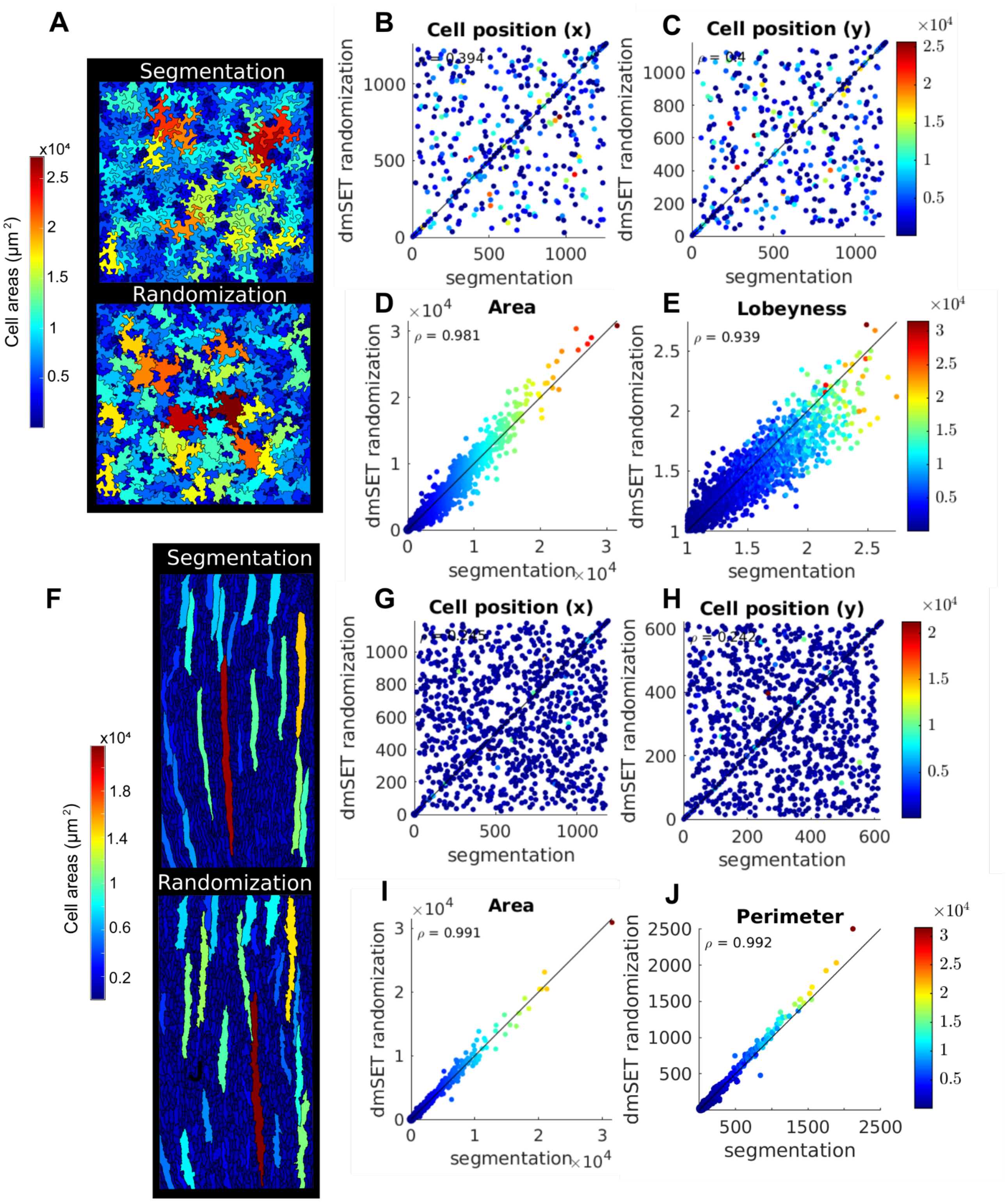


**Fig. S15. Conservation of cellular features between the segmented tissue and the corresponding randomized tissues.** (**A**) Example of a segmented tissue and a corresponding randomized tissue, generated using the dmSET method, in the leaf 25 dpg (the replicate used is the same as in Fig. 5B). (**B, C**) Comparison of the cell center coordinates (in µm) between the real tissue (segmentation) and a randomized tissue (dmSET randomization) shown in (A). (**D, E**) Comparison of cell shape features between all real tissues (segmentation) and one of their randomized tissues (dmSET randomization):

(D) cell area (in $\mu m^2$) and (E) cell lobeyness (defined as the perimeter of the cell divided by the perimeter of its convex hull). (**F**) Example of a segmented tissue and a corresponding randomized tissue, generated using the dmSET method, in the sepal (the replicate used is the same as in Fig. 5B). (**G, H**) Comparison of the cell center coordinates (in µm) between the real tissue (segmentation) and a randomized tissue (dmSET randomization) shown in (F). (**I, J**) cell shape features between all real tissues (segmentation) and one of their randomized tissues (dmSET randomization): cell area (I) and cell perimeter (J). The color bars associated with (B–C), (D–E), (G–H) and (I–J) represent the cell areas (in $\mu m^2$). Each dot represents one cell. Associated with Fig. 5. The code and data associated with this figure can be found at Open Science Framework (osf.io), DOI:10.17605/OSF.IO/RFCWS.

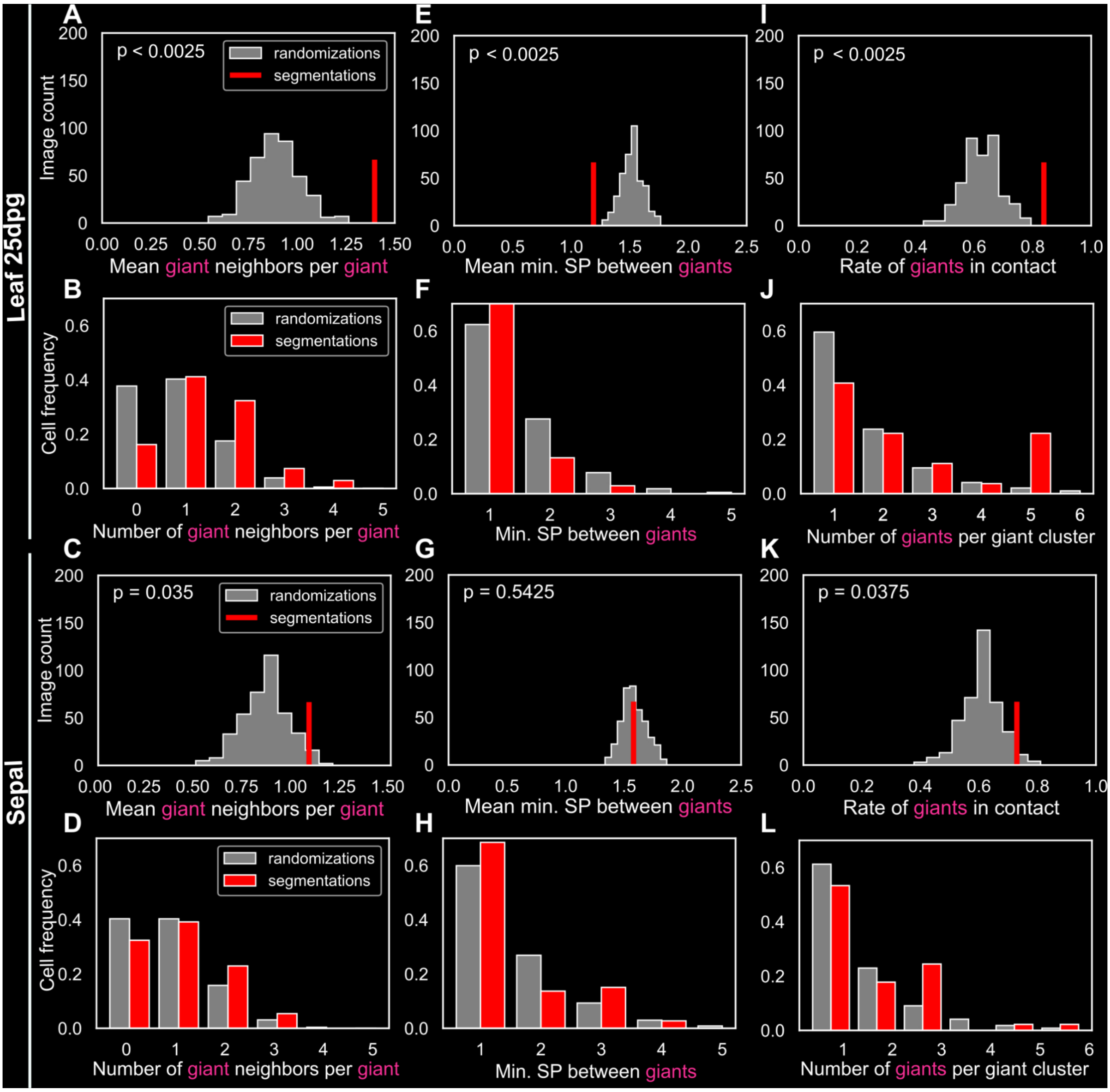


**Fig. S16. Different spatial observables were studied to statistically assess the spatial organization of giant cells.** Three measures were extracted from the cellular network of the segmentations (red) and statistically compared against the randomizations (gray). (**A–D**) The number of giant cell neighbors per giant, as a mean over giant cells (A, C) and as a distribution (B, D), as presented in Fig. 5C–D. (**E–H**) The minimum shortest path (min. SP) between two giant cells (in path length, 1 meaning that the pair of giant cells are in contact) as a mean over giant cells (E, G) and as a distribution (F, H). (**I–J**) The number of giant cells per giant cell cluster as a mean over giant cells (I, K) and as a distribution (J, L). The top panels (A, B, E, F, I, J) show the result of the pattern quantification in the wild-type leaf 25 dpg and the bottom panels (C, D, G, H, K, L) show the results in the wild-type sepal. Analyses were performed over six pooled replicates. Total number of giant cells counted (excluding giant cells at the image border) in the analysis: n = 68 (leaf, segmentations), n = 68 × 400 (leaf, randomizations), n = 74 (sepal, segmentations), n = 74 × 400 (sepal, randomizations). The code and data associated with this figure can be found at Open Science Framework (osf.io), DOI:10.17605/OSF.IO/RFCWS.

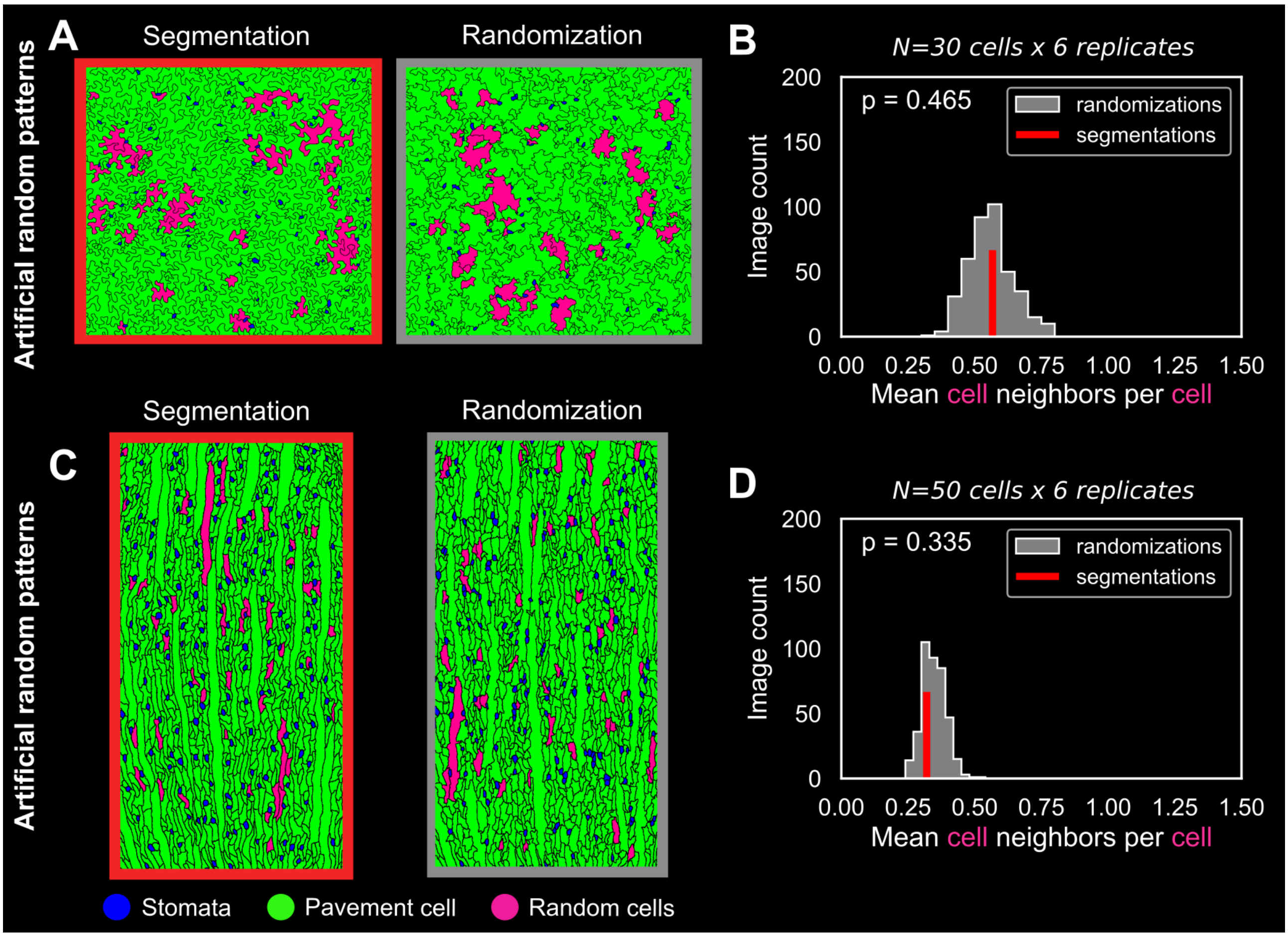


**Fig. S17: Assessment of the dmSET method on a random cell pattern.** (**A–B**) To decipher potential bias in our null model due to artifacts in randomized tissues, an artificial random cellular pattern was created in all leaf replicates by randomly selecting a population of pavement cells with an area larger than 2000 $\mu m^2$ in leaves. (A) Example of representative segmentation of a 25-dpg wild-type leaf (left) and one of its corresponding randomized tissue (right). Randomly selected cells are labeled in magenta. (B) Mean number of cell neighbors per cell within the randomly selected cells in the real tissues (segmentations) and in the randomized tissues (randomizations). The null hypothesis could not be rejected, showing that the artificial random pattern does not deviate significantly from randomness (see Materials and Methods). (**C–D**) Similar to (A–B) in the sepals by randomly selecting a population of pavement cells with an area larger than 100 $\mu m^2$. Total number of cells considered in the analysis: n = 10 × 6 (leaf, segmentations), n = 10 × 6 × 400 (leaf, randomizations), n = 30 × 6 (sepal, segmentations), n = 30 × 6 × 400 (sepal, randomizations). The code and data associated with this figure can be found at Open Science Framework (osf.io), DOI:10.17605/OSF.IO/RFCWS.

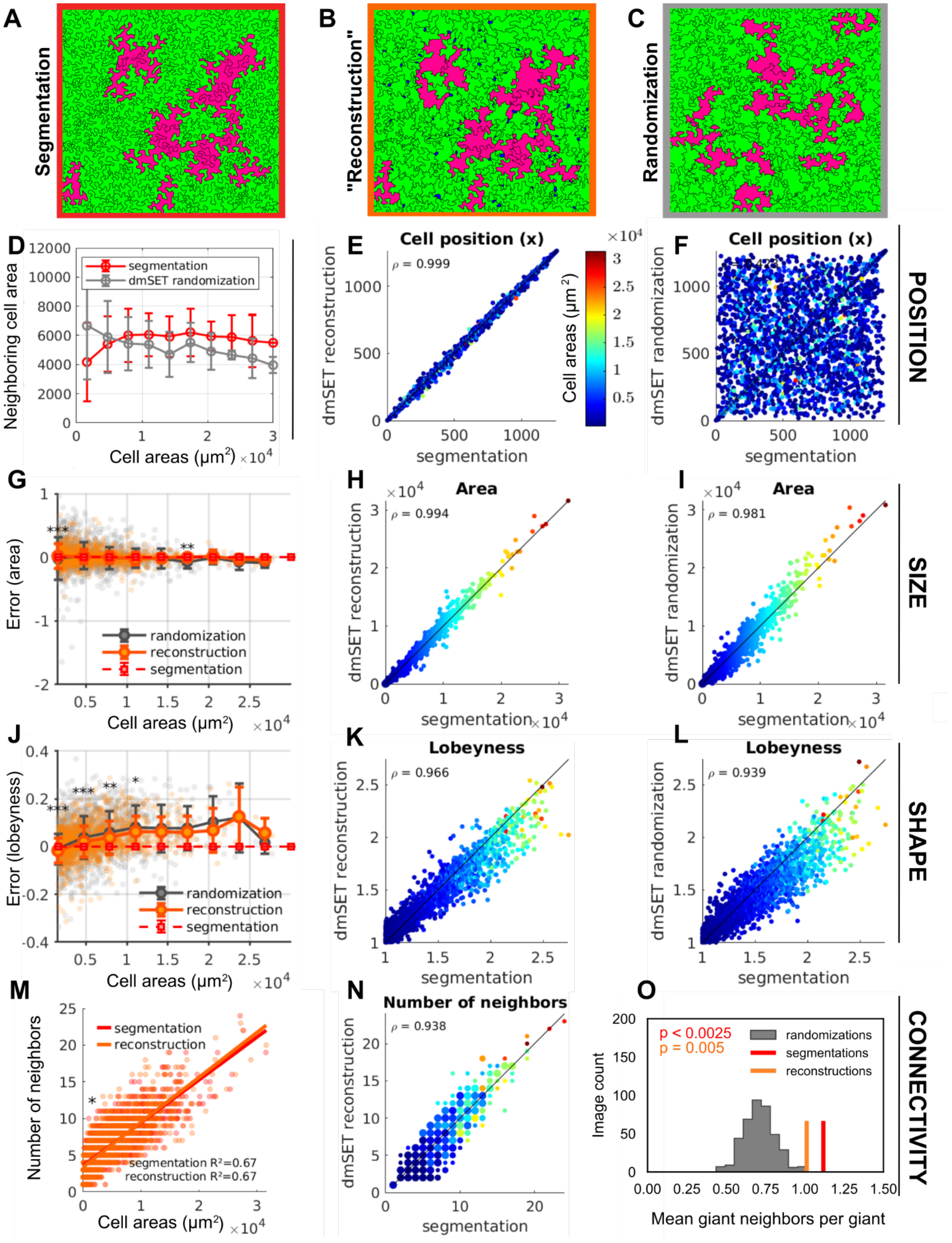


**Fig. S18: Reconstruction of the tissues using dmSET to investigate the effects of shape artifacts in randomized tissues.** (**A**–**C**) Example of (**A**) an original segmentation of a 25-dpg wild-type leaf, (**B**) a reconstruction, and (**C**) one randomized tissue. All leaf replicates were reconstructed as shown in (C) by using the same dmSET used for the generation of randomizations (see Materials and Methods), but constraining each cell's position to its original location plus a small amount of noise. (**D**) Average cell neighboring areas versus cell areas in real tissues and in randomizations. (**E**) Cell positions (in µm) in the reconstruction of all replicates are nearly equal to segmentations. (**F**) Cell positions (in µm) in one randomization of all replicates are randomly shuffled. (**G**) Relative error on cell areas between the segmentation (or reconstruction) and the randomizations. (**H**) Cell areas (in µm$^2$) in reconstructions versus segmentation. (**I**) Cell areas (in µm$^2$) in randomizations versus segmentation. (G, H, I) show that cell areas are similar in segmentations, reconstructions and in the randomized tissues. (**J**) Relative error on cell lobeyness between the segmentation (or reconstruction) and the randomizations. (**K**) Cell lobeyness in reconstructions versus segmentation. (**L**) Cell lobeyness in randomizations versus segmentation. The color bar in (E) denoting cell areas is also associated with panels (F, H–I, K–L, N). (J, K, L) show that cell shapes are affected in a similar manner in reconstructions and original segmentations for cells larger than 10,000 µm$^2$. (**M**) Number of neighbors in function of cell areas in reconstructions and in segmentations are similar. (**N**) Number of neighbors in reconstructions versus segmentation. (**O**) Mean number of giant cell neighbors per giant cell (as shown in Fig. 5C) with the comparison with the reconstructed tissues "reconstructions". (M, N, O) show that the cell connectivity is affected in the reconstruction due to the shape artifacts, but that giant cell contacts are not significantly different and the results remain the same when comparing the reconstructions with the randomizations (see Materials and Methods). The six replicates were compared together with one corresponding randomized tissue in (D–N). ρ indicates Pearson correlation coefficient in (E–F, H–I, K–L, M). Two-sample t-tests were performed in (G, J, M) at each interval: $p < 0.05$ (*), $p < 0.01$ (**), $p < 0.001$ (***). The code and data associated with this figure can be found at Open Science Framework (osf.io), DOI:10.17605/OSF.IO/RFCWS.

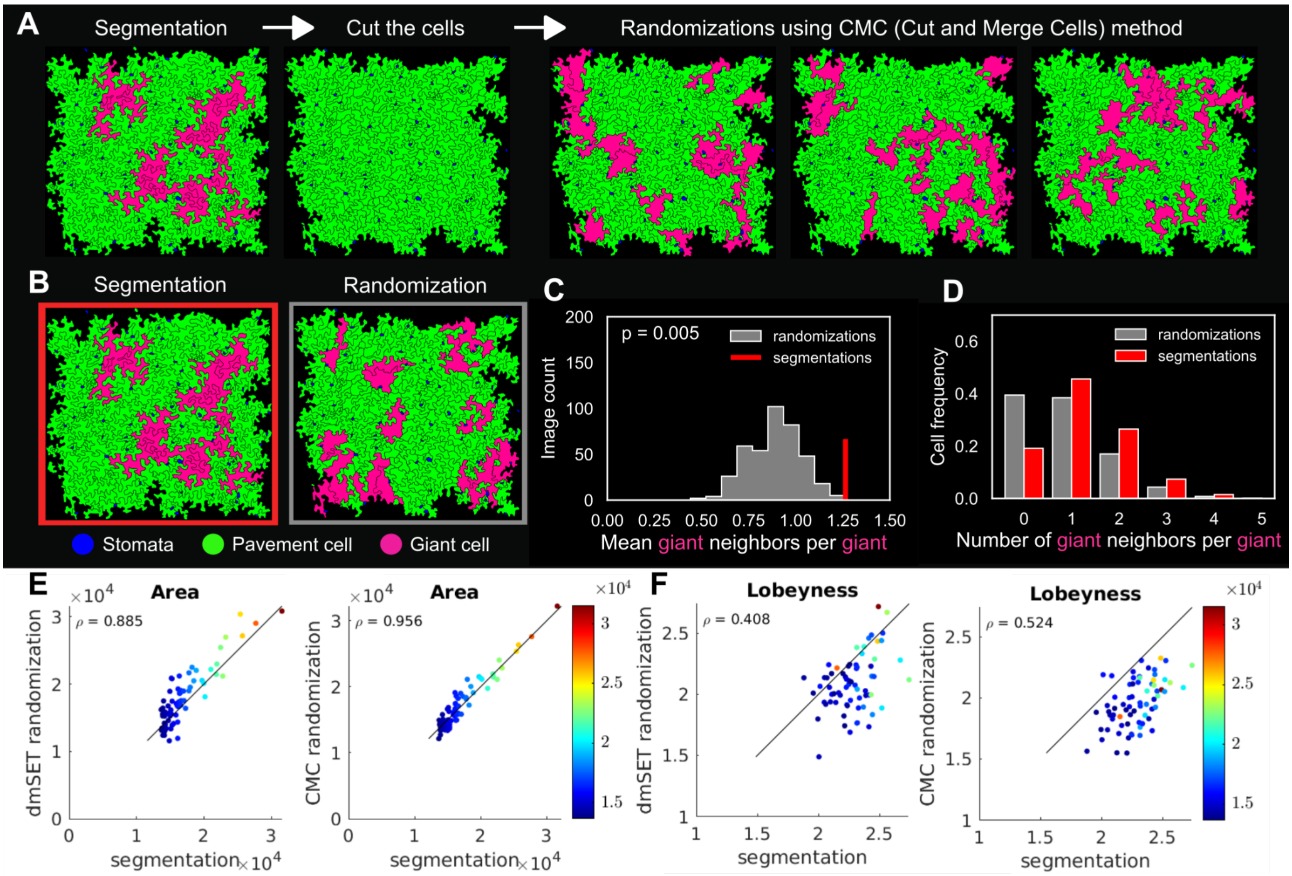

**Fig. S19. Giant cells are more clustered than in a randomized null model using the Cut and Merge Cells method.** (**A**) Overview of the alternative method referred to as Cut and Merge Cells (CMC) to generate random giant cell patterns (see Materials and Methods). Each leaf replicate was over-segmented to create templates made of small pieces of pavement cells. These templates were then used to automatically generate a random pattern of giant cells by preserving their sizes and numbers (see Materials and Methods). (**B**) Example of one initial segmentation of a 25-dpg leaf (left) and one corresponding CMC randomization (right). (**C–D**) Quantification of the giant cell patterns to compare with Fig. 5C–D. (C) Mean number of giant cell neighbors per giant cell in the real tissues (segmentation) and in its randomizations (in gray). (D) Distributions of the number of giant cell neighbors for all giant cells found in all replicates of segmentations (in red) and randomizations (in gray). (**E**) Left: Comparison of cell areas of giant cells in the dmSET randomizations and in the segmentations. Right: Comparison of cell areas of giant cells in the CMC randomizations and in the segmentations. (**F**) Left: Comparison of cell lobeyness (see Materials and Methods) of giant cells in the dmSET randomizations and in the segmentations. Right: Comparison of cell lobeyness of giant cells in the CMC randomizations and in the segmentations. Six replicates and one randomization per replicate were considered in (E–F). The color bars in (E–F) denote cell areas in $\mu m^2$. The code and data associated with this figure can be found at Open Science Framework (osf.io), DOI:10.17605/OSF.IO/RFCWS.

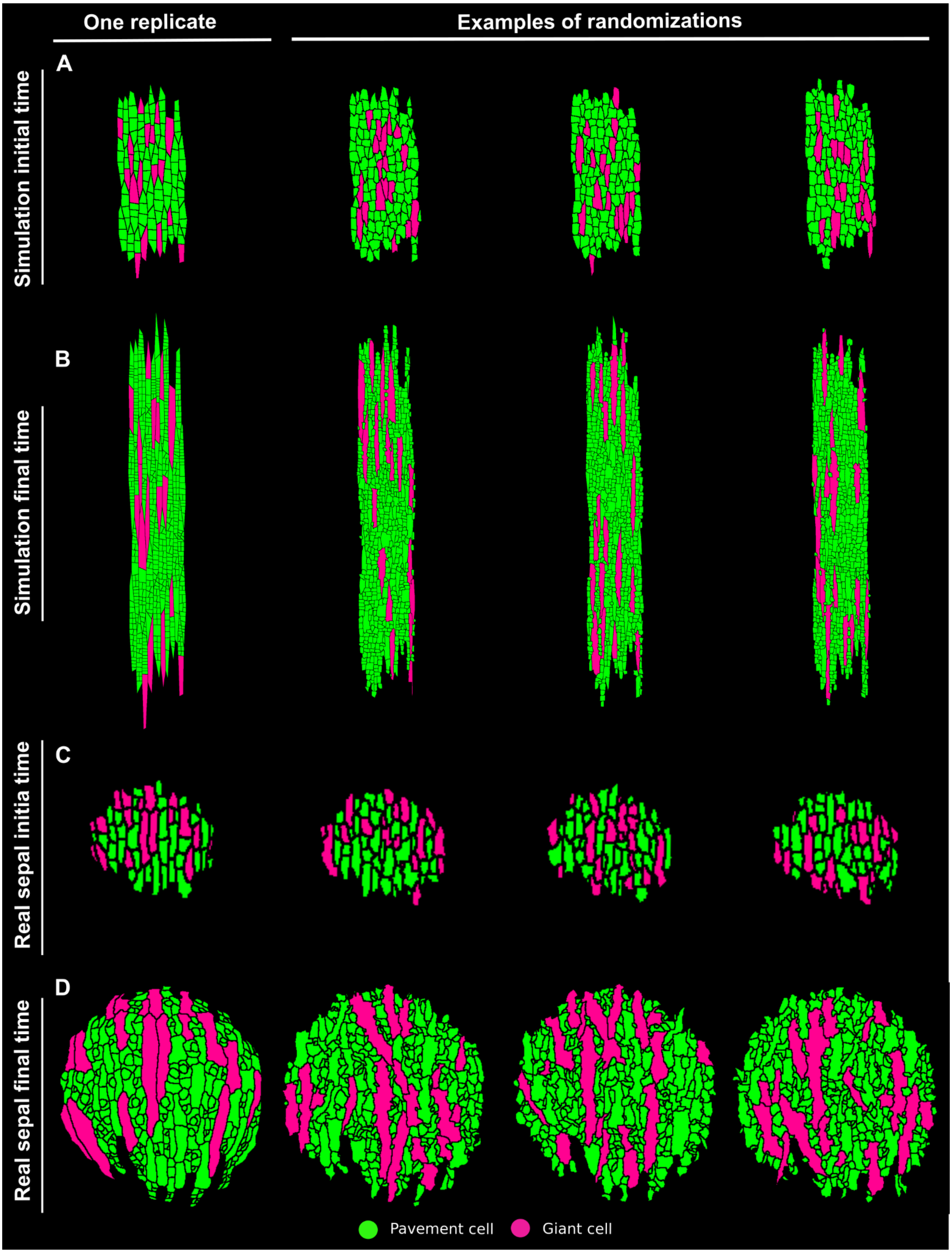


**Fig. S20. Examples of randomizations at initial and final time points in simulations and in time-lapse imaging sepal data.** One replicate is shown on the left and three corresponding randomized tissues on the right as an example. (**A–B**) Numerical simulation at time t = 55 (initial time) (A) and at time t = 135 (final time) (B). (**C–D**) Sepal from time-lapse imaging data at two different stages called here initial time (C) and final time (D). Associated with Fig. 7. The real sepals in (C) and (D) were also used for an independent analysis in Hervieux et al. 2016. The code and data associated with this figure can be found at Open Science Framework (osf.io), DOI:10.17605/OSF.IO/RFCWS.

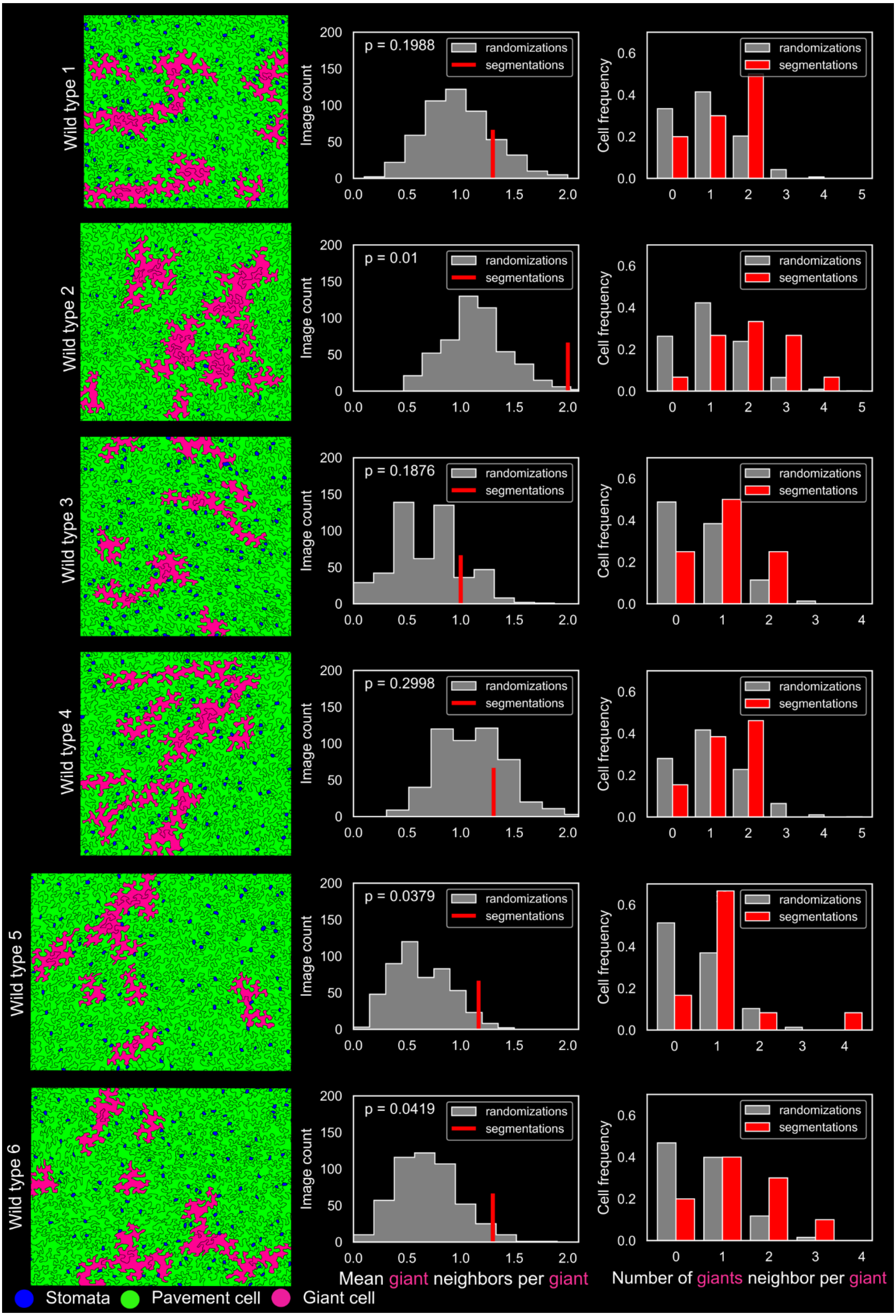
Wild type 1
p = 0.1988
Wild type 2
p = 0.01
Wild type 3
p = 0.1876
Wild type 4
p = 0.2998
Wild type 5
p = 0.0379
Wild type 6
p = 0.0419
randomizations
segmentations
Image count
Cell frequency
Mean giant neighbors per giant
Number of giants neighbor per giant
Stomata
Pavement cell
Giant cell

**Fig. S21. Quantification of the giant cell pattern in the individual replicates of the leaf.** On the left, each replicate is displayed after the cell type classification. In the middle: the corresponding results of the cellular pattern quantification showing the mean giant number of neighbors per giant, where the tissue segmentations (in red) were statistically compared with the tissue randomizations, containing 400 randomized tissues per replicate (in gray). On the right: the corresponding distributions of the number of giant cell neighbors per giant cell, both for the actual tissue (red) and the randomizations (gray). Associated with Fig. 5. The code and data associated with this figure can be found at Open Science Framework (osf.io), DOI:10.17605/OSF.IO/RFCWS.

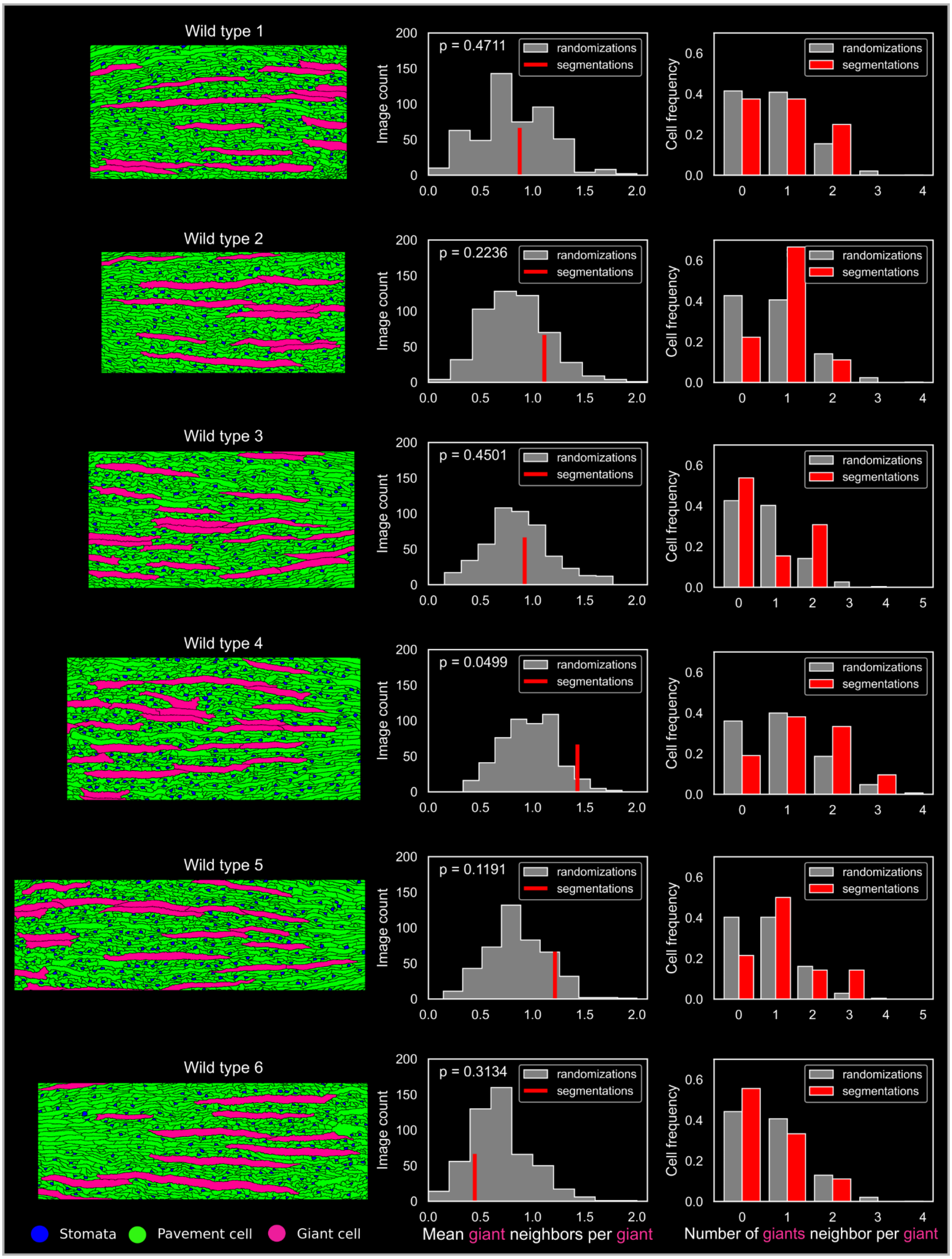


**Fig. S22. Quantification of the giant cell pattern in the individual replicates of the sepal.** On the left, each replicate is displayed (rotated 90 degrees) after the cell type classification. In the middle: the corresponding results of the cellular pattern quantification showing the mean giant number of neighbors per giant, where the tissue segmentations (in red) are statistically compared with the randomizations, containing 400 randomized tissues per replicate (in gray). On the right: the corresponding distributions of the number of giant cell neighbors per giant cell, both for the actual tissue (red) and the randomizations (gray). Associated with Fig. 5. The code and data associated with this figure can be found at Open Science Framework (osf.io), DOI:10.17605/OSF.IO/RFCWS.

### Supplementary Tables

| **Genotype** | Primer 1 (5′ to 3′) | Primer 2 (5′ to 3′) | Primer 3 (5′ to 3′) |
|---|---|---|---|
| *acr4-2* | TAGTCACTCTGTGGAATGTCTC (WT) GCACCTACAATTCCTCAATCTG (mut) | GCACCTACAATTCCTCAATCTG (WT) GCCTTTTCAGAAATGGATAAATAGCCT (mut) | |
| *atml1-3* | CAGGCAGAAGAAAATCGAGAT (WT) GAAACCAGTGTGGCTATTGTT (mut) | GAAACCAGTGTGGCTATTGTT (WT) TTGGGTGATGGTTCACGTAGTGGG (mut) | |
| *dek1-4* | TCCACAGGTAGTTTCTCTTGC | TGAAGACTGAAAGGACAAAAGGTGC | |
| *lgo-2* | CTTCTCAACCTCTCACTTCTCCAA | CCGAACACCAACAGATAATT | TTGGGTGATGGTTCACGTAGTGGG |
| *ATML1-OX* | GAAACCAGTGTGGCTATTGTT | CATATGGGAGACAGCTTTCTCATACGCG | |
| *LGO-OX* | CTTCCCTTTCTCCTAAGTTCCT | GATTATCTCACAAGTCGACAC | |

**Table S1. Primers for genotyping mutants and overexpression lines.**

| Parameter | Description | Values |
|---|---|---|
| $P_A$ | ATML1 basal production rate | 1.19 |
| $V_A$ | ATML1 auto-induction rate | 1.25 |
| $K_A$ | ATML1 concentration for half ATML1 auto-induction maximal rate | 1.9 |
| $n_A$ | Hill coefficient for ATML1 auto-induction | 5 |
| $G_A$ | ATML1 degradation rate | 1 |
| $V_T$ | Target maximal production rate | 10 |
| $K_T$ | ATML1 concentration for half ATML1-mediated target maximal production rate | 2.0 |
| $n_T$ | Hill coefficient for ATML1-mediated target induction | 1 |
| $G_T$ | Target degradation rate | 10 |
| $\Theta_T$ | Target threshold for inhibiting mitosis | 0.6 |
| $\Theta_{C,S}$ | Timer threshold for synthesis | 2 |
| $\Theta_{C,D}$ | Timer threshold for timer resetting | 3 |
| $P_C$ | Timer basal production rate | 0.1 |
| $E_0$ | Characteristic effective volume | 15 |
| | Exponential radial growth rate | 0.007 |
| | Exponential added growth rate to the vertical direction | 0.012 |

**Table S2. Parameter values used for the simulations.** Parameters used for the simulations shown in Fig. 7 and Fig. S20. All units are arbitrary.
